# The realised velocity of climate change reveals remarkable idiosyncrasy of species’ distributional shifts

**DOI:** 10.1101/823930

**Authors:** William D. Pearse, T. Jonathan Davies

## Abstract

To date, our understanding of how species have shifted in response to recent climate warming has been based on a few studies with a limited number of species. Here we present a comprehensive, global overview of species’ distributional responses to changing climate across a broad variety of taxa (animals, plants, and fungi). We characterise species’ responses using a metric that describes the realised velocity of climate change: how closely species’ responses have tracked changing climate through time. In contrast to existing ‘climate velocity’ metrics that have focused on space, we focus on species and index their responses to a null expectation of change in order to examine drivers of inter-specific variation. Here we show that species are tracking climate on average, but not sufficiently to keep up with the pace of climate change. Further, species responses are highly idiosyncratic, and thus highlight that projections assuming uniform responses may be misleading. This is in stark contrast to species’ present-day and historical climate niches, which show strong evidence of the imprint of evolutionary history and functional traits. Our analyses are a first step in exploring the vast wealth of empirical data on species’ historic responses to recent climate change.

---

The natural environment is changing rapidly. Over the last century, habitats have become increasingly fragmented and isolated^1^, the climate has become warmer, and extreme climate events more frequent^2^. Biological diversity—the species of plants and animals with which we cohabit the Earth—has responded predictably. Present-day extinction rates are estimated to be up to three orders of magnitude greater than background rates^3^ and are projected to increase further over the next several decades^4^. Current estimates suggest that over one million species may be threatened with extinction^5^, and that we have experienced a 50% decline in animal diversity in the past 40 years^6^.

The main direct human-induced drivers that impact biodiversity now are habitat loss^7^ and fragmentation^1^, but climate change is likely to become a dominant driver in the next few decades^8–10^. Projected impacts of climate change on biodiversity have attracted much attention, but the uncertainty around the magnitude of future extinctions (*e.g.*, 7.9%^10^ vs. 37%^11^ of species) highlights the inadequacy of current forecasts. In the face of changing climate species must either shift in space, to track favourable conditions, or time, for example, flowering and breeding earlier. Evidence suggests they are doing both^8,12–14^, but not all species are responding equally. Some species are expanding their distributions, sometimes invasively^15^, and some species are flowering later^16^; species may also be simply becoming less predictable^17^, perhaps as evolved responses to environmental cues break down. Simply quantifying species’ mean responses ignores the important variability in responses among species^18^. Developing robust predictions of ecological responses to climate change thus requires data from diverse species, ecosystems, and climates to account for this large inter-specific variation.

The velocity of temperature change—the rate a species would have to shift its range to maintain constant temperature—has been calculated at an average of 0.42 km/yr^19^. A meta-analysis of taxonomic groups estimated the average pole-ward shift in species’ distributions has been approximately 1.69 km/yr^14^, suggesting that species might be capable of keeping pace with shifting climate. Yet even this estimate is derived from just 21 studies with limited taxonomic coverage (361 birds, 9 mammals, and no plants), and critically any single point estimate does not reflect the large variation in responses among species or within taxa. Here we leverage the vast volume of distributional data stored in digital data repositories^20^, representing over 10,000 species of plants, fungi, mammals, birds, reptiles, amphibians, and insects, to provide the first global synthesis of species distributional responses to climate change. By contrasting historical niche dimensions with those of the present day, we can characterise the realised biotic velocity of climate change, which represents the relative degree to which species track climate across space.

We show that, on average, species’ historical (pre-1980) ranges have warmed significantly, and that species have generally shifted their distributions to ameliorate this warming, but not sufficiently to keep pace with climate change. Yet this general pattern masks a remarkably degree of variability in species responses, and many species’ ranges are now even warmer than if their distributions had remained static. Further, we find that, while species’ niche dimensions are strongly predicted by their evolutionary history and functional traits, the same is not true of their degree of tracking. Thus while species’ distributions are reasonably predictable, their changes in distributions are not.

## Results and discussion

### A comparative index of species’ niche tracking

We propose a measure that captures the extent to which a species’ distribution is tracking climate by standardising its observed niche change by the change it would have experienced had it not shifted its distribution. Formally, we define this as:

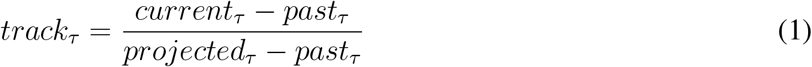

Where *current*_*τ*_ is a given quantile (*τ*) of a species’ climatic distribution across its present-day range, *past*_*τ*_ the same quantile of a species’ range in the past, and *projected*_*τ*_ the same quantile of a species’ past range under present-day climatic conditions. This index should be calculated separately for temperature, pollution, or whatever environmental variable is of interest. Thus a *track* of 0 would indicate a species that has tracked changing climate perfectly, while a *track* of 1 would be consistent with a species that had not moved whatsoever to track changing environmental conditions. Values between 0 and 1 suggest moderate tracking, while values greater than 1 or less than 0 represent a change in climate relationship greater than if the species had not moved at all. Uncertainty in either species’ past or current ranges can be accounted for through a bootstrap procedure, whereby the past and current distributions are re-sampled to standardise sampling across time periods (see methods).

Our approach has two main innovations: its focus on (1) the whole of a species’ distribution and (2) expected change. (1) When estimating species’ range distributions we typically focus on species’ limits^21^ (*e.g.*, upper and lower quantiles) while in community ecological^22^ and macro-evolutionary^23^ studies we tend to focus on central tendencies (*e.g.*, the median). However, there is no strong *a priori* reason that species can, or must, track all quantiles equally. Indeed some evidence suggests different constraints on species’ warm-range limits versus their cool-range limits, with biotic interactions more influential in the former, and climate determining the latter^24^. It is possible, therefore, that the drivers of species’ range edges and centres may be diverging under climate change. Our approach is to check all quantiles of a species’ distribution. (2) The magnitude of experienced and projected climate change shows substantial geographical variation^19^, and so species’ responses might thus be expected to also vary in space. We suggest that species’ distributional responses should be measured on the basis of the magnitude of change to which they are exposed, and the denominator of our index re-scales species’ change around an expectation under distributional stasis. This change reflects the magnitude of ecological and evolutionary selection, and while there are many ways of addressing this, we suggest ours is perhaps the simplest and most transparent, so represents a fair starting point. We note that we could also rescale our index according to the distance that would need to be travelled to maintain climate stasis (*sensu* Loarie *et al.*^19^), but prefer to present here a unitless index of tracking, which is simpler to compare across taxa and between biogeographic regions.

### Niche tracking across the tree of life throughout the past sixty years

To assess the extent to which species have tracked climate, we used the most comprehensively sampled time-series data for species’ distributions (GBIF^20^) and climate (UEA’s CRU^25^). This represents over 630 million raw occurrence records collected over the past century. We focus on five quantiles: the 5th, 25th, 50th (median), 75th, and 95th. Here, we present species’ range data collected from 1955–2015 for 10,700 species (4,879 plants, 1,517 fungi, 273 mammals, 433 birds, 147 reptiles and amphibians, and 3,451 insects), each observation matched to one of nine climate variables in the year in which they were sampled (mean, minimum, maximum temperature, precipitation, ‘rainy-day’ frequency, vapour pressure, potential evapotranspiration, ‘frost day’ frequency, and cloud cover). In the methods we give more detail of our reproducible analysis pipeline, and in the supplementary materials provide instructions for repeating our analyses with different data curation and cleaning choices. Our metric was designed to account for known biases in the GBIF data^26^, and in the methods we outline our simulations which address the influence of sampling uncertainty.

Almost all species are tracking some climate variable through time (99.5% have at least one *track*_*τ*_ between 0 and 1). Most species are, to some extent, tracking temperature (see Figure 1); 65% track at least one quantile and 43% track at least two. The three temperature indices are, however, the least-tracked of any climate variable, while precipitation is the most-tracked (86% of species track at least one of its quantiles). Notably, however, many taxa are not tracking climate through time, and many have experienced more change than if their distributions had remained static. Just as most species are tracking at least one quantile of one climate variable, 99.9% of species have ‘overshoot’ (have a *track*_*τ*_ less than 0; for example becoming colder in a warming region) in at least one quantile of at least one climate variable. As we show in Figure 1, these results are consistent with our null simulations where species’ range change is a combination of essentially random movement and a moderate degree of subsequent climate filtering. Thus while our results confirm that, by-and-large, species are tracking climate change, the magnitude of this varies across species, taxa, and aspects of climate (see Figure 1). For example, while fungi and insects are, on the whole, broadly tracking the median of temperature (median *track*_50%_ of 0.46 and 0.64, respectively), plants and mammals track temperature (median *track*_50%_ of 1.16 and 1.07, respectively) more poorly than precipitation (median *track*_5%_ of 0.79 and 0.89, respectively).

**Figure 1:**
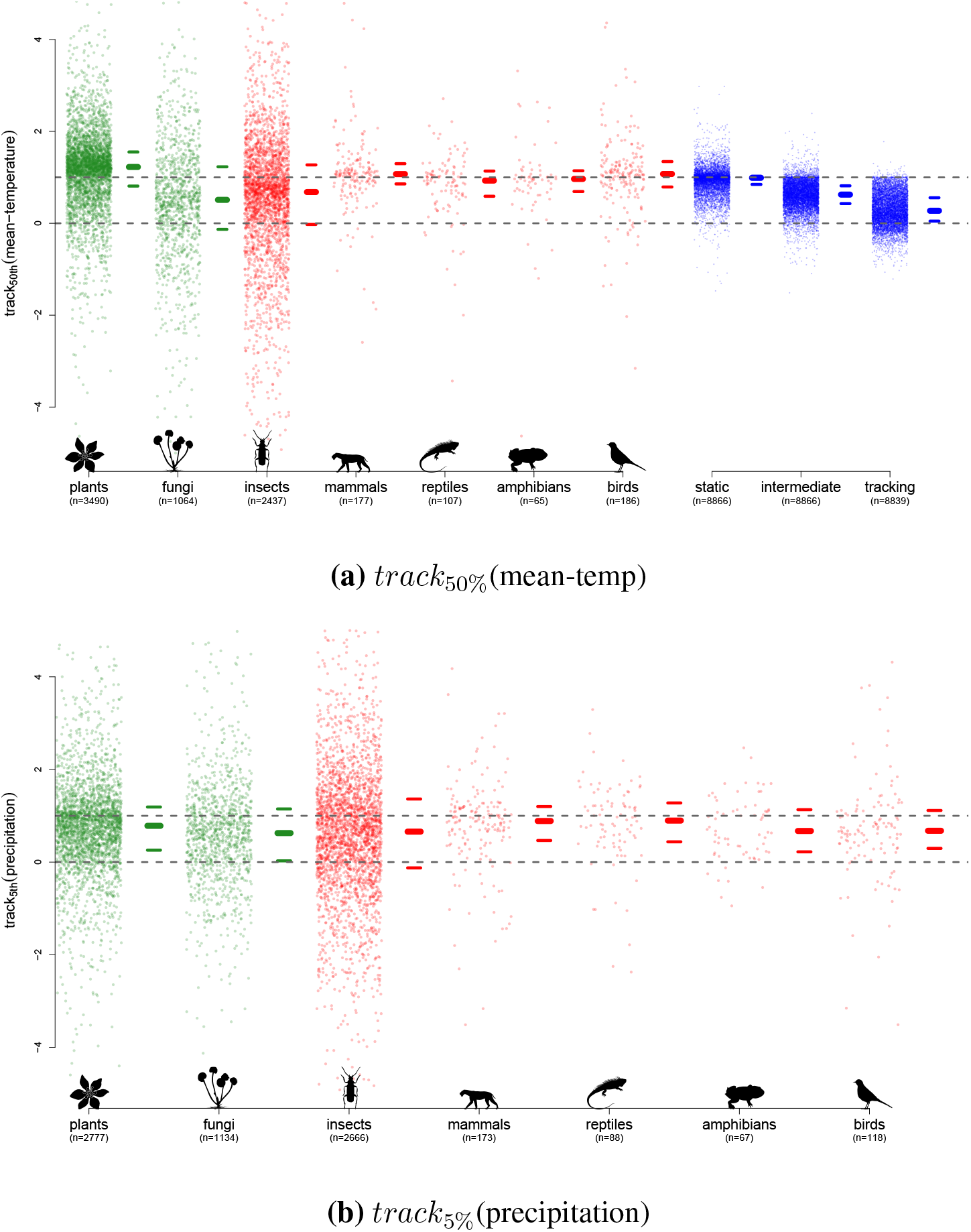
Most species are tracking some aspect of climate, but there is pronounced variation among species and clades. In (a) we show the 50th quantile (median) of our tracking index of mean annual temperature, and in (b) the 5th quantile of our index of precipitation. On the left of each plot, our seven major taxonomic groups plotted (in green for plants and fungi, and in red for animals). At the far-right of (b), we show the output from our simulations of species that are perfectly tracking, imperfectly tracking, or not tracking climate at all (in blue; see Methods for details). To the right of each distribution of points the median (thickest line) and inter-quartile range (thinner lines) are shown as horizontal lines for each distribution. While the majority of species are tracking global climate to some extent, there is is profound variation among species and clades. Figure 1a makes clear the variation among and within taxonomic groups; the distributions of fungi, insects, and reptiles are, broadly, tracking changes in mean temperature well, while other taxa (*e.g.*, mammals and plants) are not. Comparing Figures 1a and 1b makes clear how different taxonomic groups and species may be tracking different aspects of climate; plants are broadly tracking the lower-limit of precipitation, for instance.

### Idiosyncratic drivers of species’ tracking

To examine the potential drivers of species’ ranges and range tracking, we first examined the phylogenetic signal of species’ climate quantiles. We focused on mammals, birds, reptiles, and plants, since we were able to find broadly inclusive, dated phylogenies for these taxa. We found strong evidence of phylogenetic signal in species’ past (median Pagel’s *λ*^27^ of 0.60) and current (median *λ* of 0.57) distributions, but limited signal in our index (median *λ* of *<*0.001). Past climate showed stronger phylogenetic signal than our index in 158 of all 180 (4 taxa, 5 quantiles, and 9 climate variables) possible taxon-climate-quantile combinations (144 for current climate; see Figures 2 and 3). This confirms the known result that species’ current-day and past ranges are strongly influenced by evolution^28^, but suggests that species’ realised velocity of change (our tracking index) may be associated with different factors.

**Figure 2:**
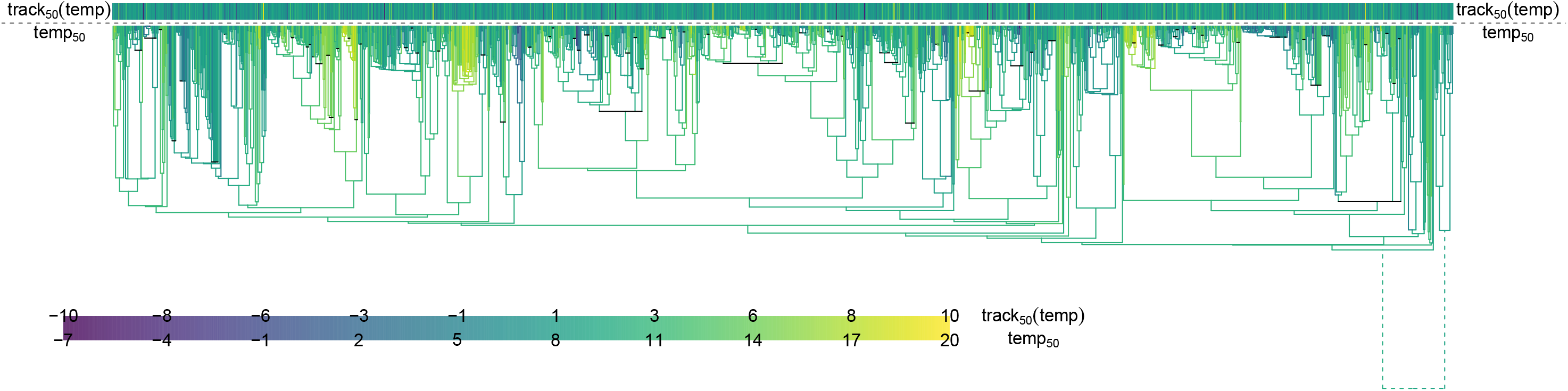
Despite strong phylogenetic signal in species’ static distributions, species’ changes in distributions are not phylogenetically patterned. Here we show the plant tree of life, with the two oldest branches shortened for compactness (emphasised with dashed lines). Internal branches are shaded by ancestral state reconstructions, assuming Brownian motion, of species’ median mean-temperature in the past. Phylogeny tips are coloured according to the track index of the same underlying data (compare with Figure 1). This general pattern of strong phylogenetic signal of climate (shown by the coloured pattern to the internal branches), but not of degree of tracking, is characteristic of almost all of our underlying climate data (see also Figure 3).

**Figure 3:**
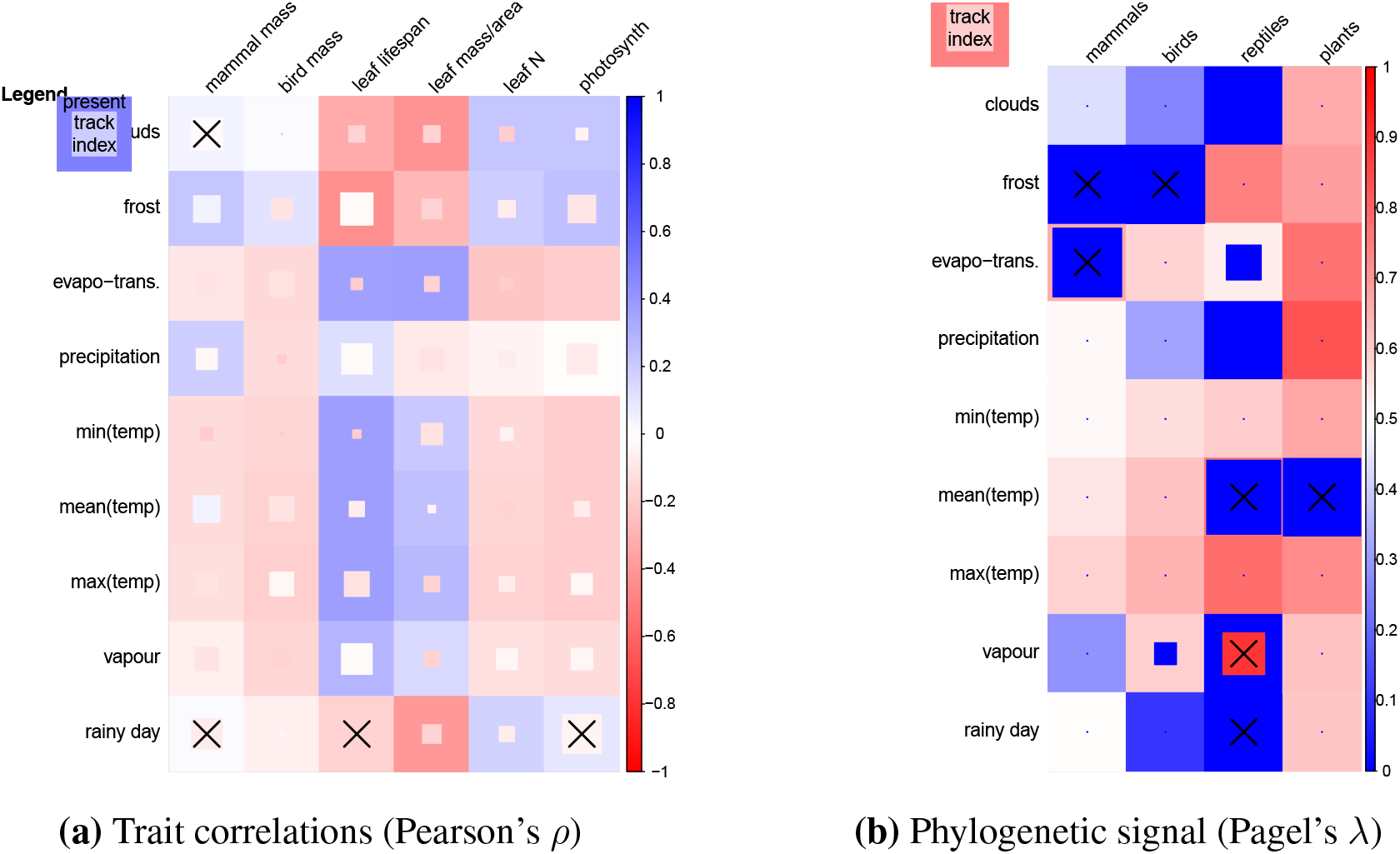
Species’ static climate variables show strong trait correlations and phylogenetic signal, but their relative degree of tracking does not. In (a), we show the correlation (Pearson’s *ρ*) between the median of each climate variable (rows) with each of our included traits (columns) using colour (see legend at the right). In each cell, the outer, larger box depicts the correlation between species’ present-day climate and the trait, while the smaller, inner box the correlation for species’ relative tracking (whose size is proportional to intensity of correlation). The 4 (of 54) track-index correlations that are statistically significant (at *α*_5%_) are highlighted with crosses. In (b), we show, for each taxon, the phylogenetic signal (Pagel’s *λ*^27^) of major climate variables and relative tracking indices (rows) using colour (see legend at the right). As with (a), in each cell the outer, larger box depicts the correlation between species’ present-day climate and the trait, while the smaller, inner box the correlation for species’ relative tracking (whose size is proportional to phylogenetic signal strength). The single track-index metric for which phylogenetic signal was stronger than the underlying data is highlighted with a cross.

We additionally examined species’ traits as potential drivers of species’ distributions and potential to track changing environment. We focused on mammals, birds, and plants, since global, openly-released trait datasets exist for these taxa. Again, we found statistically significant associations between traits and past distributions of 191 of 270 (at *α*_5%_; 6 traits, 9 climate variables, and 5 quantiles) taxon-climate-quantile combinations (193 for current distributions), but not between the traits and our tracking index (13 combinations or 5%; see Figure 3). While the traits we used are strongly associated with species’ climate responses^29,30^ (hence their strong predictive power of species’ distributions, but not degree of tracking), we emphasise that we could not find sufficient data to assess the potential impact of dispersal ability^31^. These findings, while surprising, are in keeping with emerging evidence that species’ traits are a relatively poor predictor of change at range edges^32^.

### Species’ movement through climate space is disconnected from their position within it

Given our index is poorly predicted by traditionally important variables, we now consider whether index values are predictive of each other. It is well known that the axes of global climate are not independent and that they are not changing independently^2,25^, and so we would expect species’ relative tracking of climate to show similar patterns. As we show in Figure 4, principal components analysis of species’ current and past climate distributions, which we refer to as their climate space, shows strong correlations across climate variables. Yet our index does not; species’ relative degree of tracking is both of much higher dimension (the amount of variance explained by each principal component axis is similar) and shows relatively less correlation across climates axes. We argue, therefore, that while species’ positions in climate space show strong associations, species’ relative tracking of their position in climate space does not. This result explains the poor predictive power of species’ traits and evolutionary history for our index: the ecological and evolutionary rules that determine species’ climate relationships are changing. Thus, in our analysis, we can explain species’ current distributions only by the degree to which they resemble past distributions.

**Figure 4:**
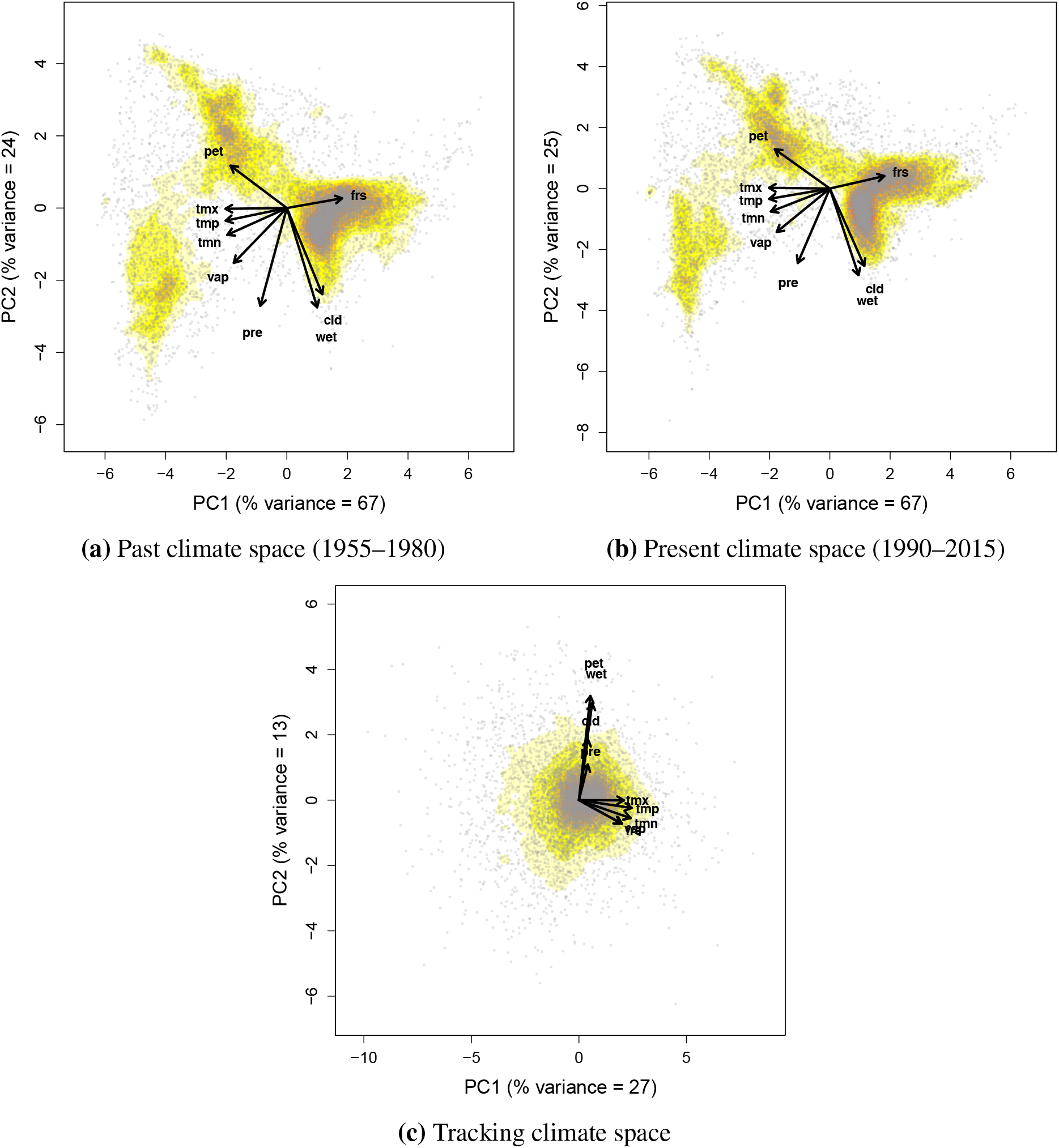
Species’ movement through climate space differs from their current and past positions within it. Principal component analysis (PCA) biplots of (a) species’ median past climate values, (b) median current climate, and (c) the median track-index of all climate variables. Each PCA axis explains some amount of variation in the underlying data (labelled on the axes themselves). In (a) and (b), species’ climate variables are correlated with one-another, and the space reflects global patterns in climate. In (c), this pattern has broken down: while some associations among climate axes still remain (notably temperature), the general pattern is one of uncorrelated shifts through climate space.

There are alternative explanations for our results, of which we highlight three: (1) we are ignoring important niche axes, (2) we are ignoring biotic interactions, and (3) our index highlights random variation in distribution. (1) Our climate variables are not all-encompassing, and we ignore other drivers such as geology and habitat-type. Yet we do detect strong pattern in current and past distributions, suggesting that the absence of pattern in our indices is meaningful. (2) Competition and facilitation among species are likely strong drivers of species’ ability to track climate^33^. Yet, again, they are surely also likely to have driven species’ past distributions, and so it is unclear to us why only our metric, and not also current and past distributions, is so poorly predicted by our trait and phylogenetic data. (3) If species were perfectly tracking climate, *current*_*τ*_ would be equal to *past*_*τ*_ and so our index values could reflect sampling error or some other purely stochastic input. Yet such noise, if truly random, would mean our empirical index values would be centred at 0 or resemble our simulated data, neither or which is the case (see Figure 1).

## Conclusion

Our results are consistent with the processes underlying species’ responses to climate change being distinct from those that generated their distributions in the past. This is in keeping with emerging evidence from the paleo-ecological literature that species’ historic ranges were more variable than previously thought, and seemingly influenced by different environmental factors than today^34,35^. During and after the last ice-age, for example, ‘no-analogue’ conditions generated previously-unseen species assemblages^36^. Equally, our observation of idiosyncrasy in species’ degree of tracking is consistent with species’ population crashes (and so extinctions) becoming less predictable during previous mass extinction events^37,38^. We call upon others to examine the additional drivers of our metric of relative change, and to extend its definition to include more nuanced definitions of climate space. We predict, however, that if the current mass extinction event continues, it is likely that patterns of idiosyncrasy among species’ declines and distributional changes will become more common.

## Acknowledgements

WDP was funded by NSF ABI-1759965, NSF EF-1802605, and USDA Forest Service agreement 18-CS-11046000-041. TJD was funded by *Fonds de Recherche Nature et Technologies* grant number 168004. The images in Figure 1 are taken, with gratitude, from http://phylopic.org/; all are under the Public Domain Dedication 1.0 license and uncredited unless otherwise stated below. The reptile image was drawn by Jack Mayer Wood, the plant by Jason McNair, the insect by Birgit Lang, the mammal by Scott Hartman (used under the Creative Commons Attribution 3.0 Unported license), and the amphibian by Nobu Tamura (used under the Creative Commons Attribution 3.0 Unported license).

## Author Contributions

WDP and TJD contributed to all aspects of the study.

## Author information

The authors declare no competing financial interests. Correspondence and requests for materials should be addressed to

## Methods

All analyses were conducted in *R* version *3.6.1*^40^, and all names in *italics* refer to *R* packages. Code to reproduce all analyses in their entirety is available in the online supplementary materials and online at https://github.com/willpearse/track-index. In the supplementary materials, we provide extended figures summarising all tracking index quantiles and climate variables in the same detail as presented in the main text.

### Data collation

Species’ distribution data were downloaded from the Global Biodiversity Information Facility (GBIF)^20,41^. We downloaded only data highlighted by GBIF as not having any spatial issues, meaning they contained no obvious location errors (*e.g.*, latitude/longitude swaps or rounded coordinates). These occurrence data include records that were flagged as having potential issues and corrected by GBIF; our results were qualitatively identical if we excluded these records and so we include them here for completeness. We used ‘human observation’ occurrences (*e.g.*, records from surveys), and not vouchered specimens (*e.g.*, pressed specimens in a herbarium) because preliminary analysis revealed qualitatively identical results across the two methods, but with many fewer observations vouchered observations.

We only worked with species that had at least 1000 records on GBIF within our focal time periods (1955– 1980 and 1985–2015). We focused on these time periods because they contain sufficient observations for large-scale analysis. Since climate change was underway within the 1980s^2^, this also allows us to have reasonably large (if naïvely defined) ‘pre-change’ and ‘post-change’ periods. Our 1000 observation threshold limits us to better-studied species, but our results are robust to increases (*e.g.*, 10,000 observations) in this threshold. We only made use of data for which species identity was known, and we ignored variation among sub-species. The GBIF taxonomy is, itself, to some extent checked and harmonised, and so we retained in our analysis any species within GBIF that was not present in another datasets (see Cornwell *et al.*^42^ for a discussion of the GBIF taxonomy for plants). In the supplementary materials, we give instructions for re-running our analysis with differing curation thresholds to verify our choices.

Climate data were downloaded from Harris *et al.*^25^’s nine global time-series. We split the data into yearly (averaged) 0.5° cells, from which we estimated each of our nine climate variables for each species observation using *raster*^43^. Plant trait data were taken from Wright *et al.*^29^ (leaf lifespan, leaf mass-per-unit-area, leaf Nitrogen mass, and leaf photosynthetic capacity), mammal body mass data from the Amniote Trait Database^44^, and bird body mass data from EltonTraits^45^. We were unable to find sufficient coverage of the other taxa in this study in open-access trait databases to facilitate further analysis. We used existing global mammal^46^, bird^47^, amphibian^48^, plant (‘ALLOTB’ phylogeny^39^), and reptiles^49^ phylogenies, and in cases where posterior distributions of trees were available we used a single draw from that distribution. Missing species were added into the phylogenies using *congeneric.merge* in *pez*^50^.

### Calculating and testing our track index

As we describe in the main text, we quantify the realised velocity of climate change using a simple index of climate tracking. Our index, *track*_*τ*_, scales the observed magnitude of species’ shift in climate (*current*_*τ*_ − *past*_*τ*_) according to the degree of change to which that species was exposed (*projected*_*τ*_ − *past*_*τ*_). Our metric is comparable across species, and can also be calculated in such a way as to control for changes in sampling. The sampling of species through time and across space is known to be uneven across the data in GBIF^26,51^. To ensure our method was statistically robust to such changes, we employed a bootstrap procedure during the calculation of *track*_*τ*_. To do this we randomly re-sampled, with replacement, the occurrences making up *current*_*τ*_ and *projected*_*τ*_ to be of the same number as *past*_*τ*_, and vice-versa for *past*_*τ*_ and *current*_*τ*_. We repeated this process 999 times, and calculated a *track*_*τ*_ value for each re-sample, generating 999 *track*_*τ*_ values whose medians we report here. This process accounts for uneven sampling by generating a pseudo-posterior distribution of conservatively-estimated values. In each bootstrap, the better-sampled time period is sub-sampled to match the poorer-sampled period, accounting for differences in sampling. Equally, the poorer-sampled period is re-sampled in order to increase the variance around the index, essentially cross-validating our estimate. When applied to empirical data, our bootstrap approach reveals that the certainty differing tracking quantiles (and climate variables) is uneven. We therefore excluded uncertain (*i.e.*, estimates with high variation across pseudo-posteriors) and outliers from figures within the main text, and so report sample sizes within Figure 1 (and its counterparts in the supplementary materials).

To assess the performance of our (bootstrapped) track index, we simulated species’ with ranges that tracked climate to varying extents through time. Using the climate data from above in the years 1962 and 2002 (the mid-points of the ranges of our data), we simulated species with varying maximum possible range sizes (2×2, 5×5, 10×10, or 20×20 grid-cell extents) and varying latitudinal shifts in range through time (−4, −3, …, 3, or 4 grid-cells). We also varied species’ degree of environmental tracking (*α*; taking a value of 0, 0.5, or 1) and probability of occupancy(*σ*; taking a value of 0.5,. 75, or 1). Together, these latter two parameters define a species’ probability of being present in a particular cell within its range: 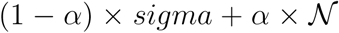. 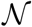 is defined as a scaled Normal probability density function with a mean equal to the median of the species’ distribution in 1962 and a variance of 1. Thus a species with an *α* of 1 is present only in cells within its range that resemble its climatic centre (*i.e.*, are similar to the median value of the species’ climatic distribution), and a species with an *α* of 0 is present in proportion *σ* of the cells within its potential range. We simulated all possible combinations of parameters across 100 possible centre-points of species’ ranges, providing a full range of potential range shifts and 26,598 total simulation runs. As with our empirical data, in simulation runs where, by chance, simulated species were not present in the past or current time-period, they were excluded from the analysis.

### Trait, phylogenetic, and PCA analyses

We calculated the Pearson’s correlation between all estimated climatic niche variables (*current*_*τ*_, *past*_*τ*_, *projected*_*τ*_, *track*_*τ*_ and the bootstrapped *track*_*τ*_). While we show only a handful of these correlations in Figure 3, we provide all correlations in the supplementary figures (and note that the qualitative results of all correlations are the same). For the phylogenetic signal analyses, we likewise calculated Pagel’s *λ* (using *phytools*^52^) for all indices, reporting all results in full in the supplementary materials (results are, again, qualitatively identical to the results in the main text). Principal component analyses were performed on all current, past, and track indices across all variables for a given quantile (*i.e.*, the 5th, 25th, 50th, 75th, and 95th). While we report only the 50th quantiles in the main text, in the supplementary materials we give results for all quantiles, which are qualitatively identical to the results in the main text.

