## supplemental figures for "The realised velocity of climate change reveals remarkable idiosyncrasy of species’ distributional shifts"

Our supplementary materials are organised according to the figures in the main text. We replicate the figures (and so analyses) calculated for the subsets of the data shown there across the entire range of the data below. Figure legend are terse and perfunctory because the interpretation of them can be done with reference to the figures in the main text (which are references in the legends of the supplementary figures).

### 1 Overall plots of tracking index values (*c.f.* figure 1)

#### 1.1 Cloud cover

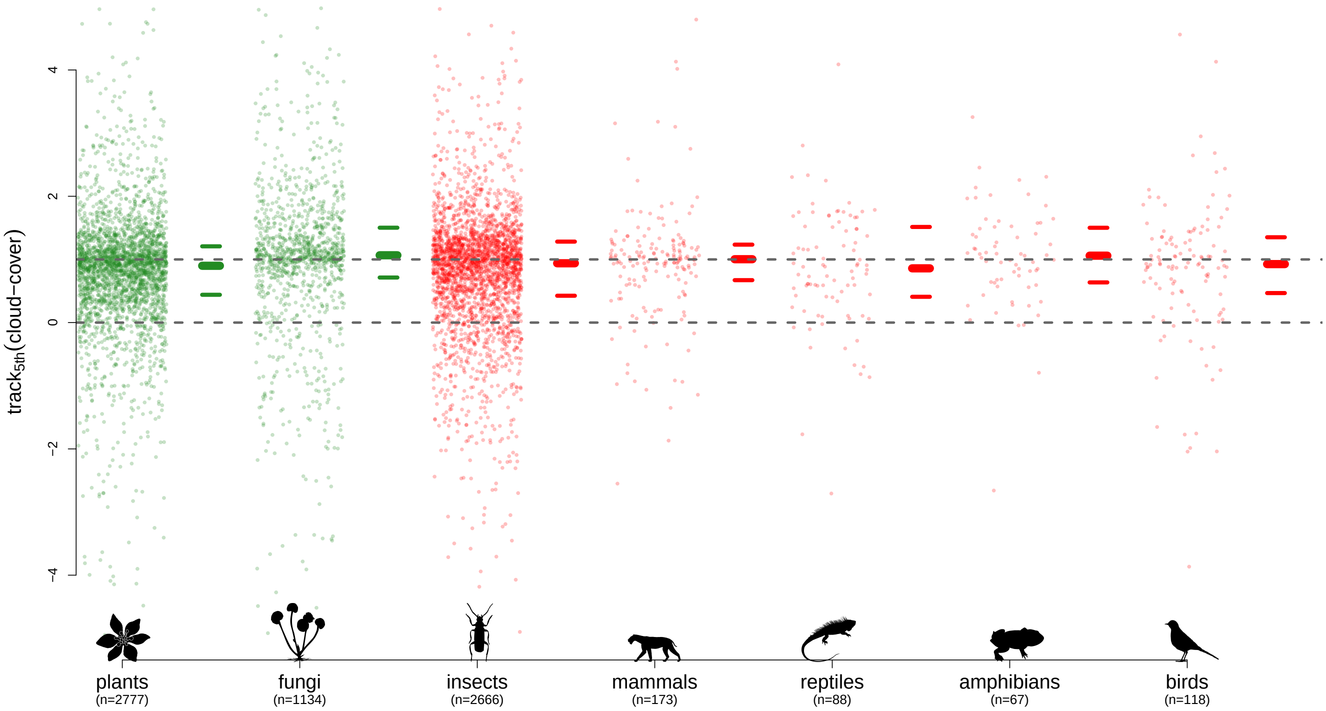


**Figure 1: Cloud cover, 5th quantile.** Legend as figure 1 in main text.

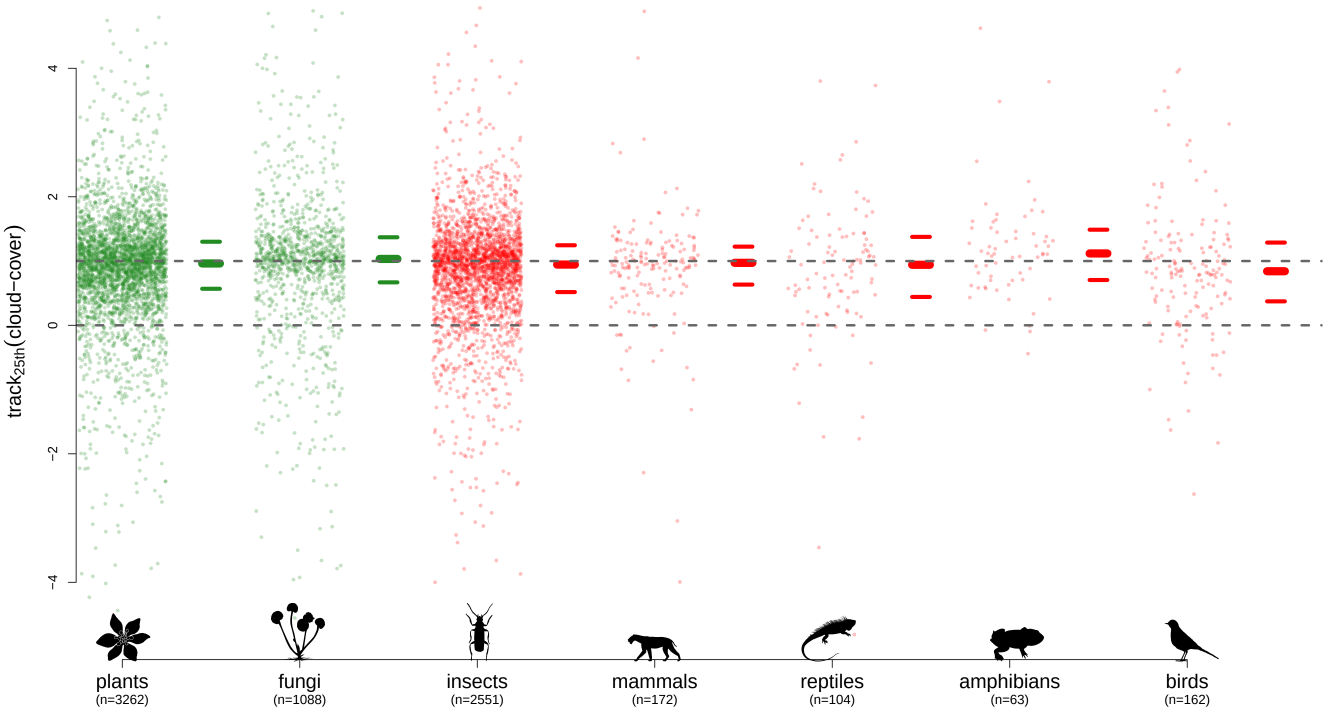


**Figure 2: Cloud cover, 25th quantile.** Legend as figure 1 in main text.

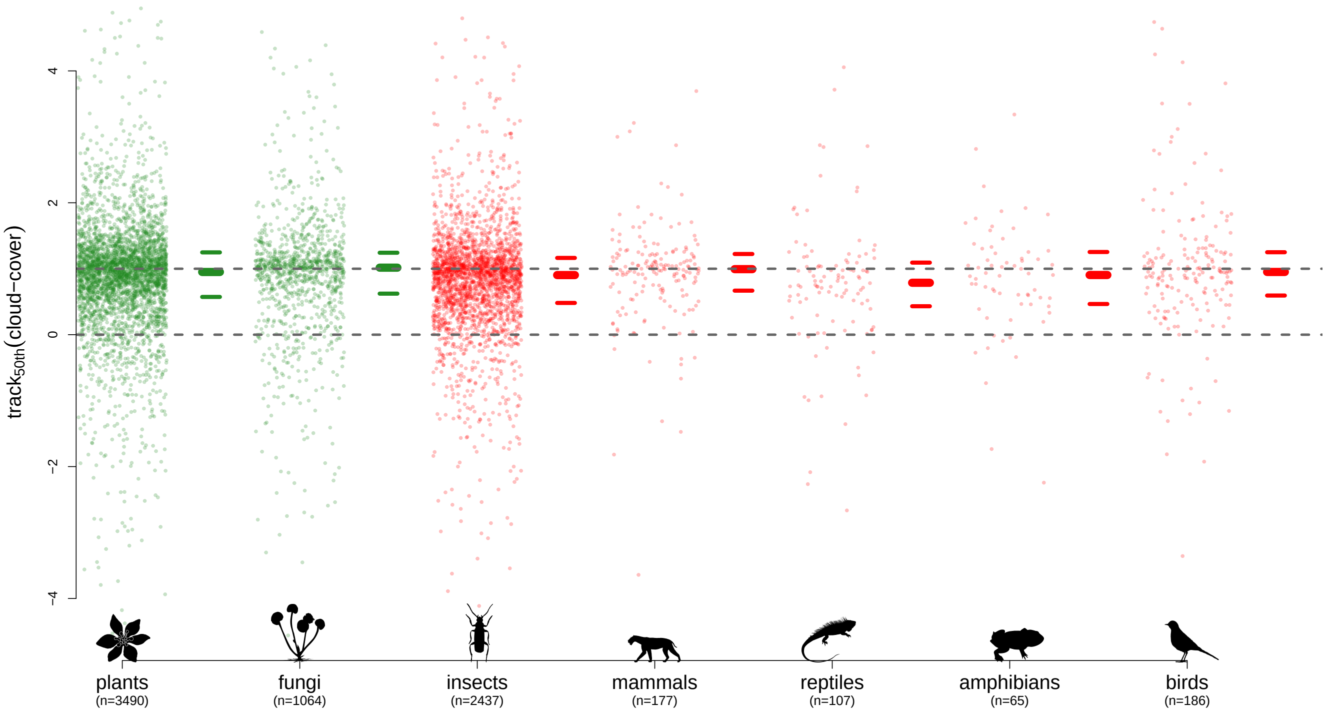


**Figure 3: Cloud cover, 50th quantile (median).** Legend as figure 1 in main text.

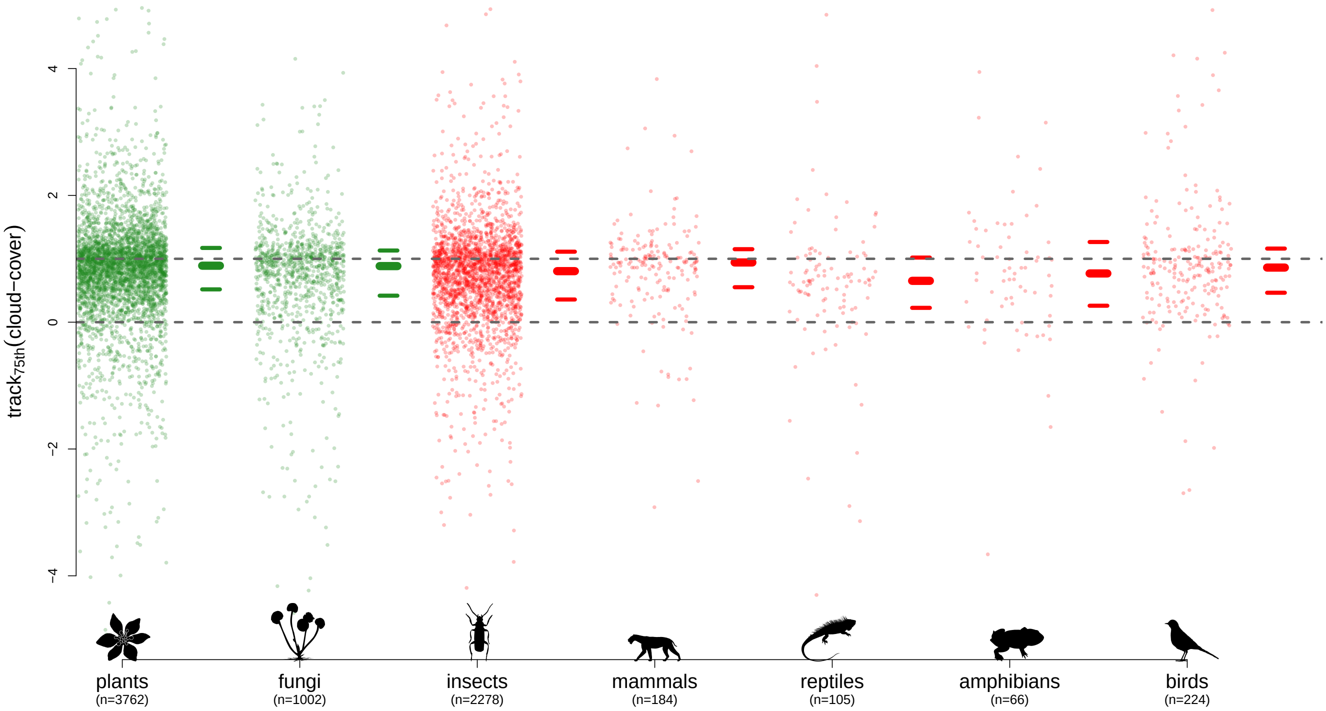


**Figure 4: Cloud cover, 75th quantile.** Legend as figure 1 in main text.

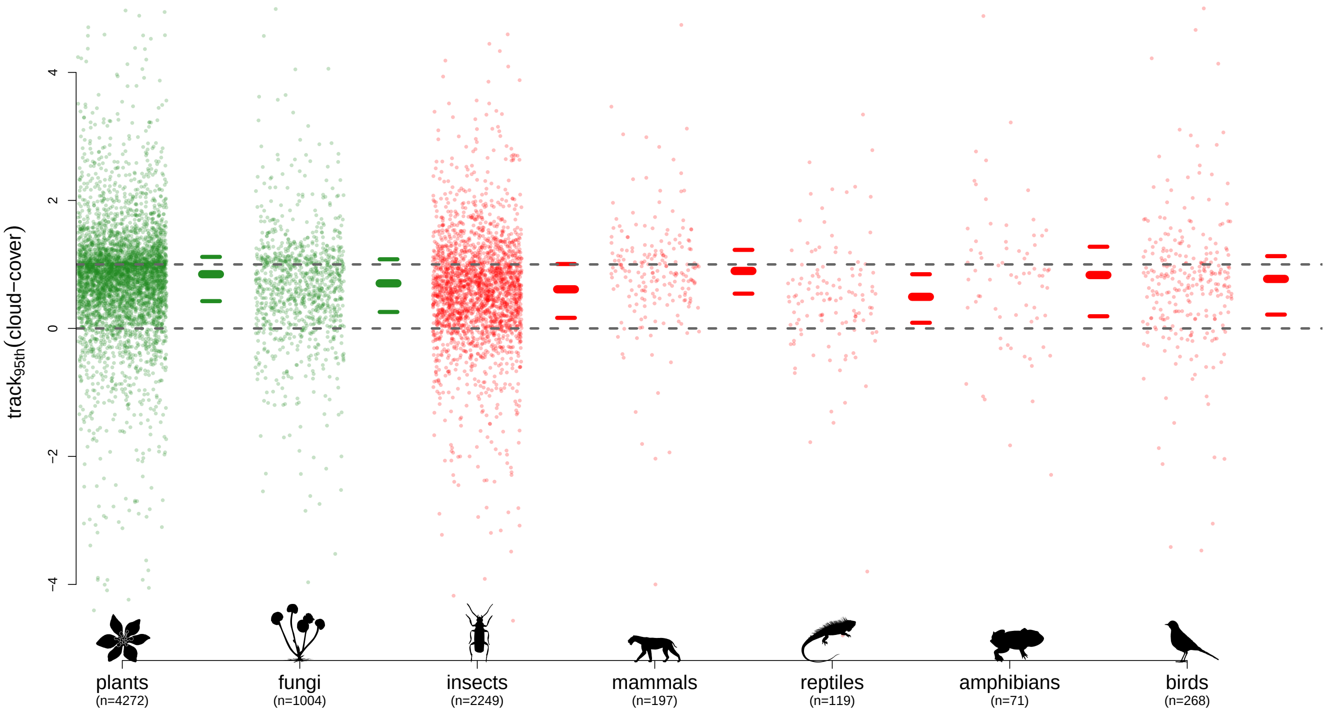


**Figure 5: Cloud cover, 95th quantile.** Legend as figure 1 in main text.

#### 1.2 Frost days

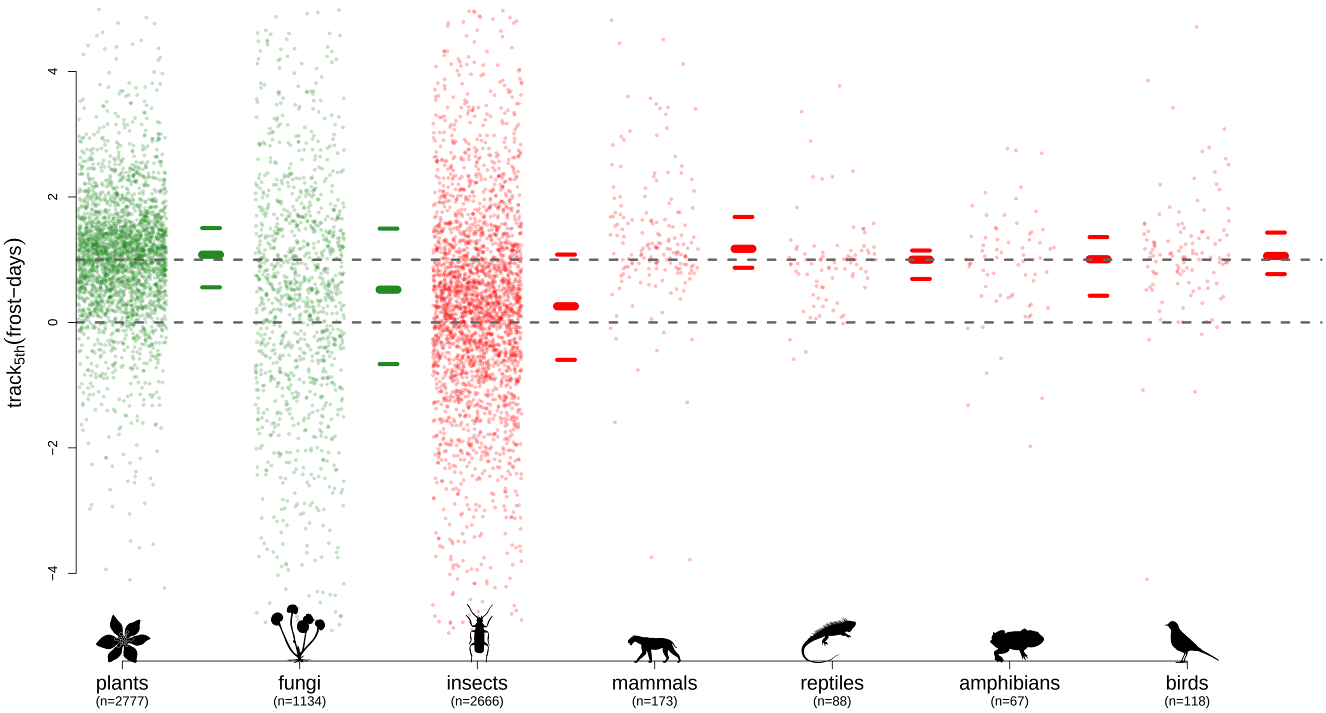


**Figure 6: Frost days, 5th quantile.** Legend as figure 1 in main text.

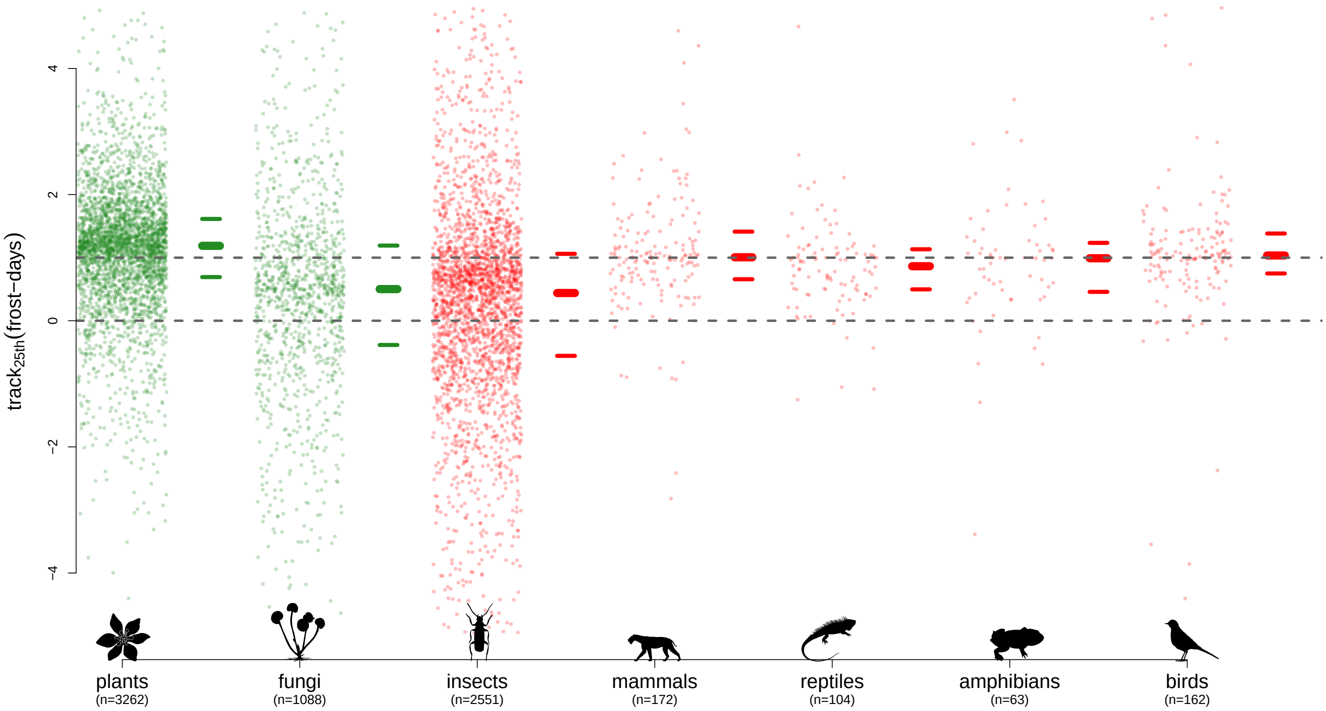


**Figure 7: Frost days, 25th quantile.** Legend as figure 1 in main text.

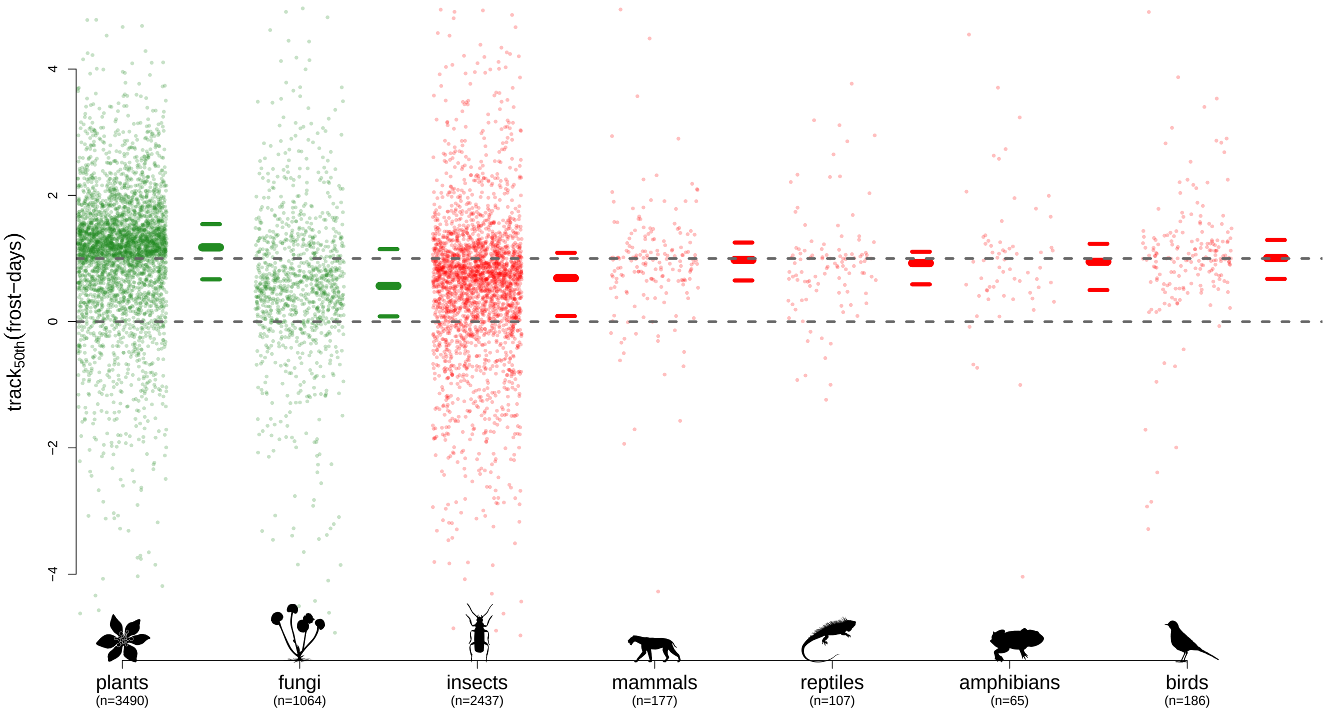


**Figure 8: Frost days, 50th quantile (median).** Legend as figure 1 in main text.

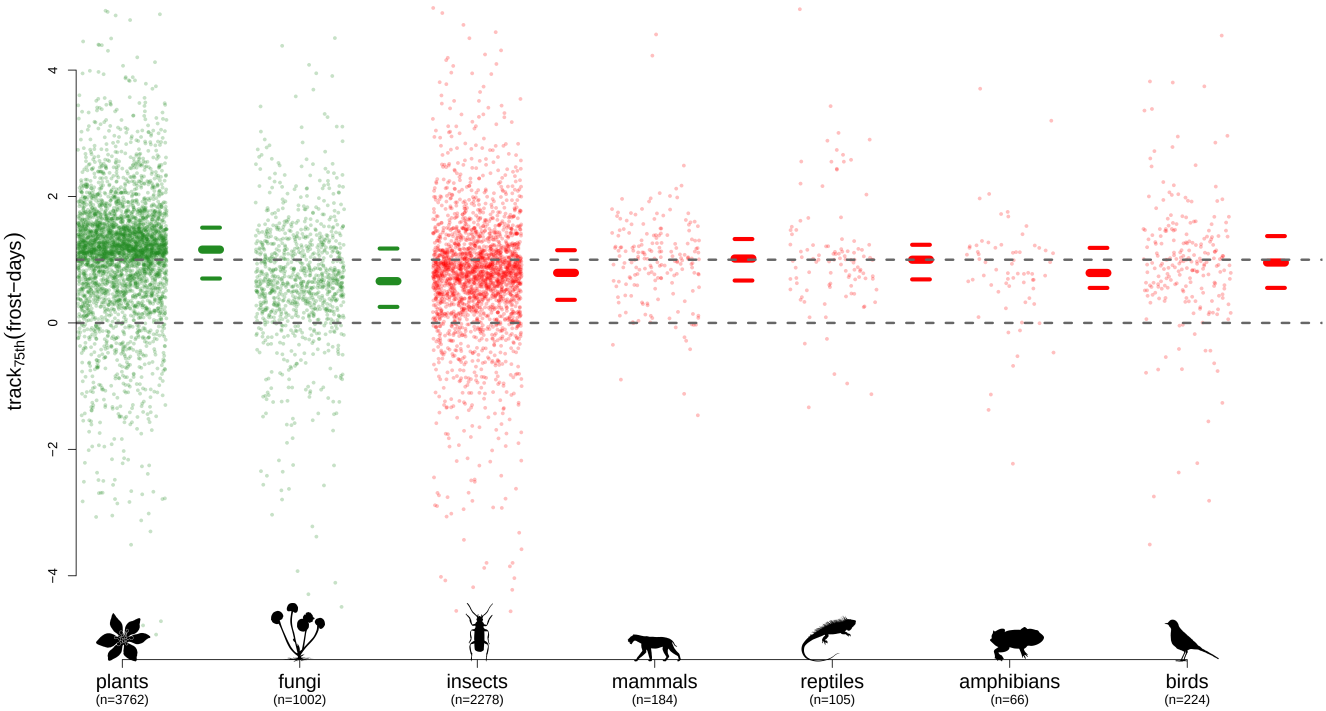


**Figure 9: Frost days, 75th quantile.** Legend as figure 1 in main text.

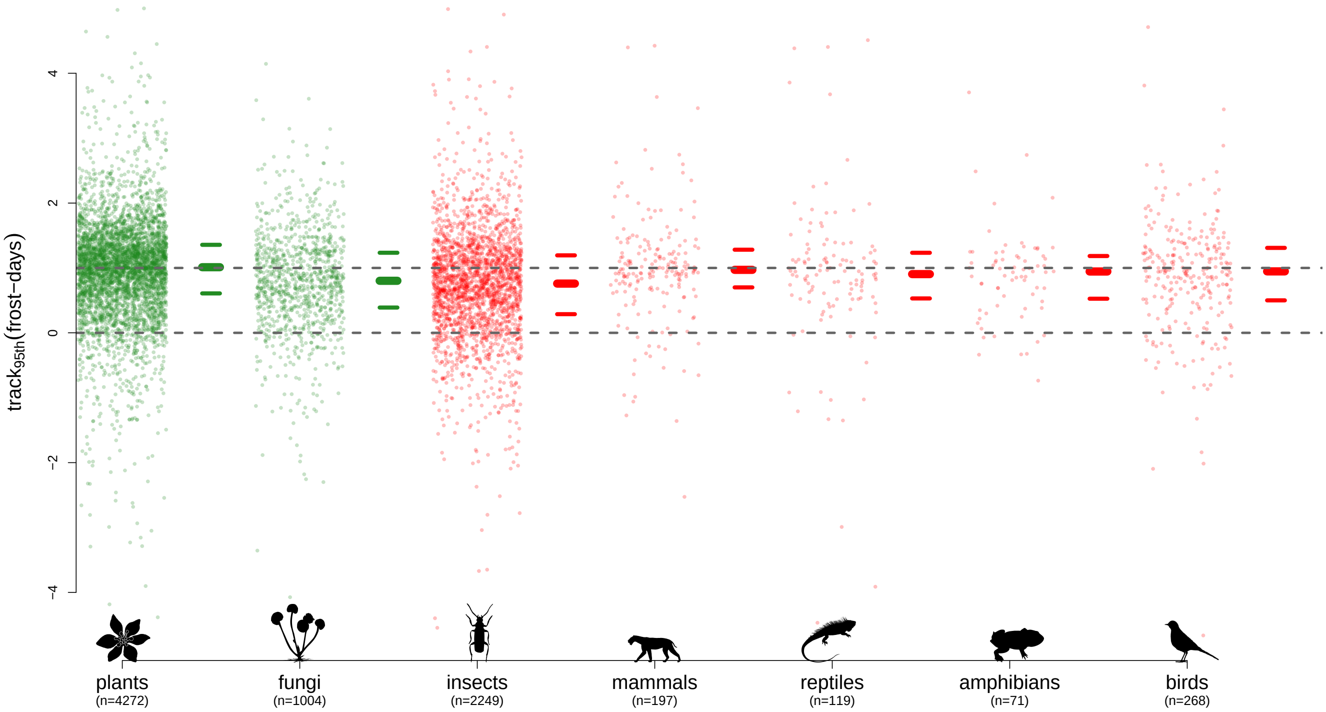


**Figure 10: Frost days, 95th quantile.** Legend as figure 1 in main text.

#### 1.3 Potential evapotranspiration

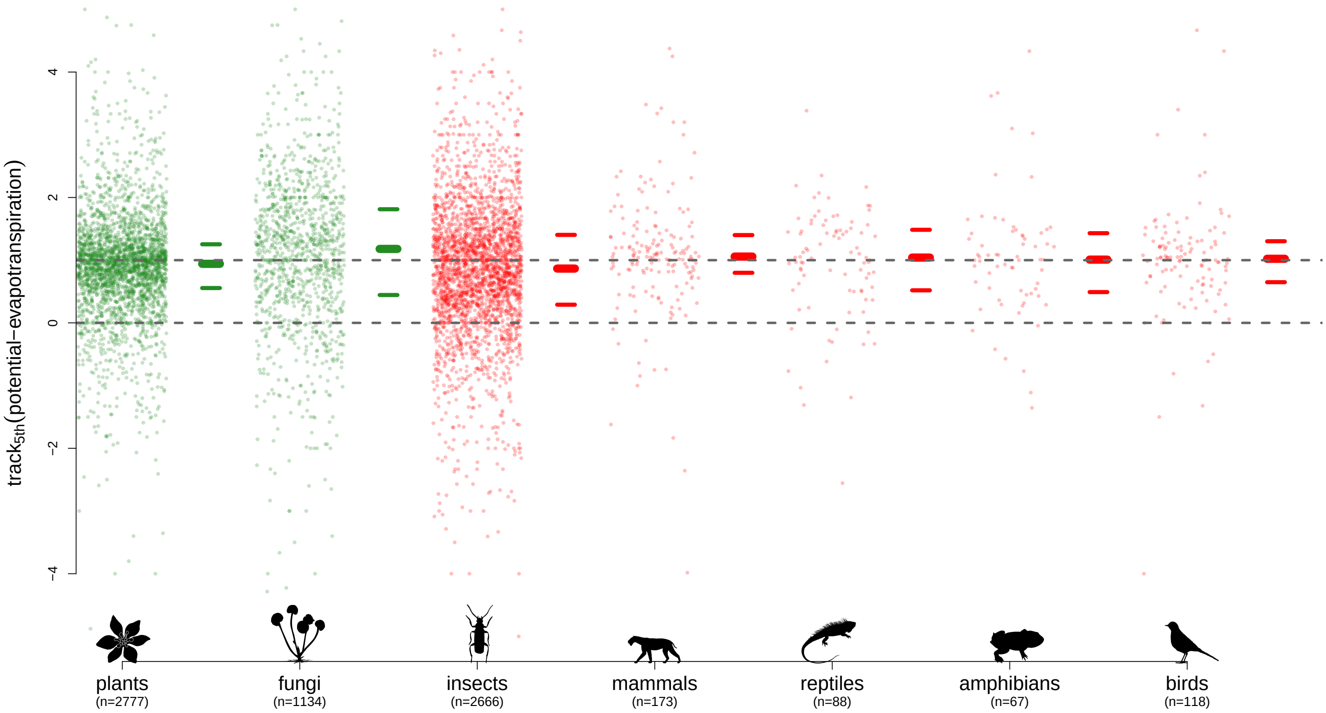


**Figure 11: Potential evapotranspiration, 5th quantile.** Legend as figure 1 in main text.

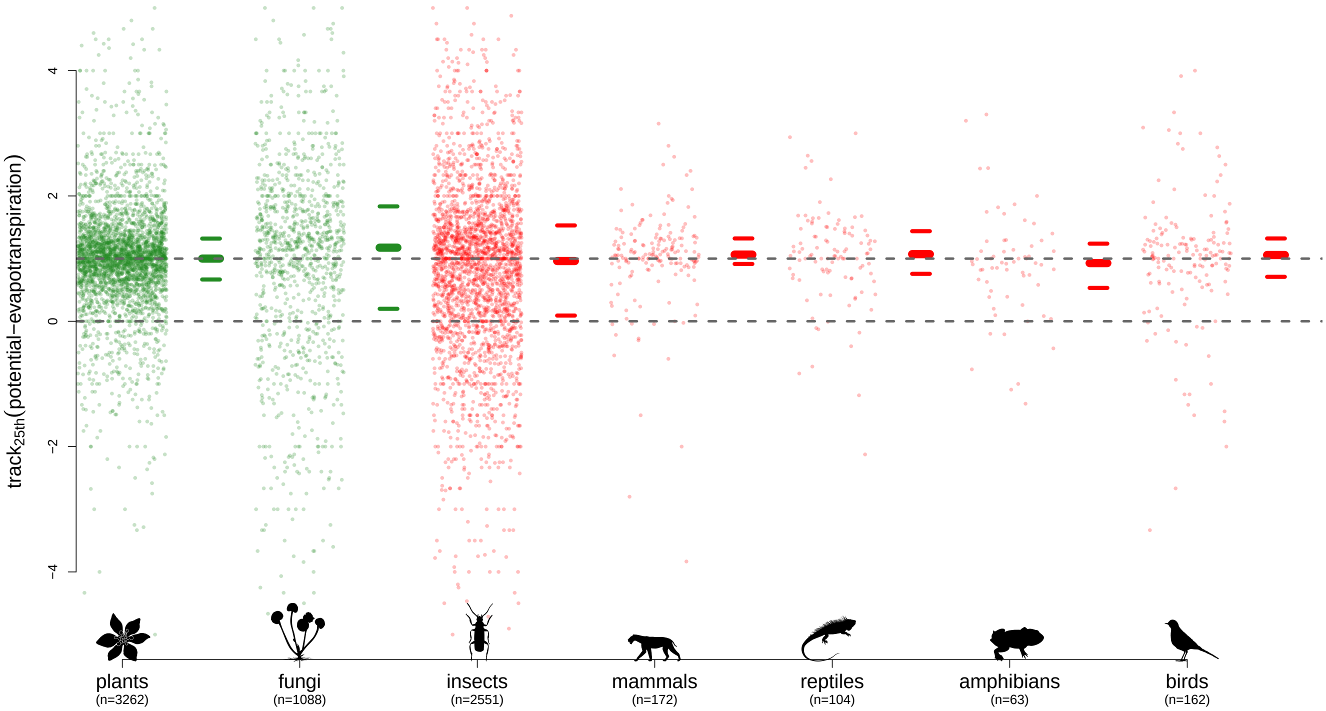


**Figure 12: Potential evapotranspiration, 25th quantile.** Legend as figure 1 in main text.

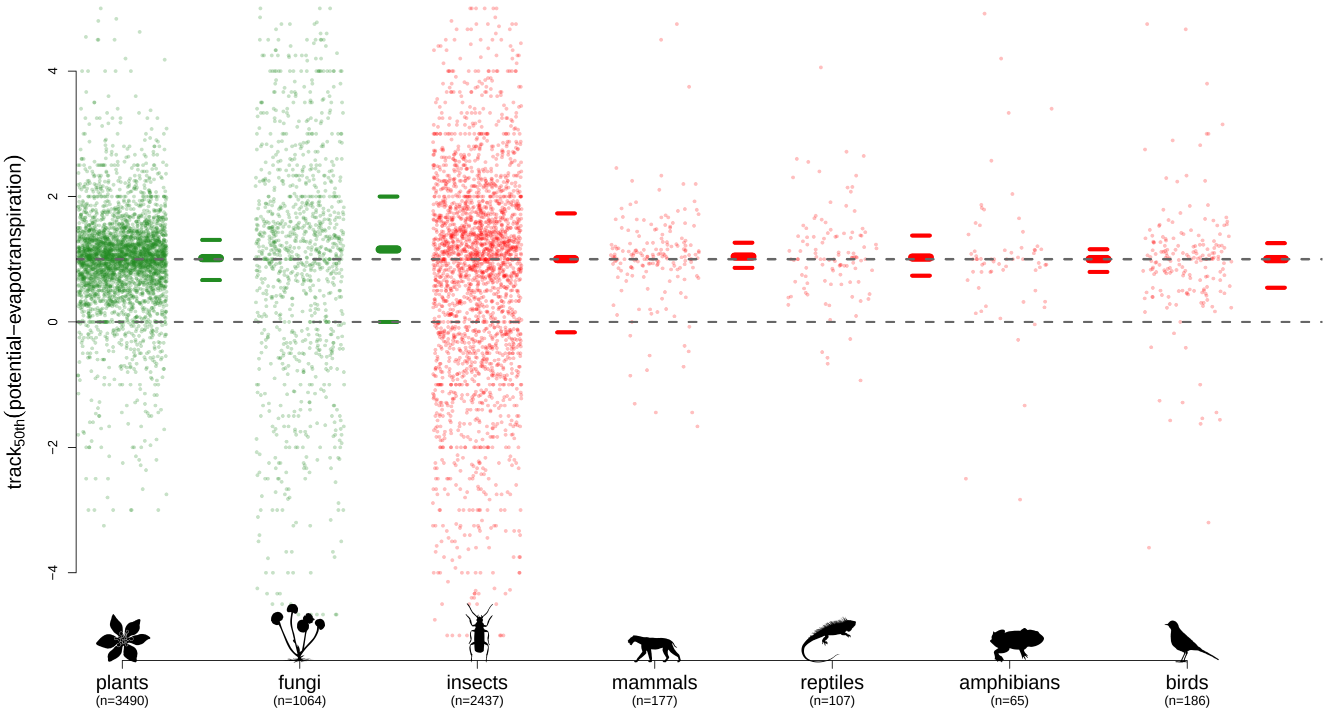


**Figure 13: Potential evapotranspiration, 50th quantile (median).** Legend as figure 1 in main text.

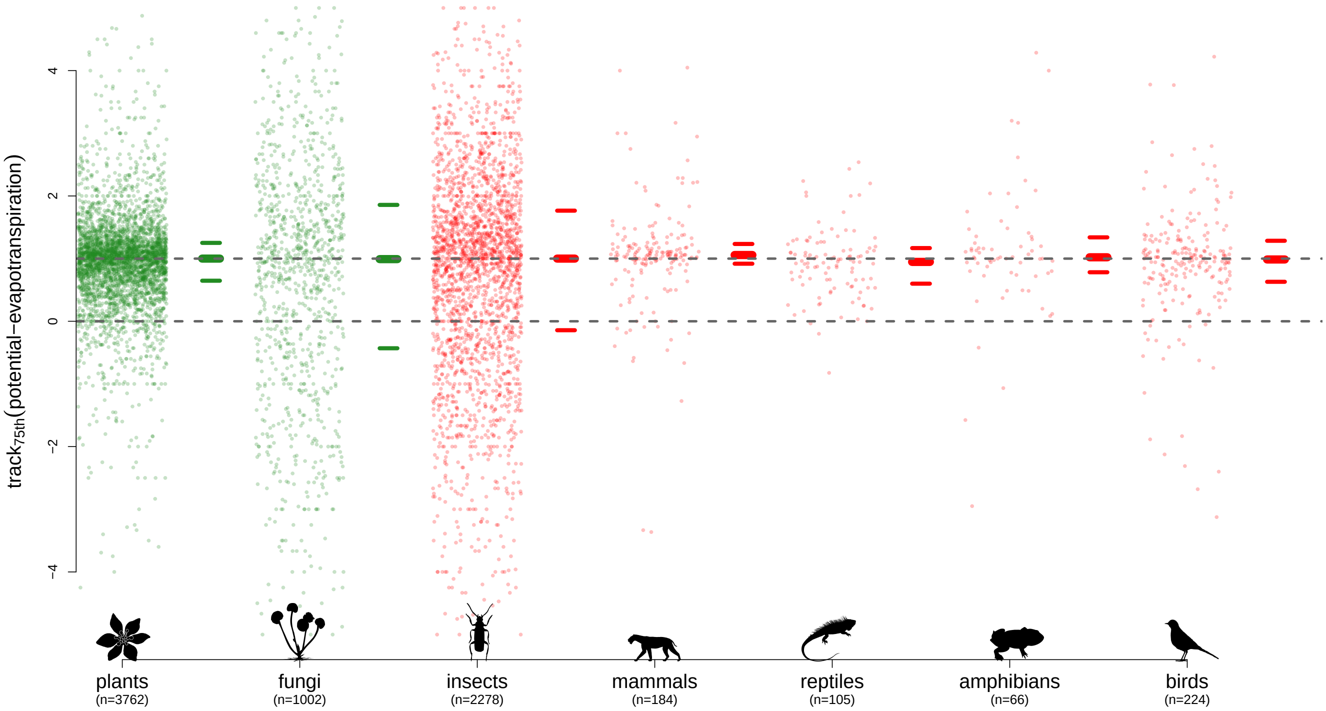


**Figure 14: Potential evapotranspiration, 75th quantile.** Legend as figure 1 in main text.

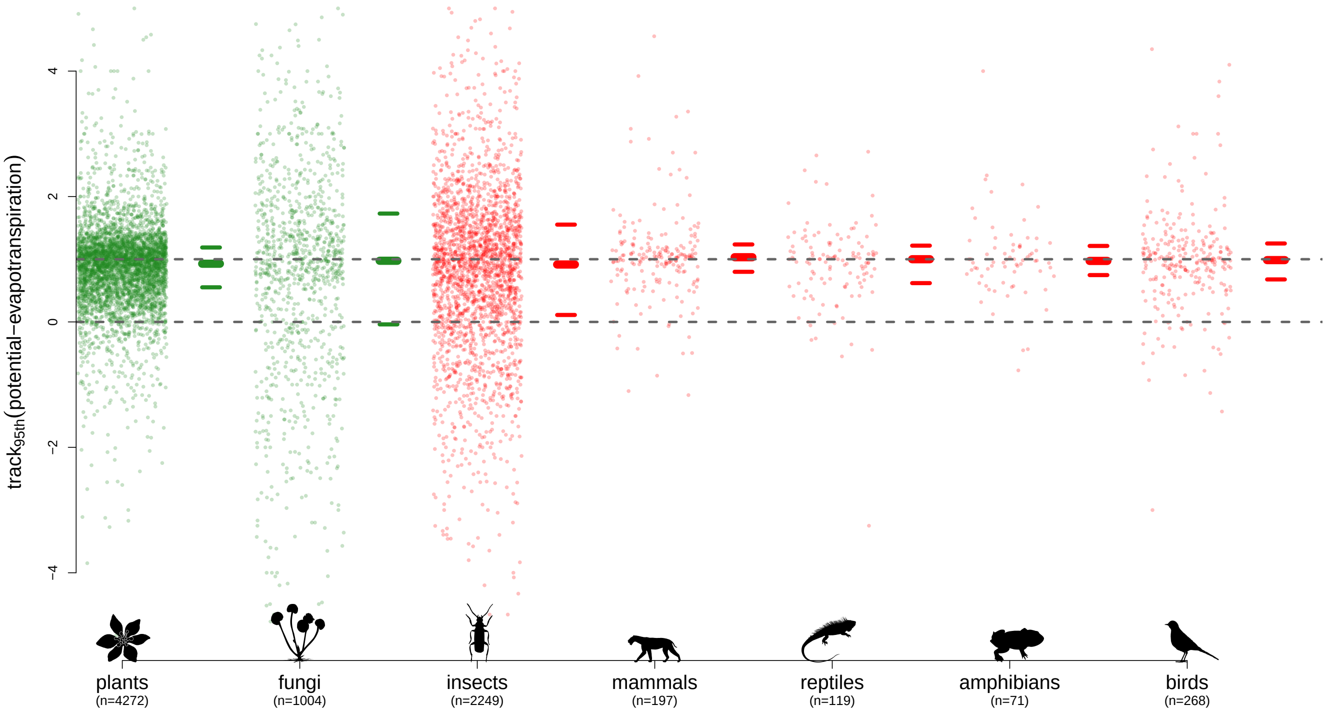


**Figure 15: Potential evapotranspiration, 95th quantile.** Legend as figure 1 in main text.

#### 1.4 Precipitation

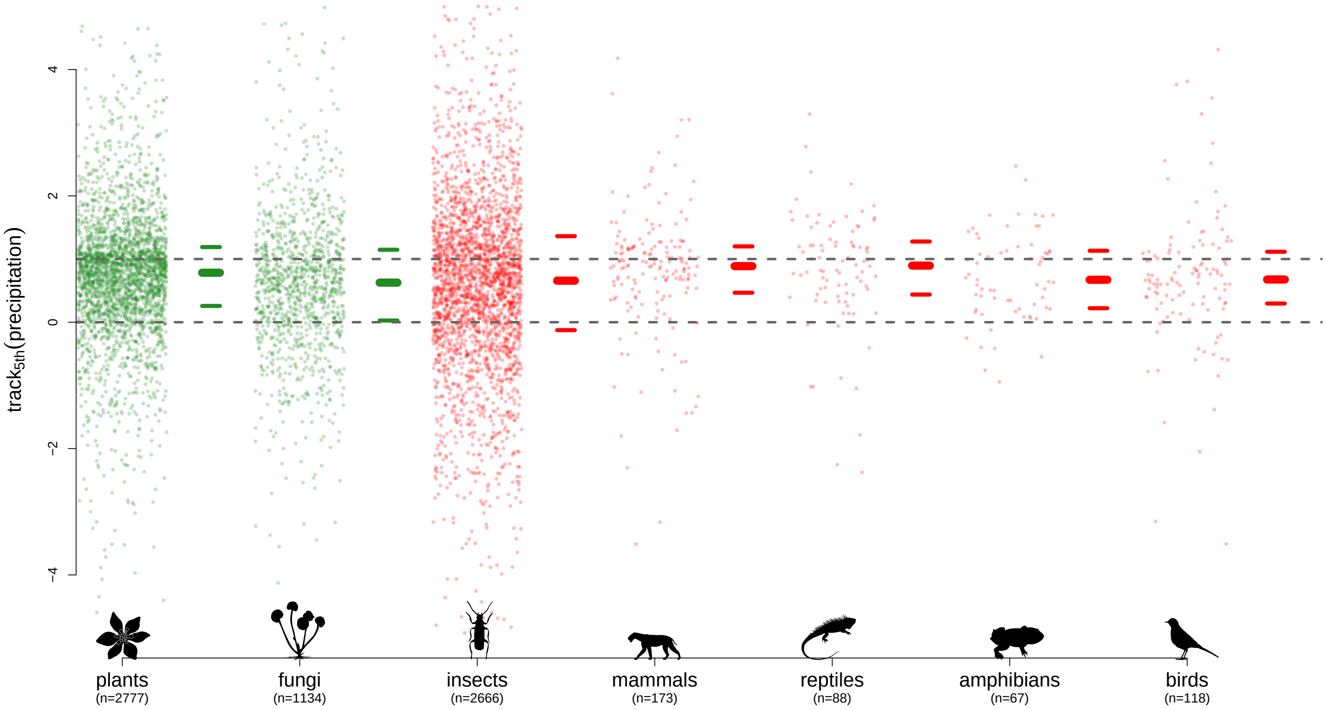


**Figure 16: Precipitation, 5th quantile.** Legend as figure 1 in main text.

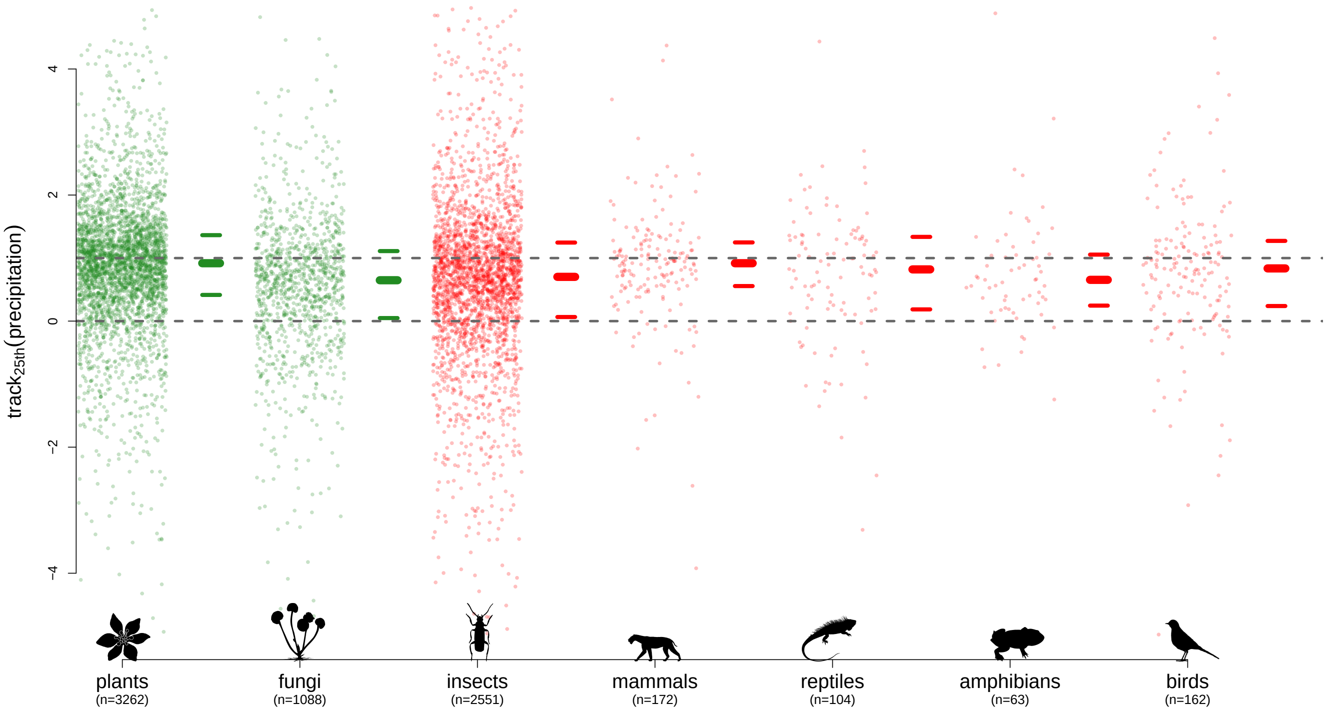


**Figure 17: Precipitation, 25th quantile.** Legend as figure 1 in main text.

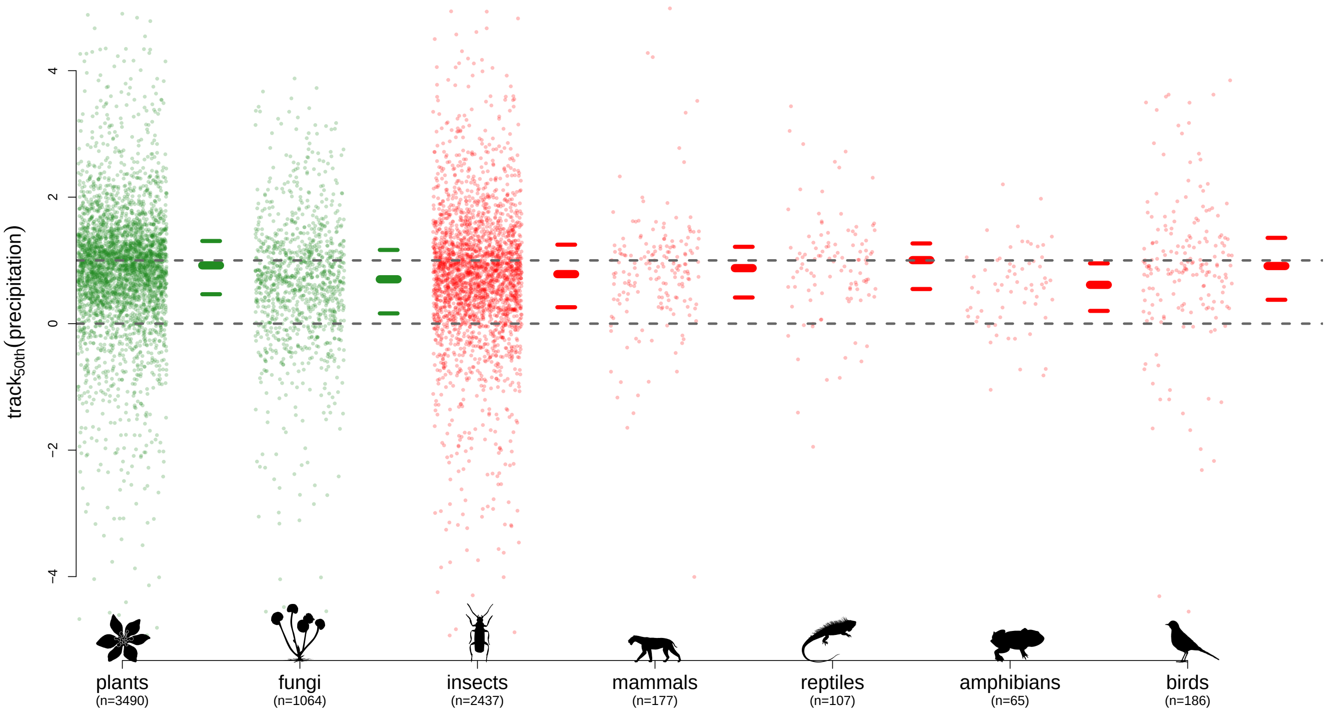


**Figure 18: Precipitation, 50th quantile (median).** Legend as figure 1 in main text.

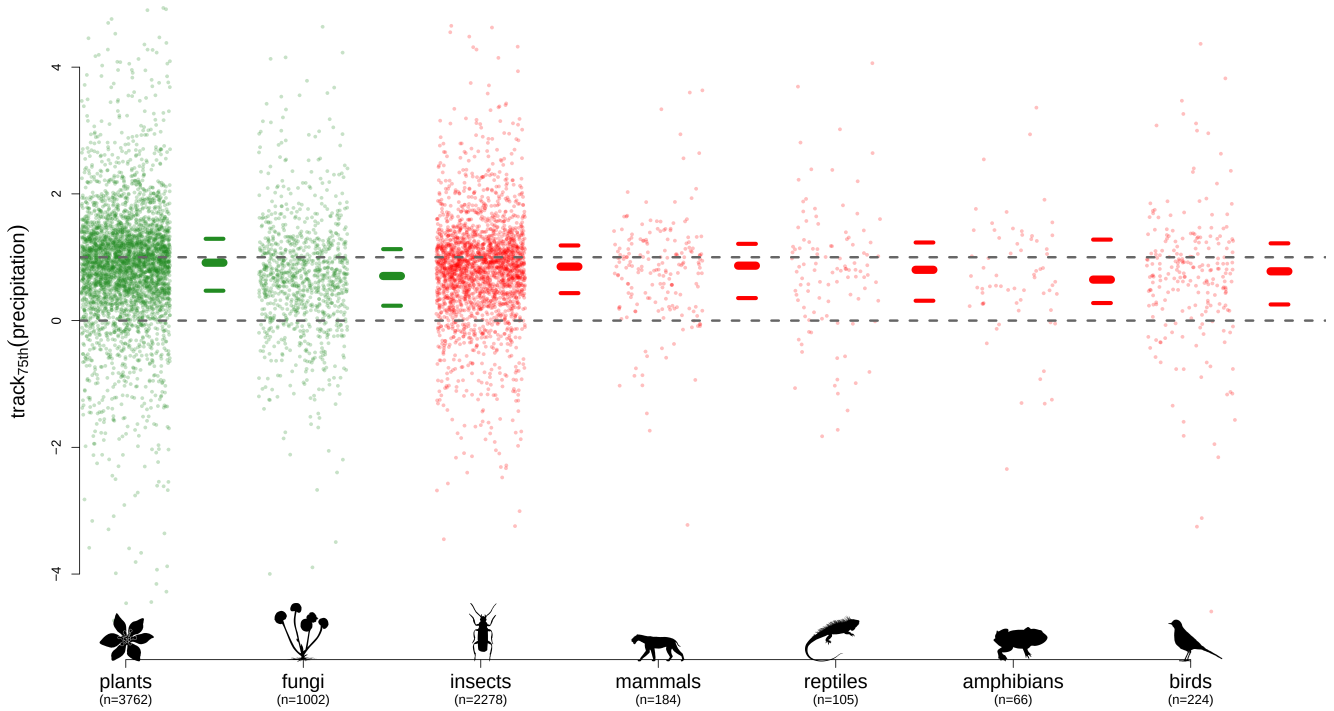


**Figure 19: Precipitation, 75th quantile.** Legend as figure 1 in main text.

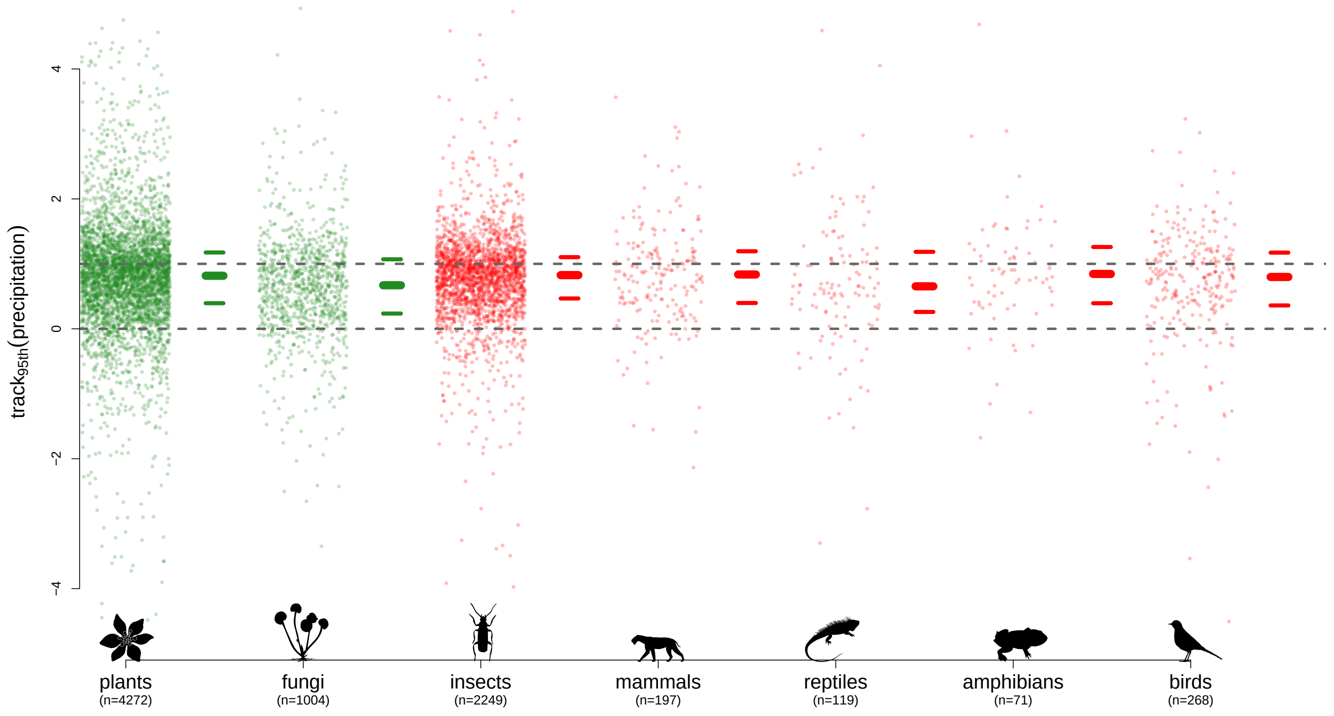


**Figure 20: Precipitation, 95th quantile.** Legend as figure 1 in main text.

#### 1.5 Minimum temperature

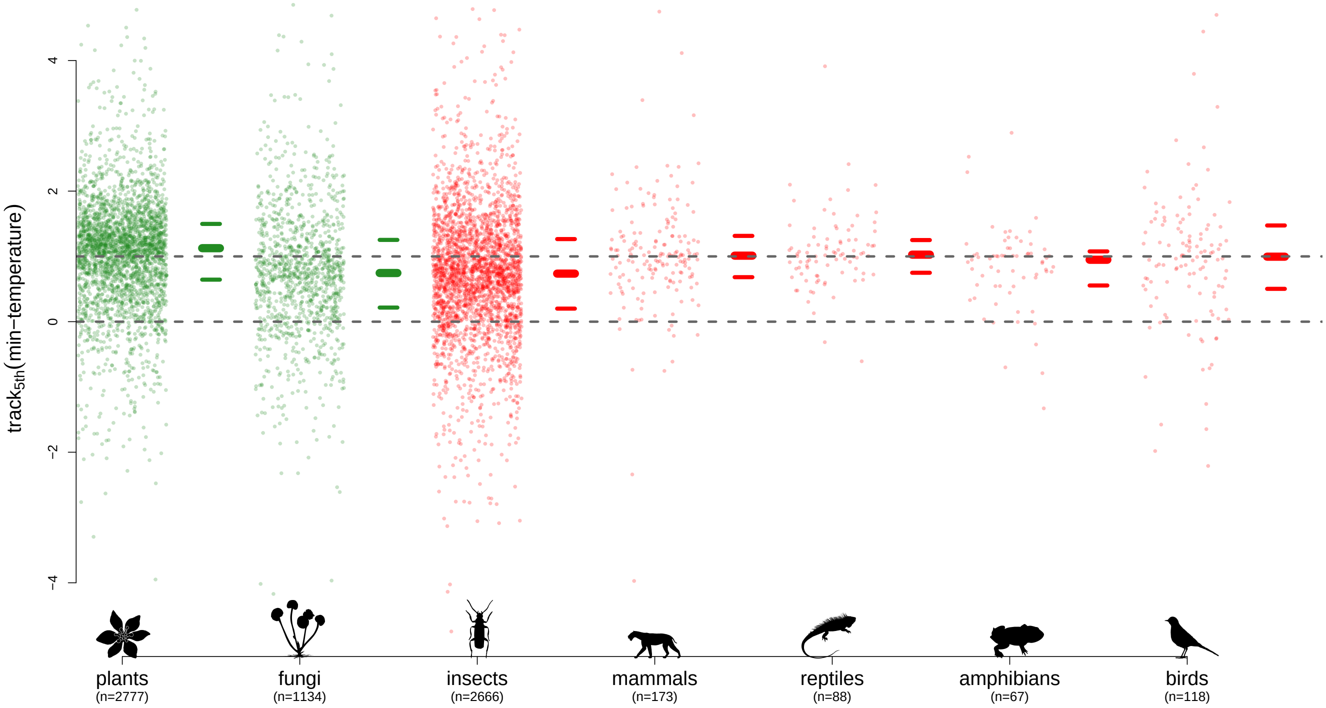


**Figure 21: Minimum temperature, 5th quantile.** Legend as figure 1 in main text.

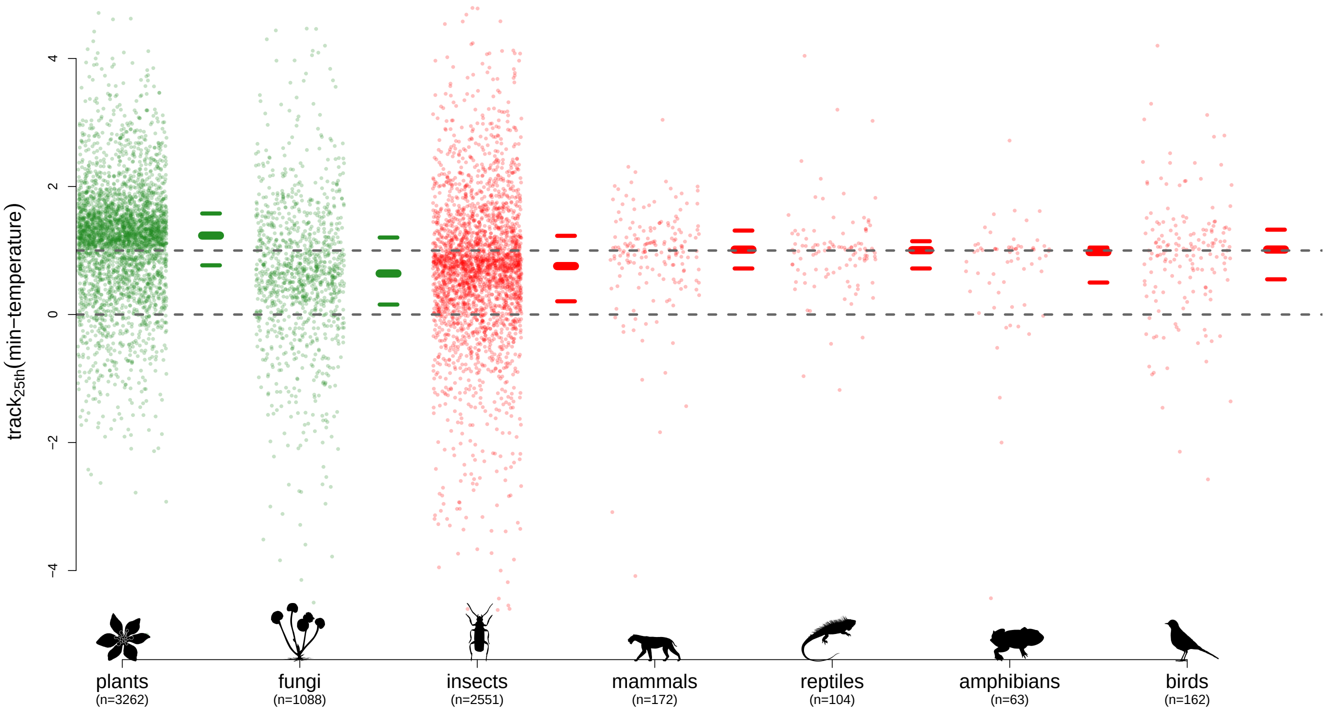


**Figure 22: Minimum temperature, 25th quantile.** Legend as figure 1 in main text.

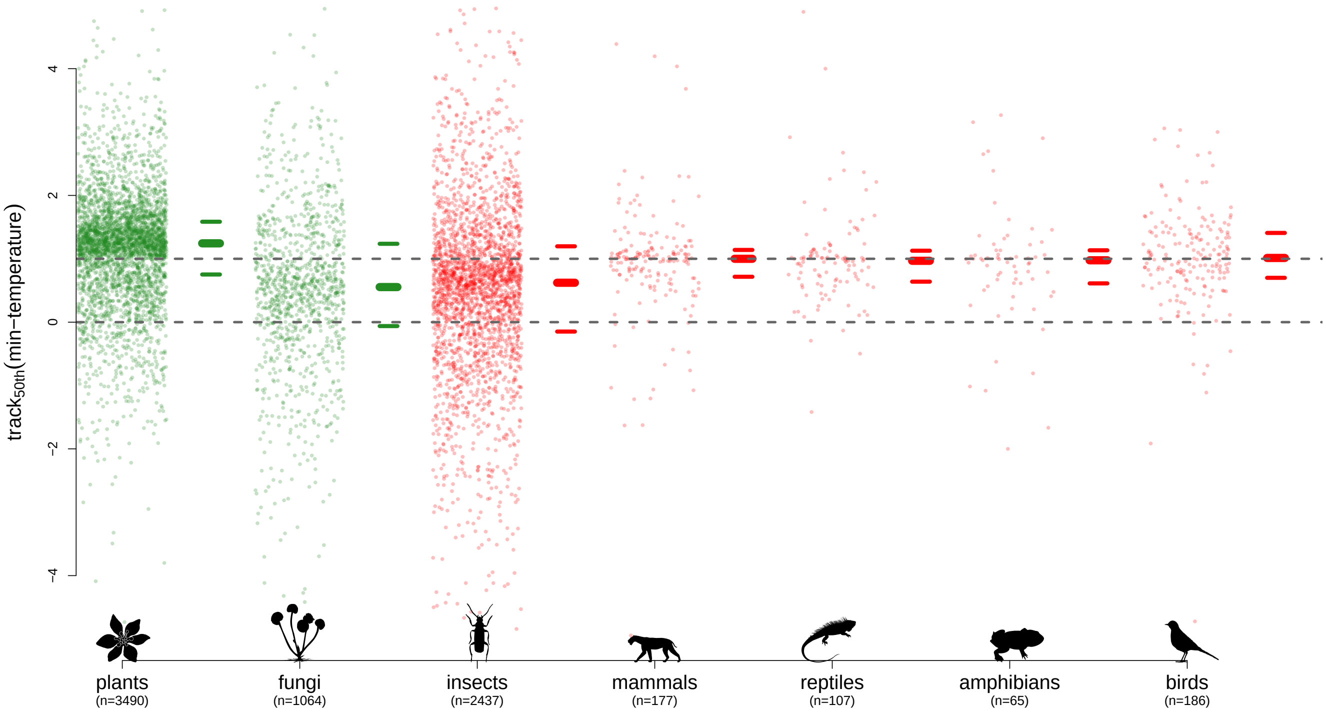


**Figure 23: Minimum temperature, 50th quantile (median).** Legend as figure 1 in main text.

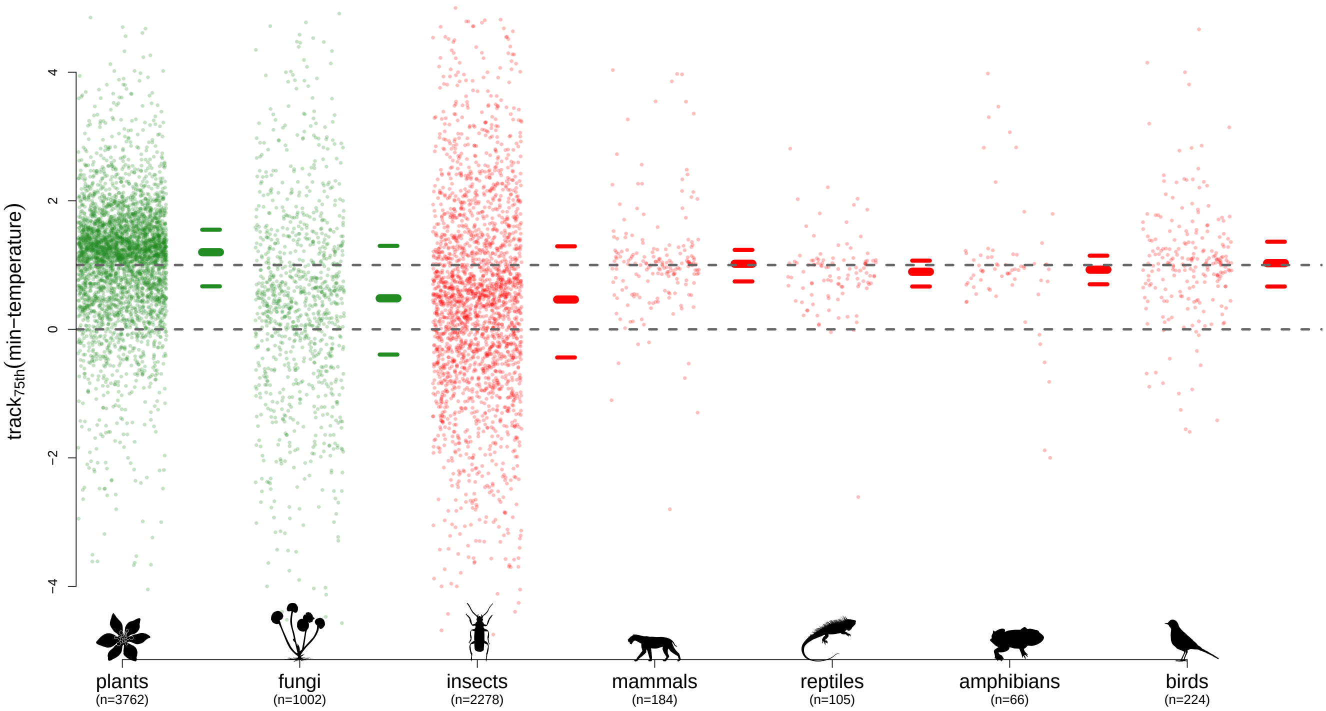


**Figure 24: Minimum temperature, 75th quantile.** Legend as figure 1 in main text.

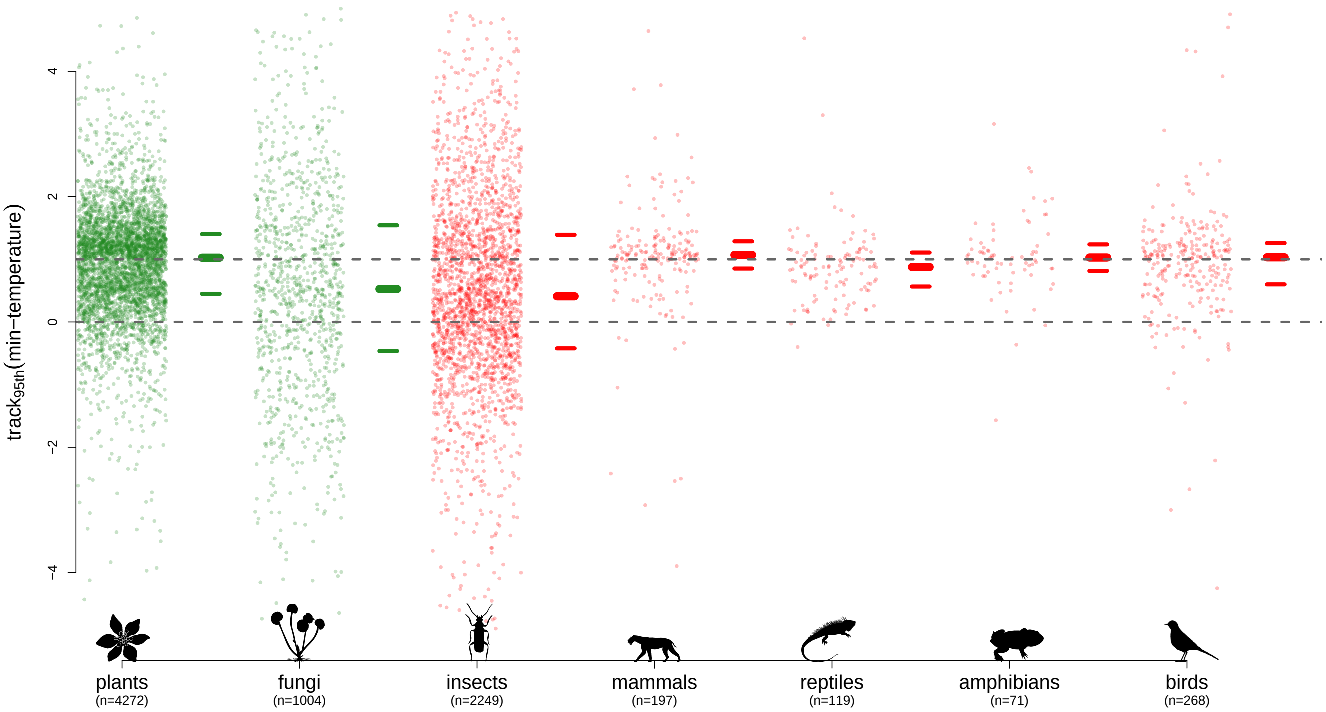


**Figure 25: Minimum temperature, 95th quantile.** Legend as figure 1 in main text.

#### 1.6 Mean temperature

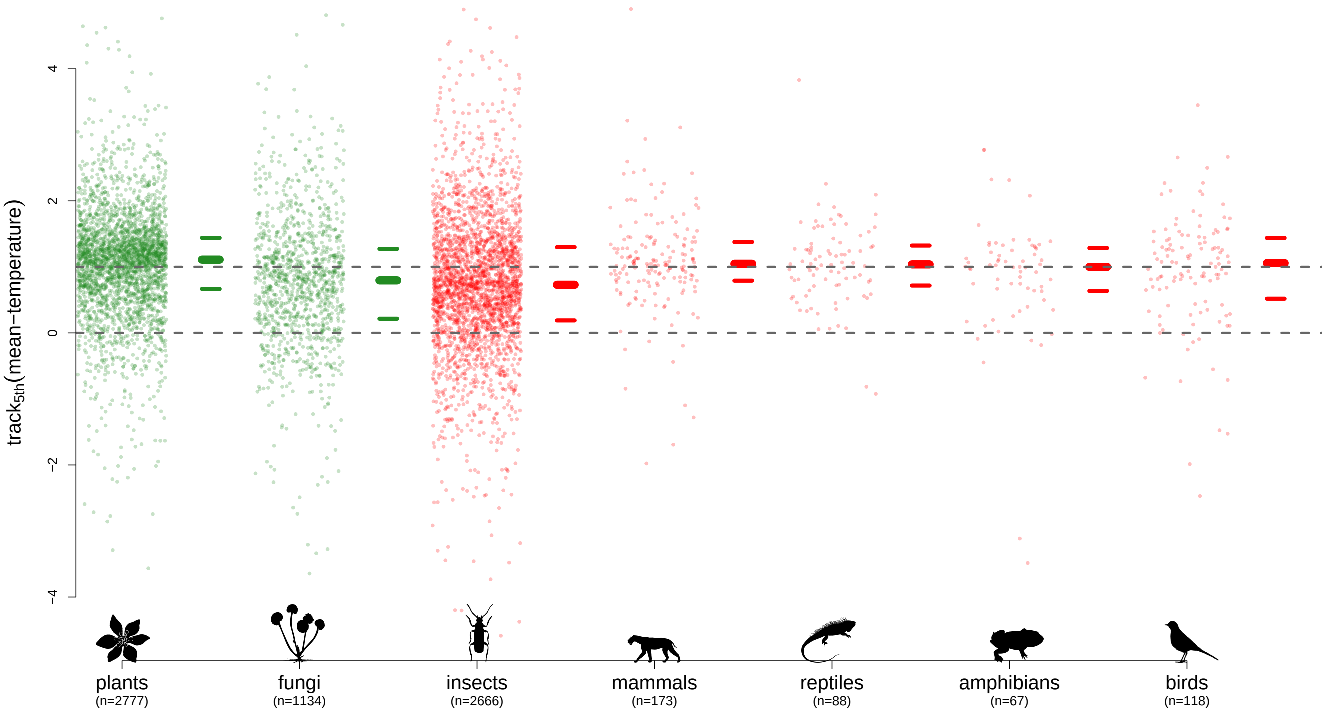


**Figure 26: Mean-temperature, 5th quantile.** Legend as figure 1 in main text.

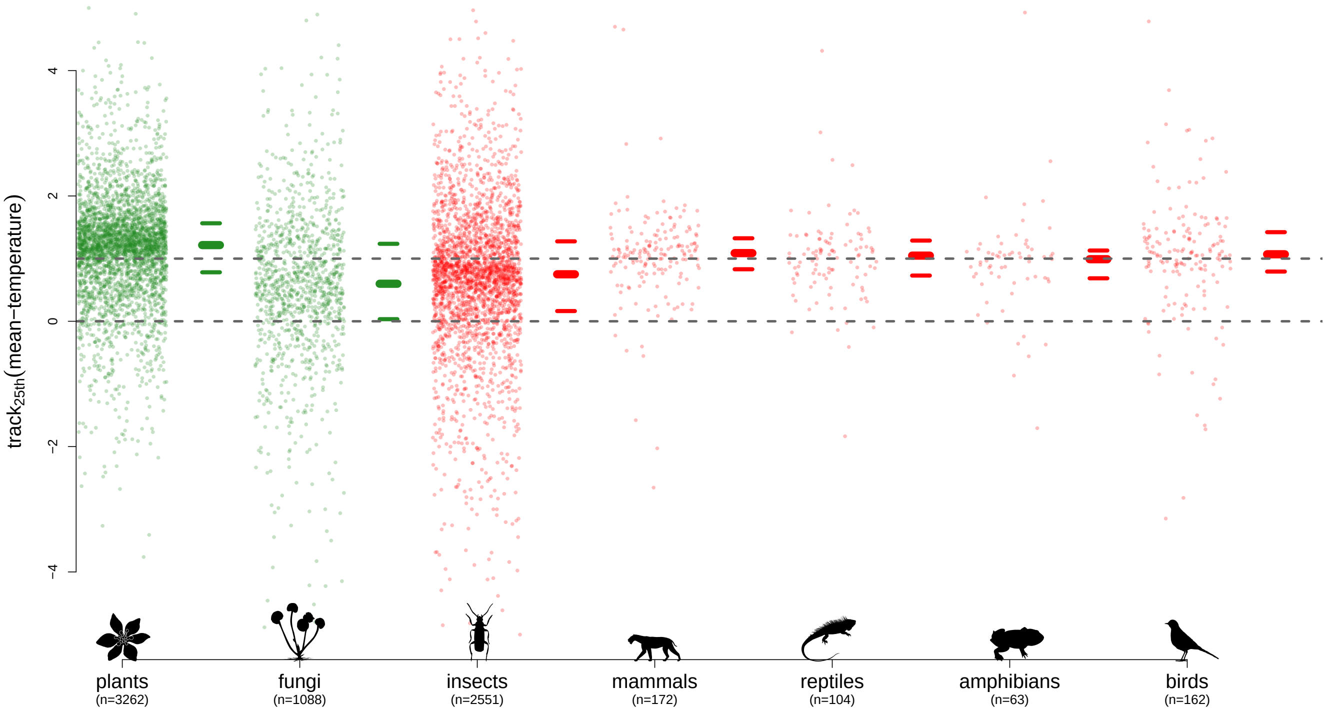


**Figure 27: Mean-temperature, 25th quantile.** Legend as figure 1 in main text.

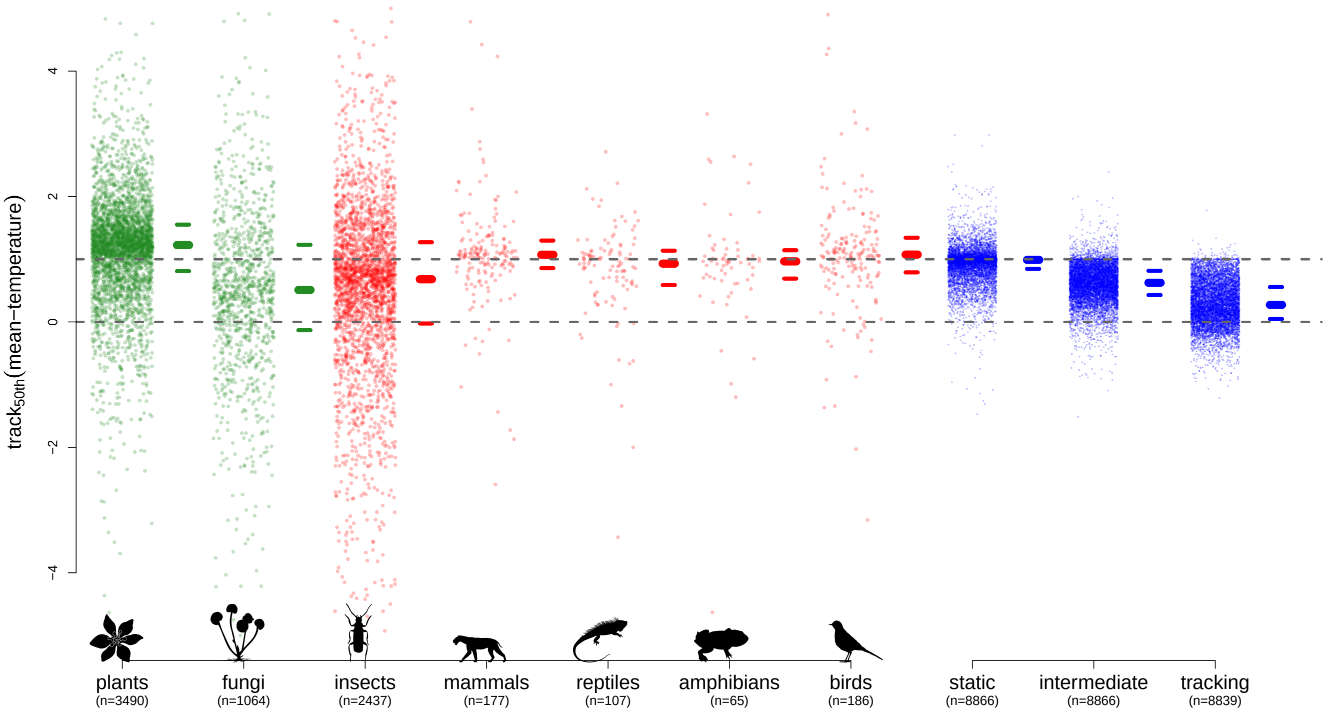


**Figure 28: Mean-temperature, 50th quantile (median).** Legend as figure 1 in main text.

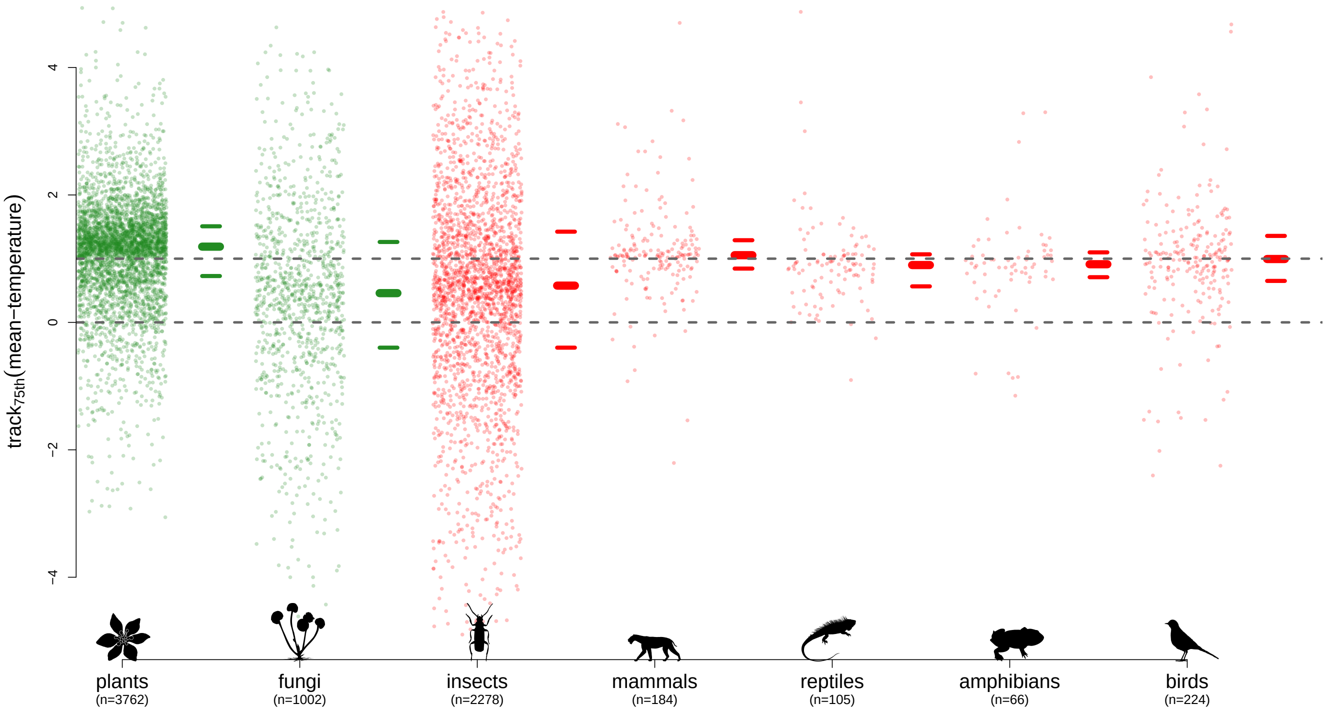


**Figure 29: Mean-temperature, 75th quantile.** Legend as figure 1 in main text.

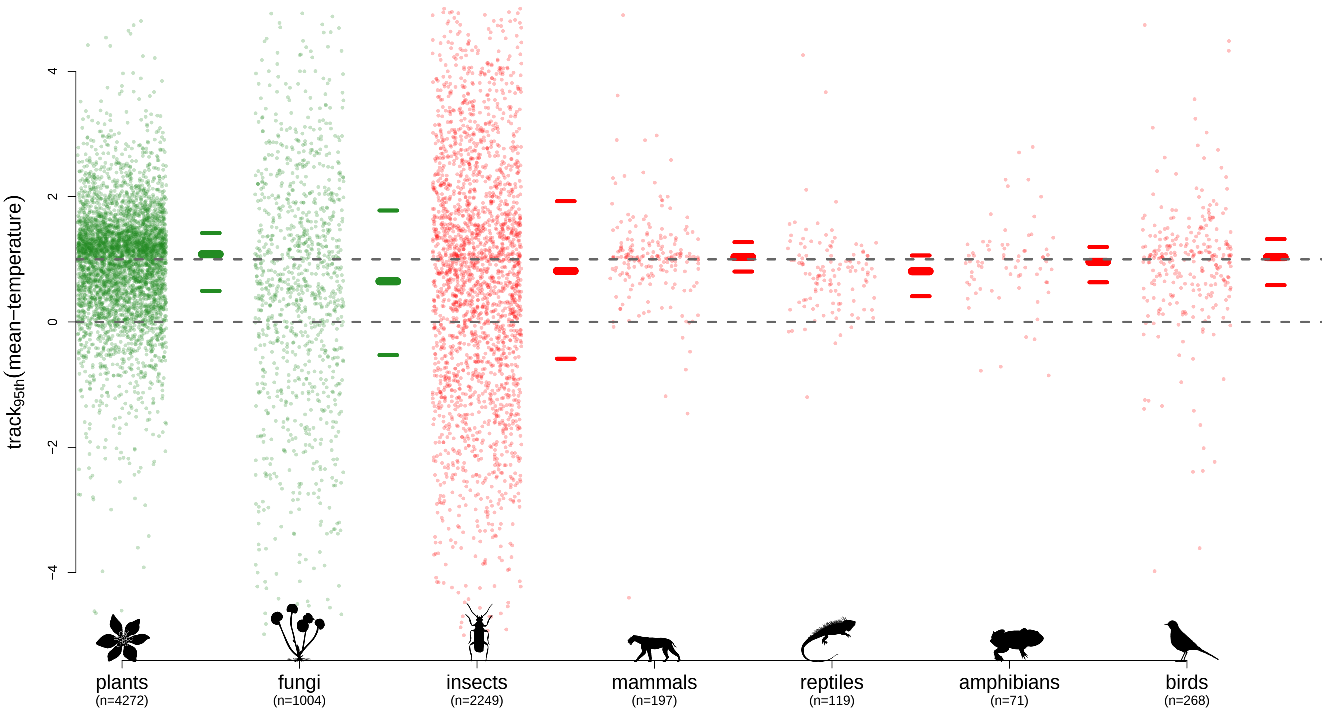


**Figure 30: Mean-temperature, 95th quantile.** Legend as figure 1 in main text.

#### 1.7 Maximum temperature

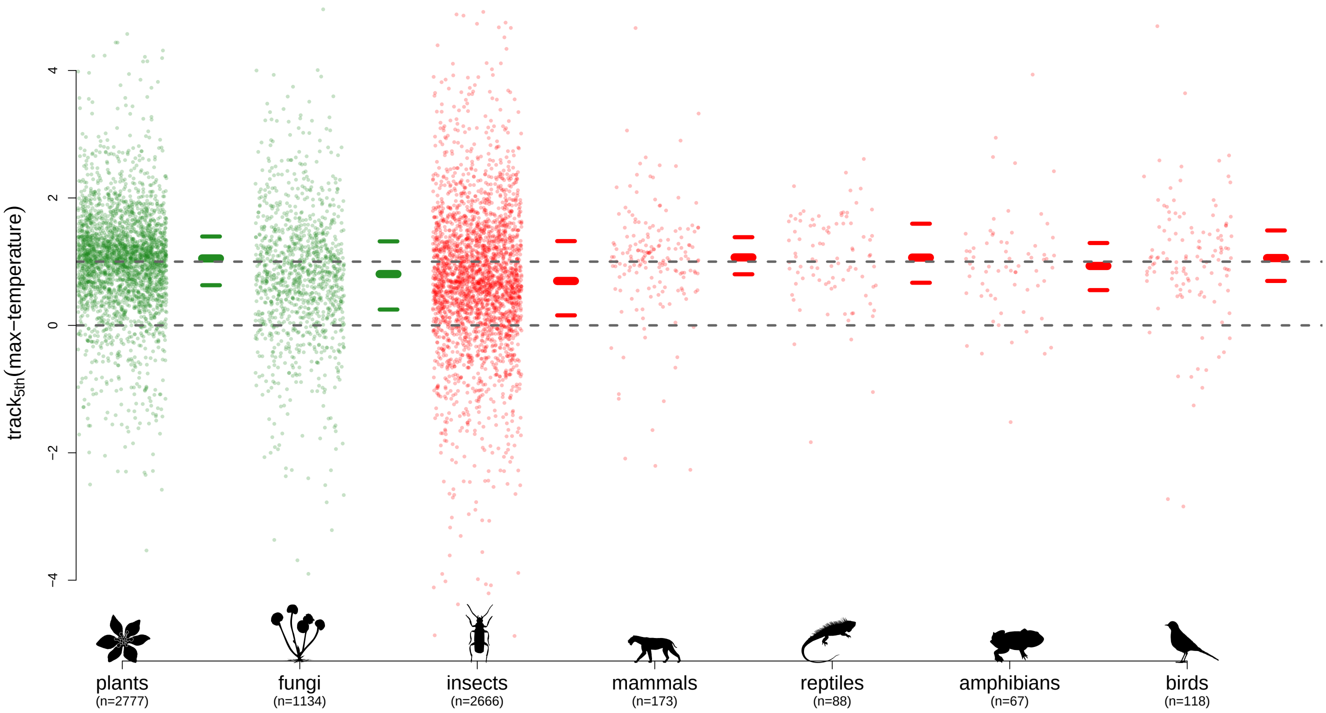


**Figure 31: Maximum temperature, 5th quantile.** Legend as figure 1 in main text.

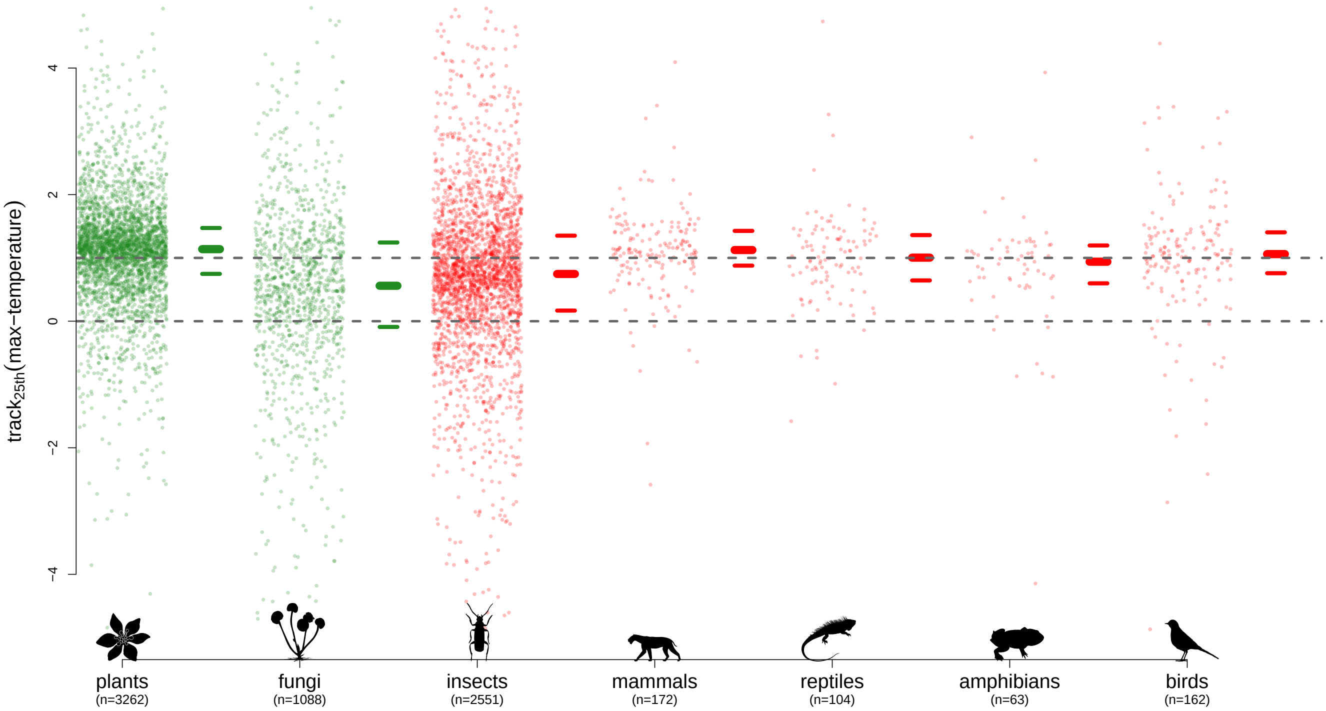


**Figure 32: Maximum temperature, 25th quantile.** Legend as figure 1 in main text.

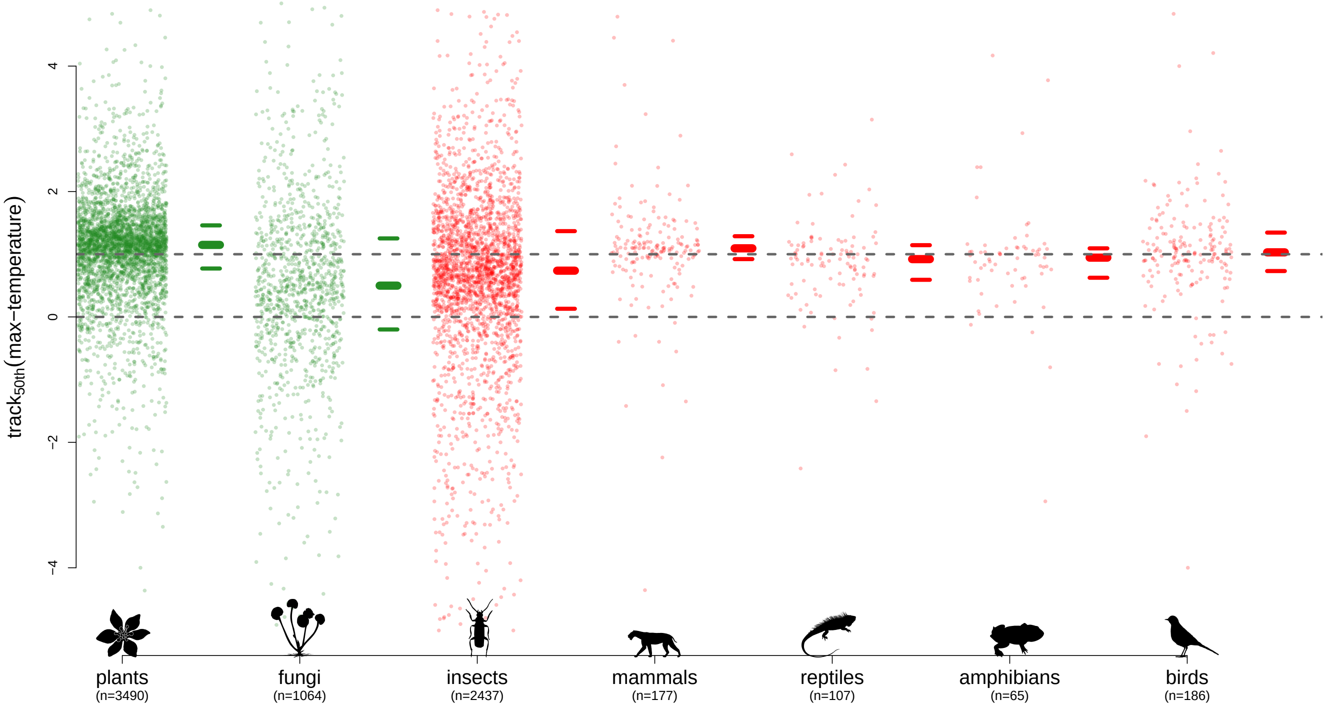


**Figure 33: Maximum temperature, 50th quantile (median).** Legend as figure 1 in main text.

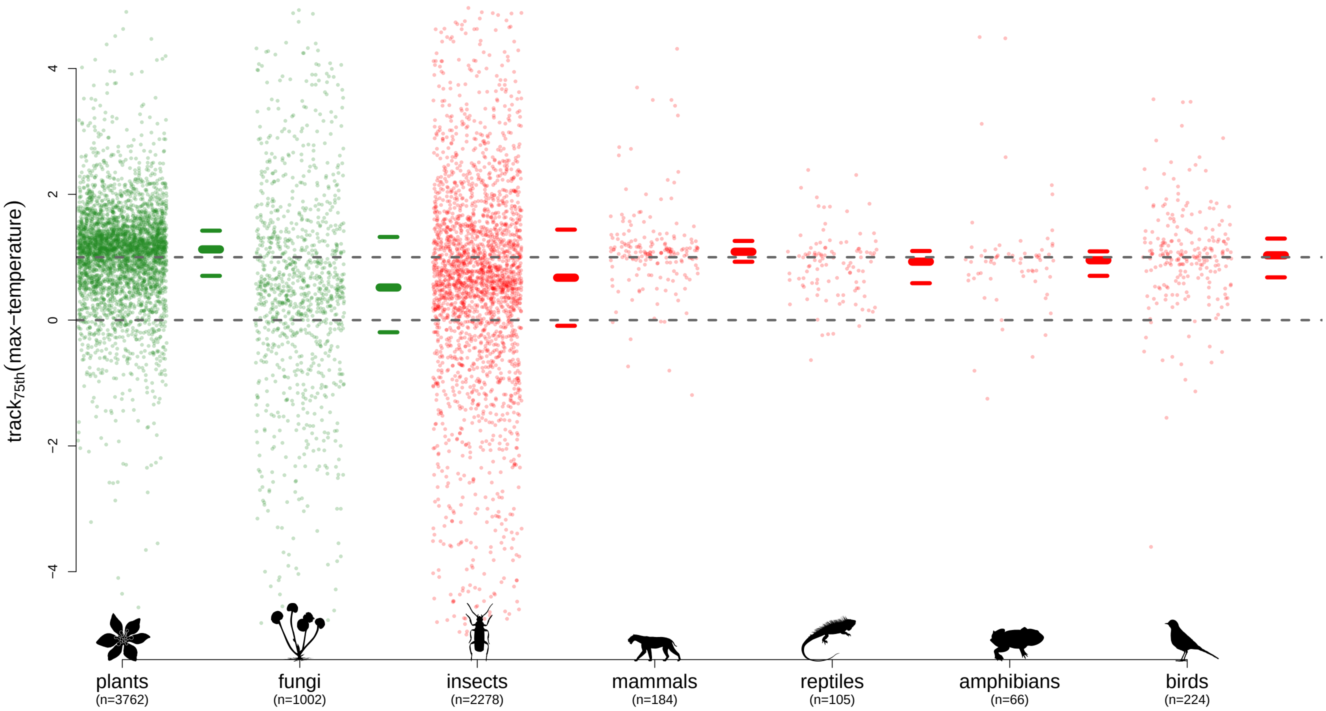


**Figure 34: Maximum temperature, 75th quantile.** Legend as figure 1 in main text.

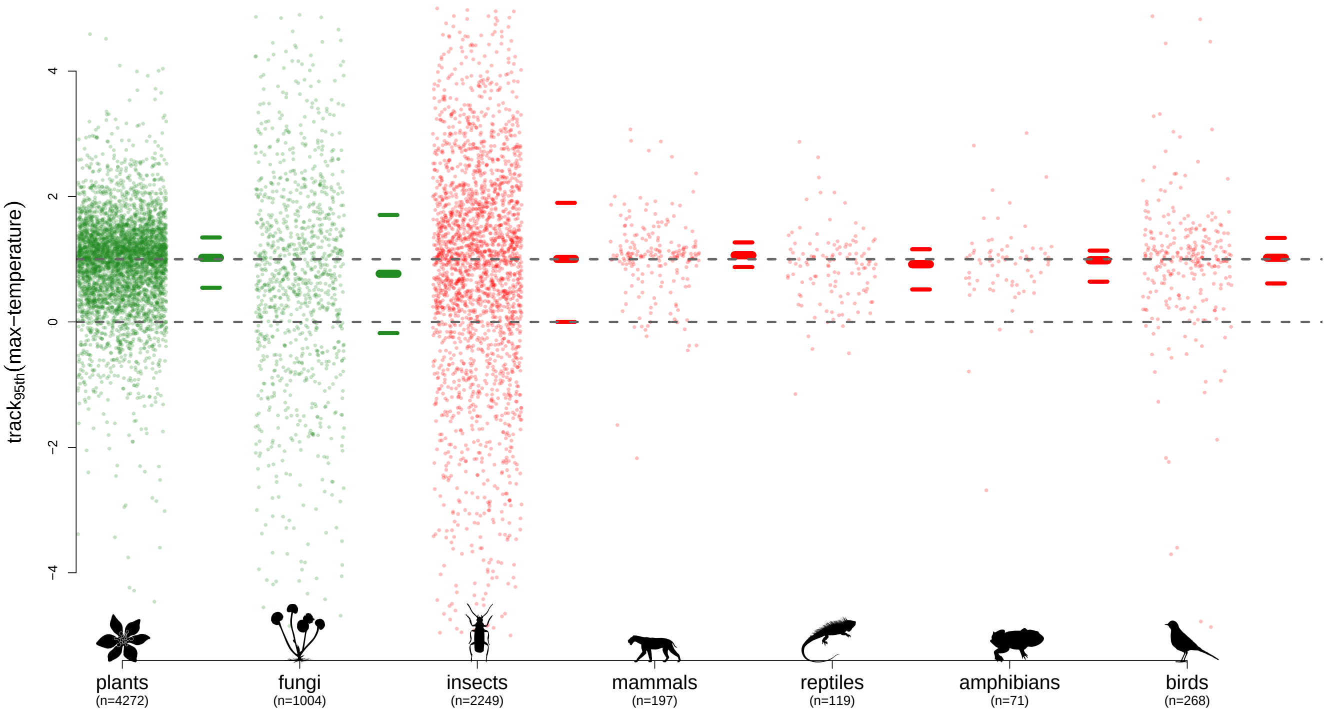


**Figure 35: Maximum temperature, 95th quantile.** Legend as figure 1 in main text.

#### 1.8 Vapour pressure

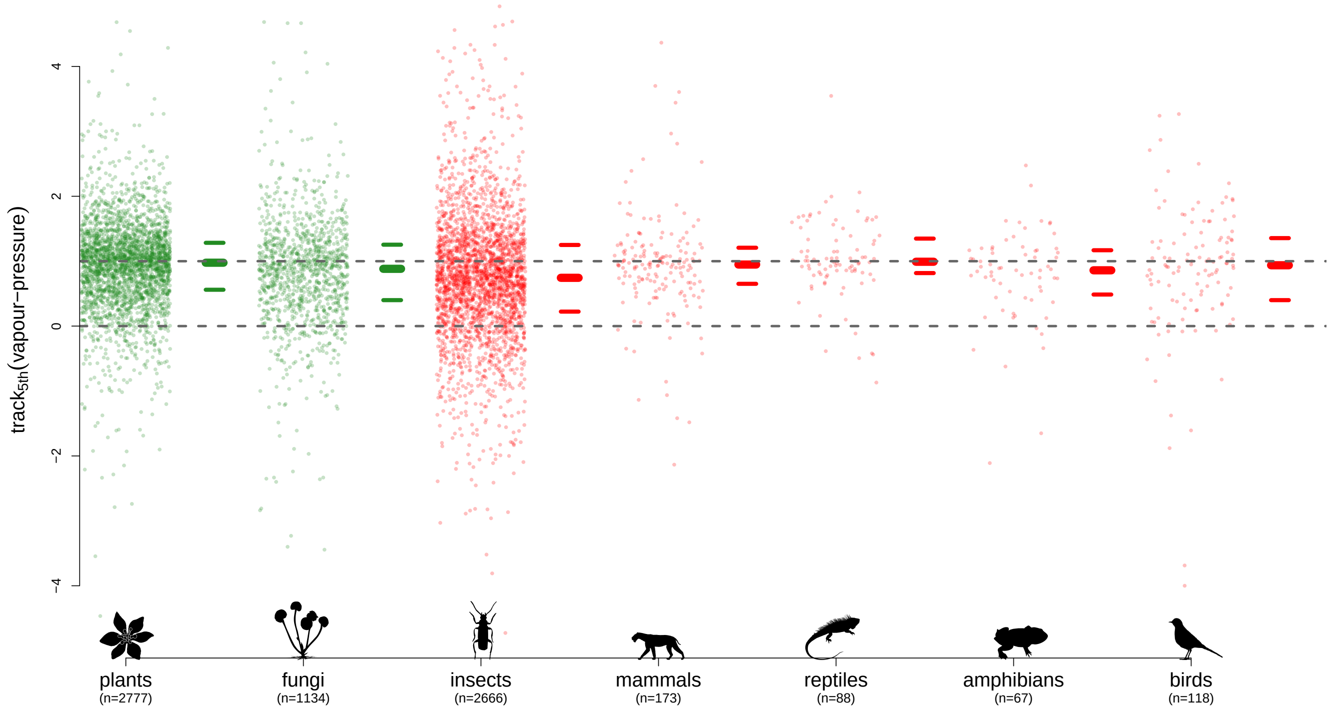


**Figure 36: Vapour pressure, 5th quantile.** Legend as figure 1 in main text.

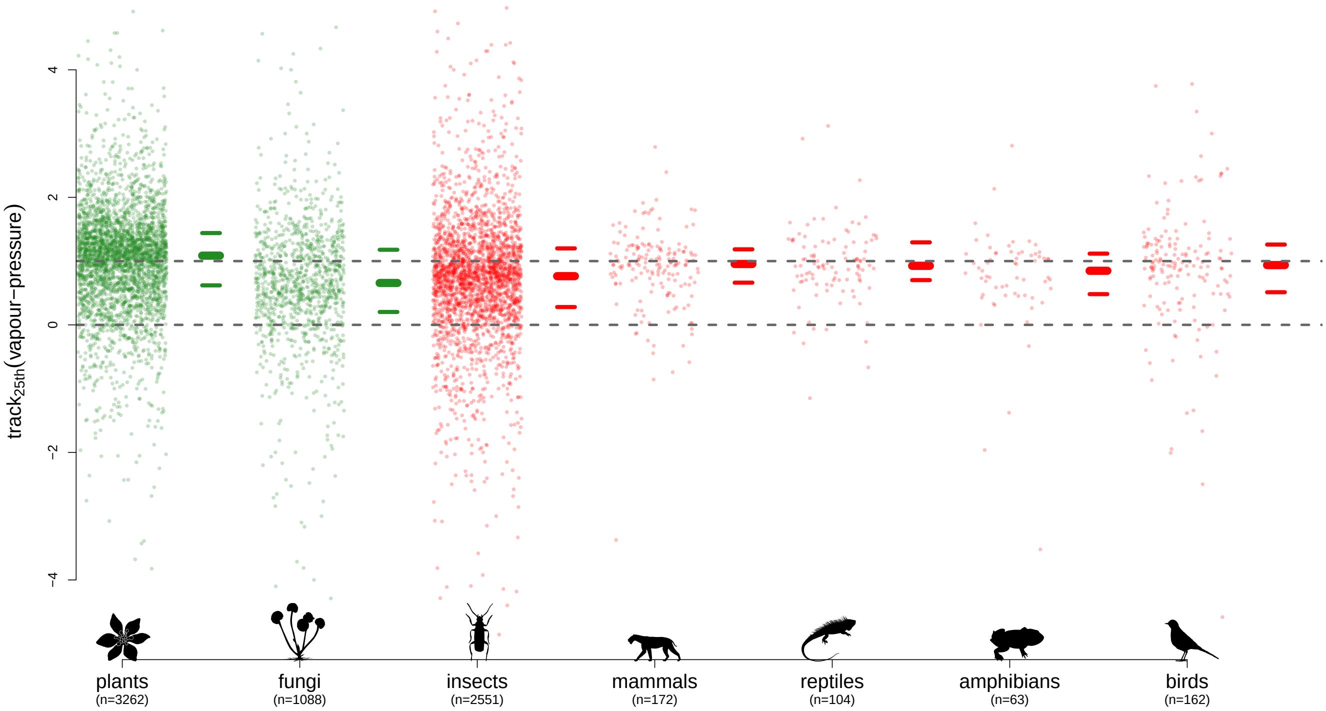


**Figure 37: Vapour pressure, 25th quantile.** Legend as figure 1 in main text.

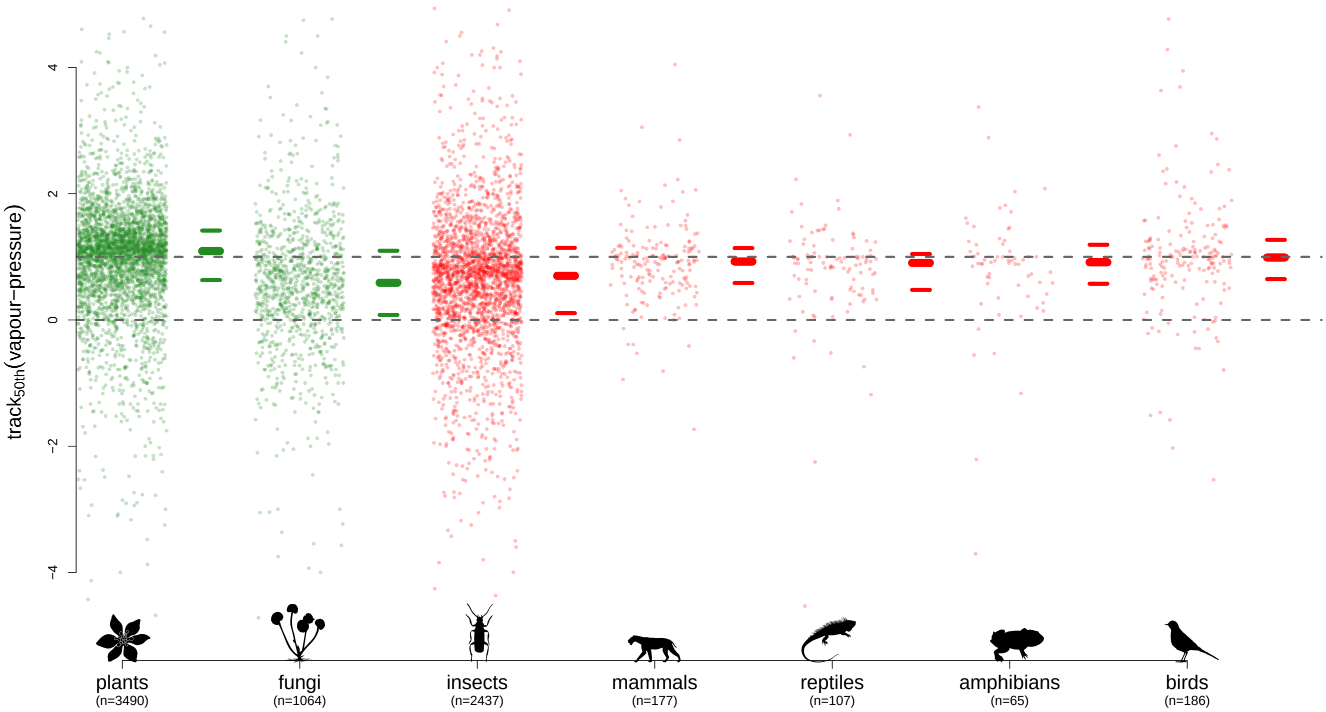


**Figure 38: Vapour pressure, 50th quantile (median).** Legend as figure 1 in main text.

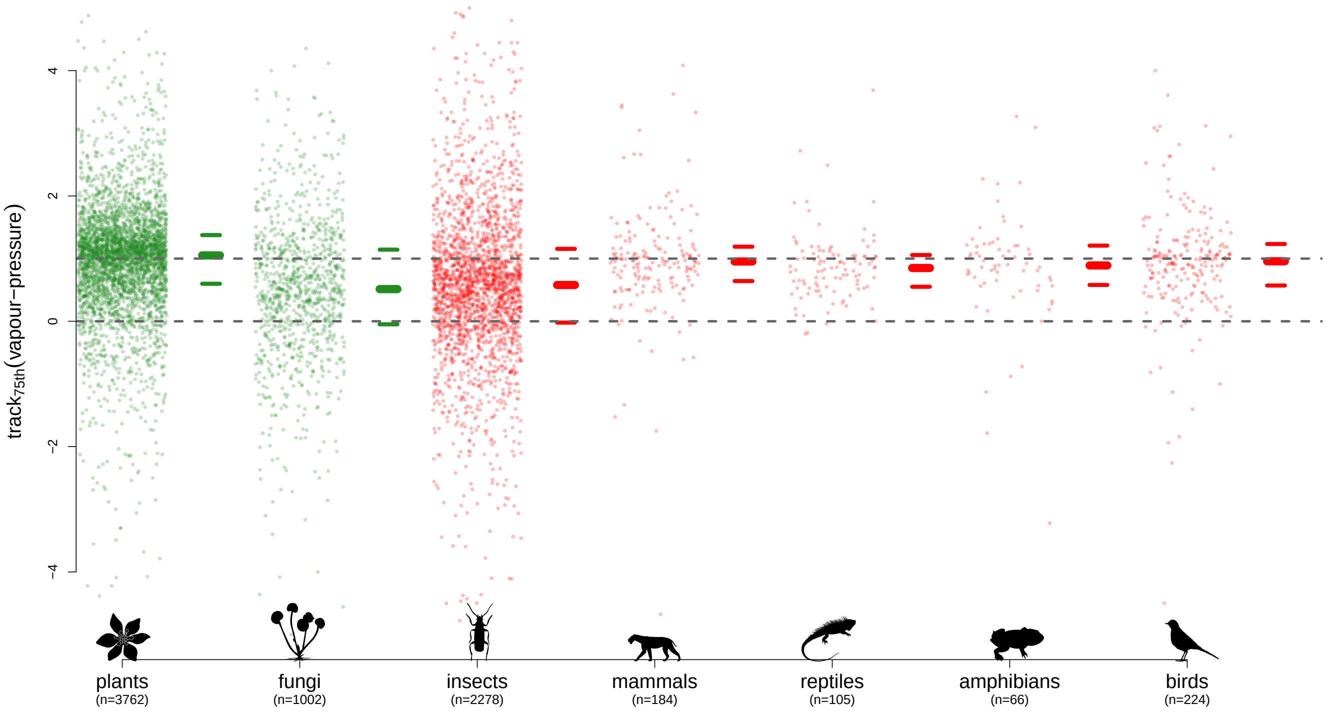


**Figure 39: Vapour pressure, 75th quantile.** Legend as figure 1 in main text.

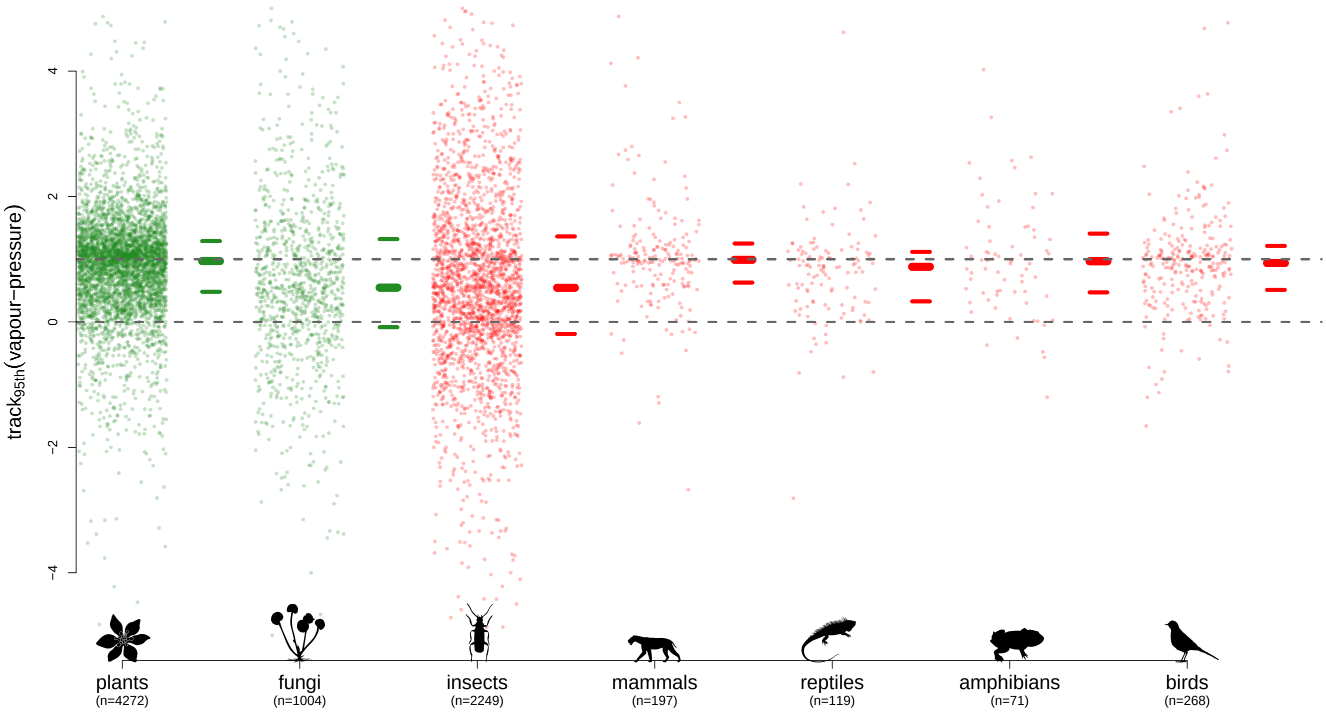


**Figure 40: Vapour pressure, 95th quantile.** Legend as figure 1 in main text.

#### 1.9 Rainday counts

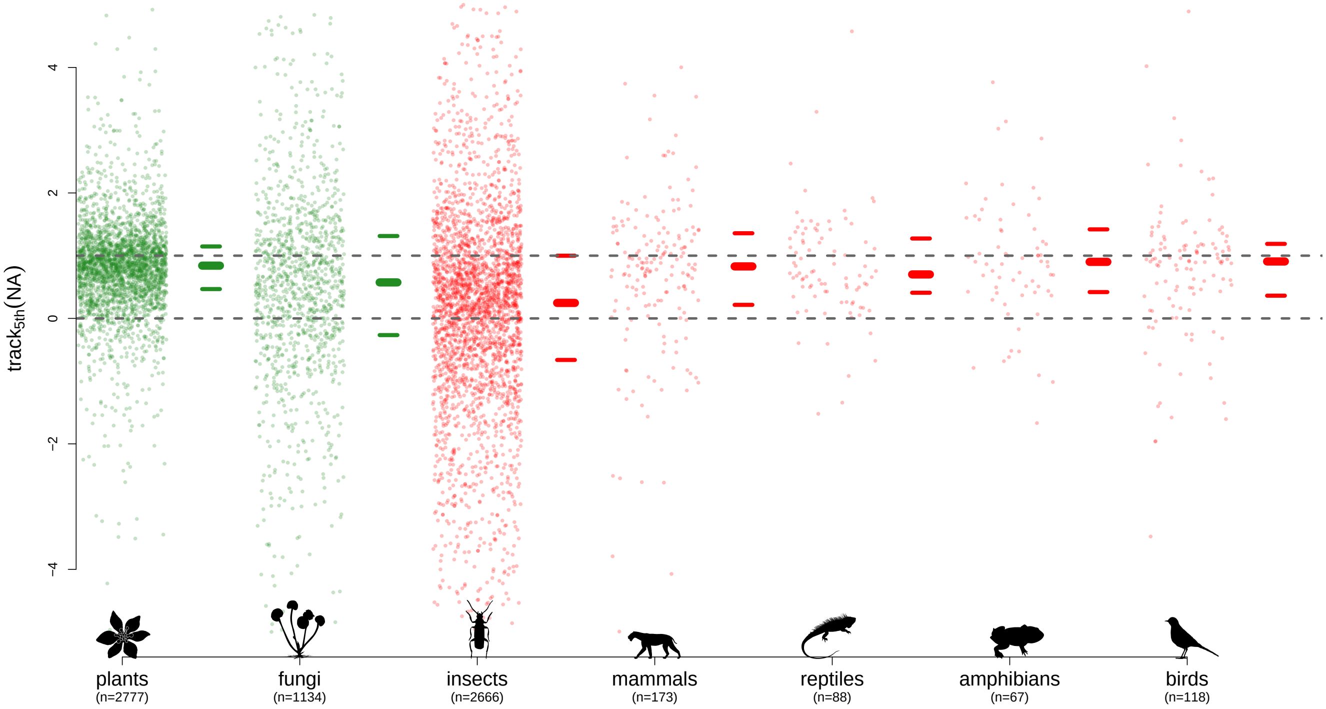


**Figure 41: Rainday counts, 5th quantile.** Legend as figure 1 in main text.

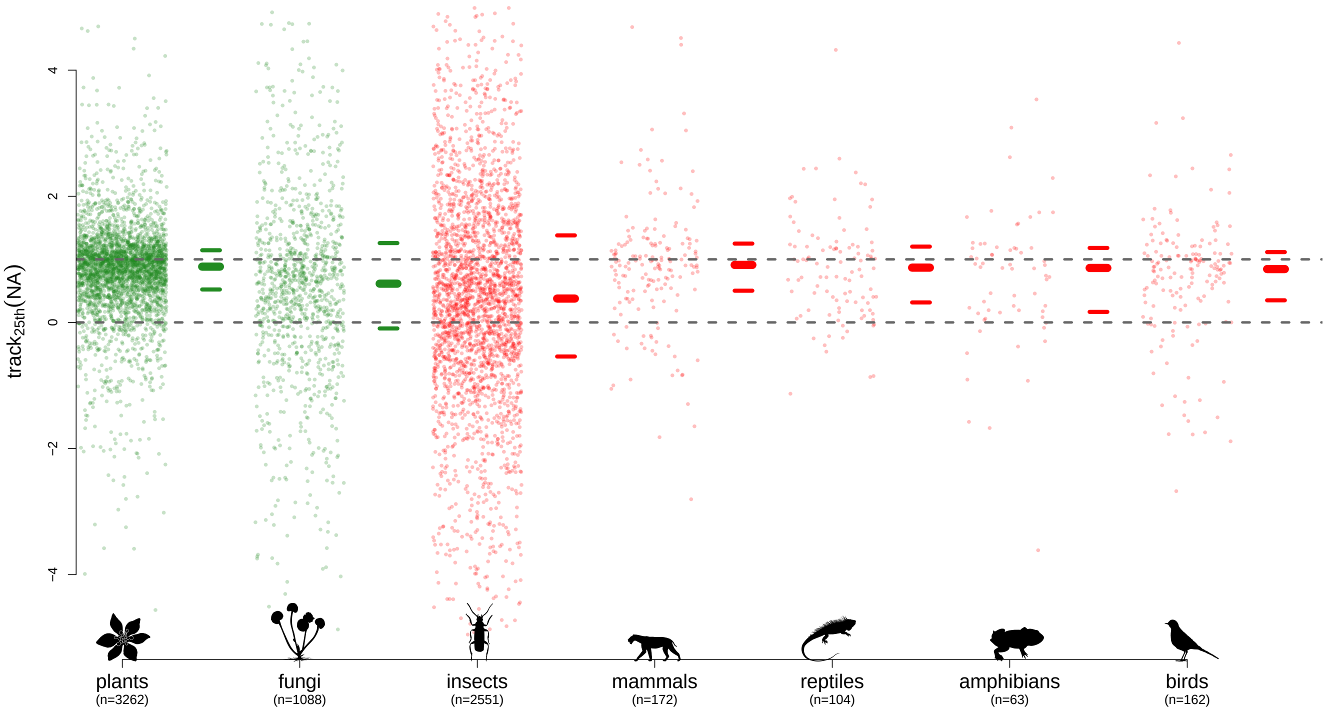


**Figure 42: Rainday counts, 25th quantile.** Legend as figure 1 in main text.

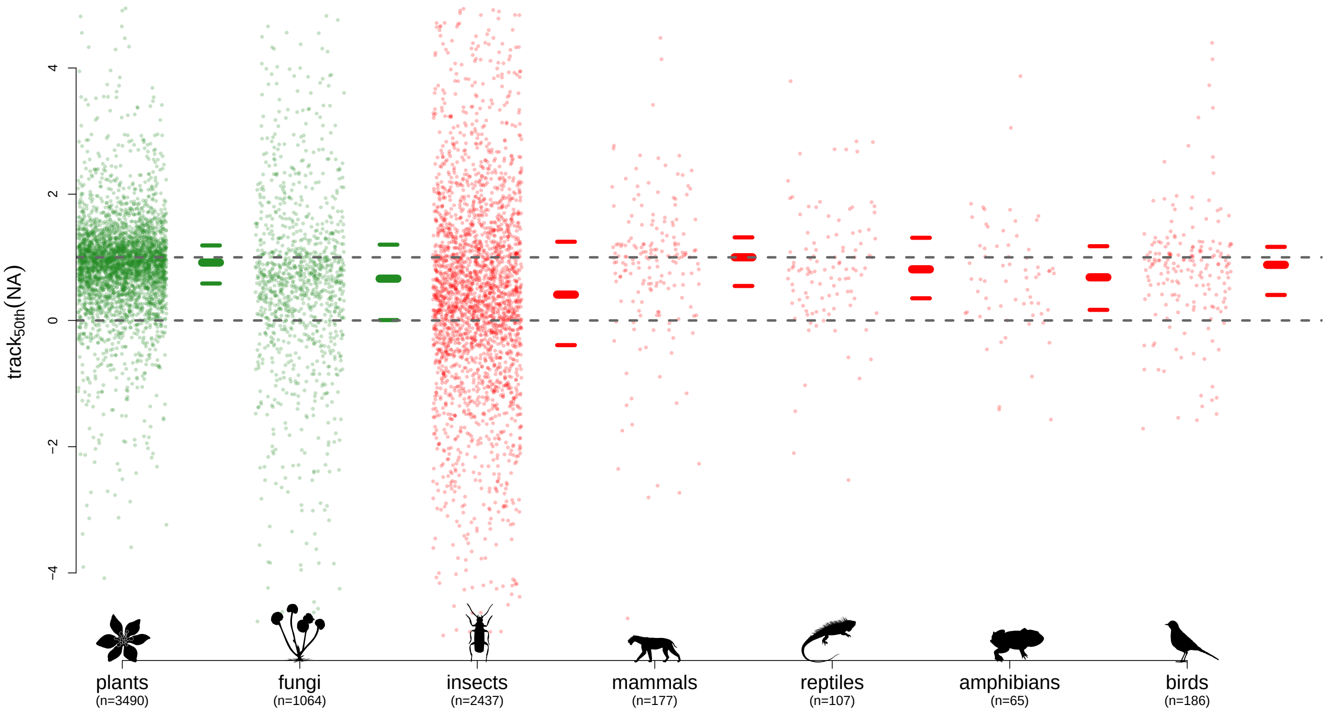


**Figure 43: Rainday counts, 50th quantile (median).** Legend as figure 1 in main text.

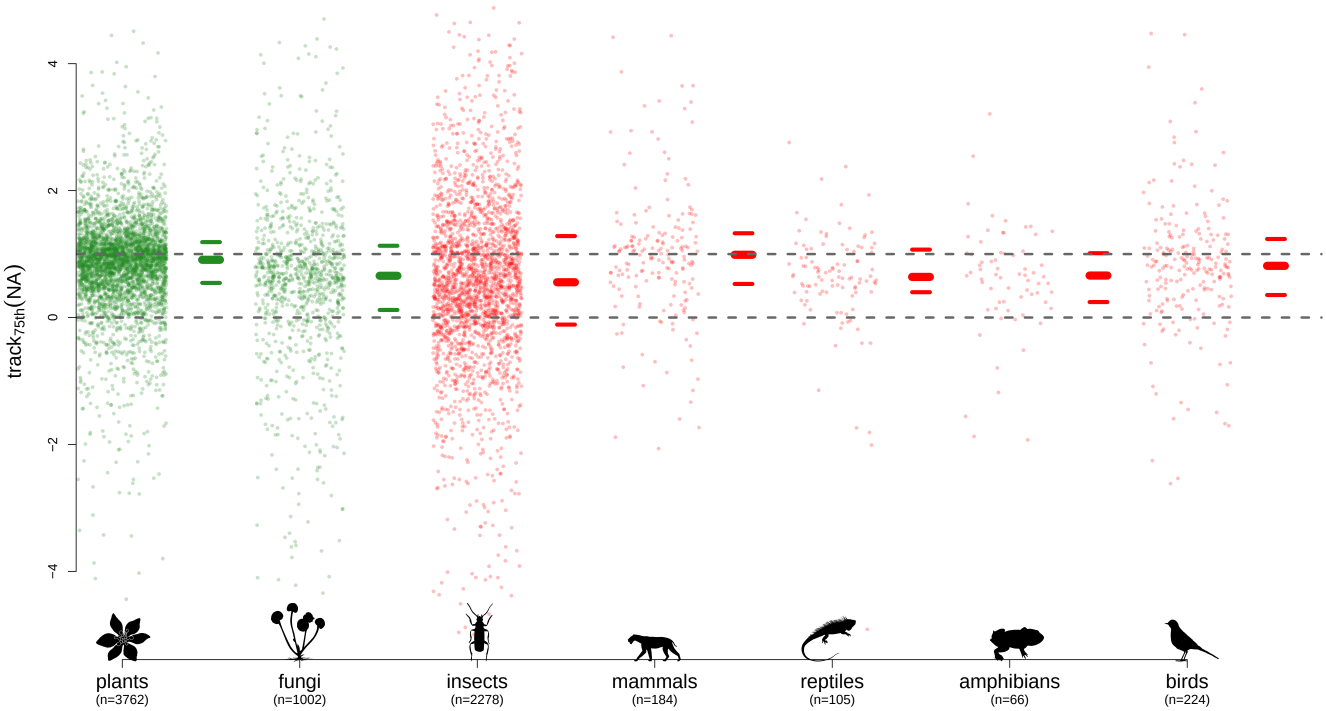


**Figure 44: Rainday counts, 75th quantile.** Legend as figure 1 in main text.

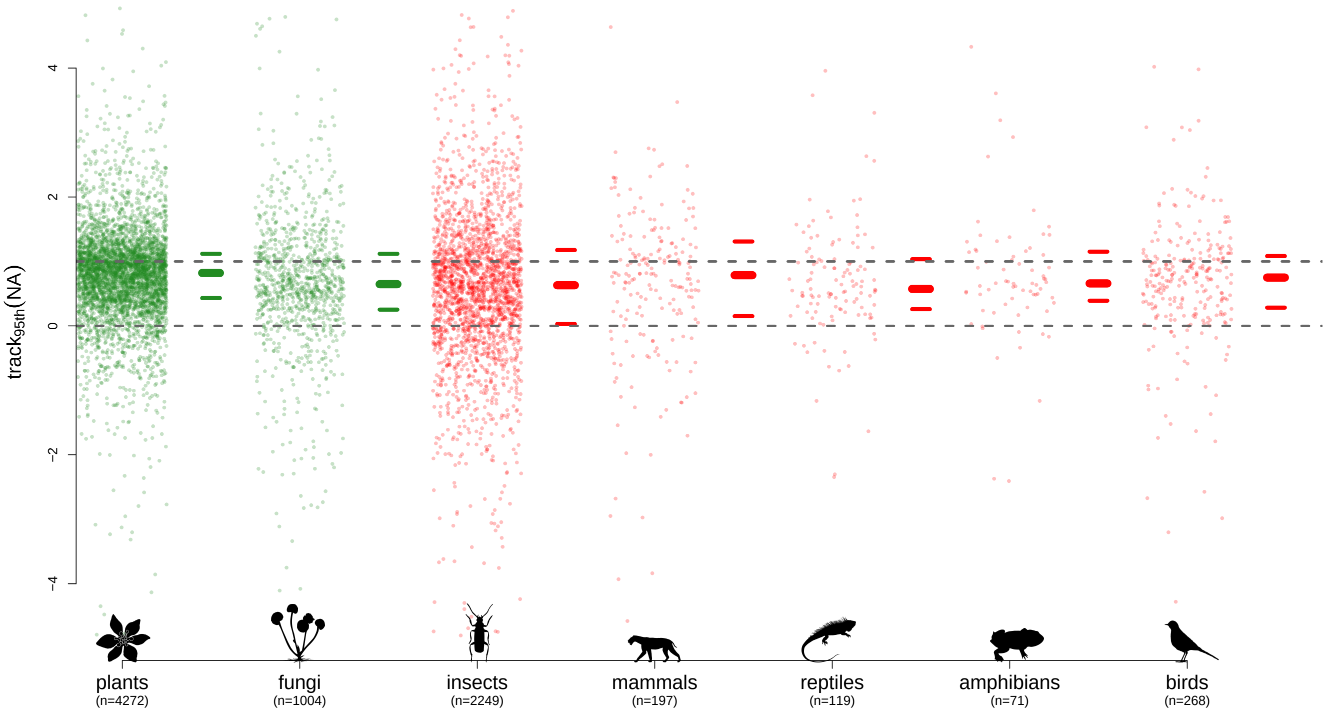


**Figure 45: Rainday counts, 95th quantile.** Legend as figure 1 in main text.

### 2 Trait correlations and phylogenetic signal (*c.f.* figure 3)

Please note that, since figure 2 in the main text is an exemplar of the data underlying the phylogenetic signals shown in these figures, we do not replicate it here across all data-types as it is superfluous.

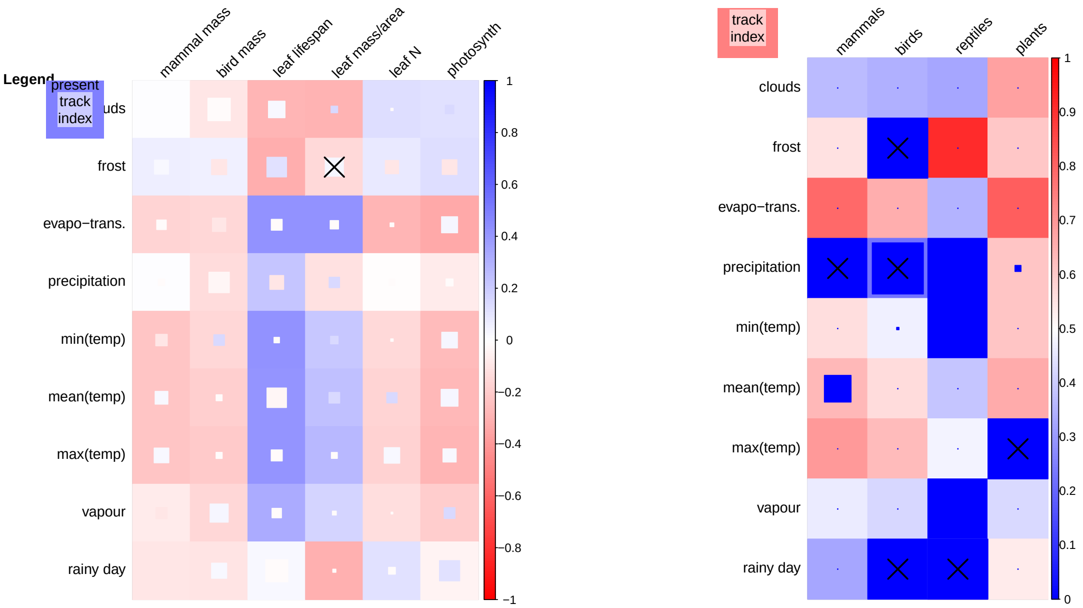


**(a)** Trait correlations **(b)** Phylogenetic signal

**Figure 46: Trait correlations and phylogenetic signals of the 5th quantiles.** Legend as figure 3 in the main text.

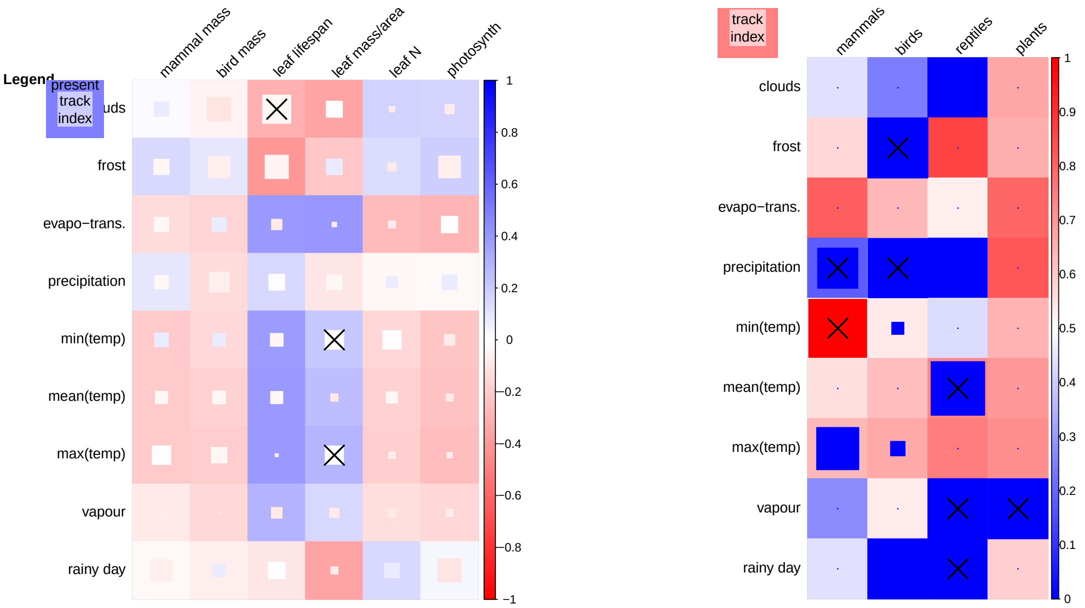


**(a)** Trait correlations

**(b)** Phylogenetic signal

**Figure 47: Trait correlations and phylogenetic signals of the 25th quantiles.** Legend as figure 3 in the main text.

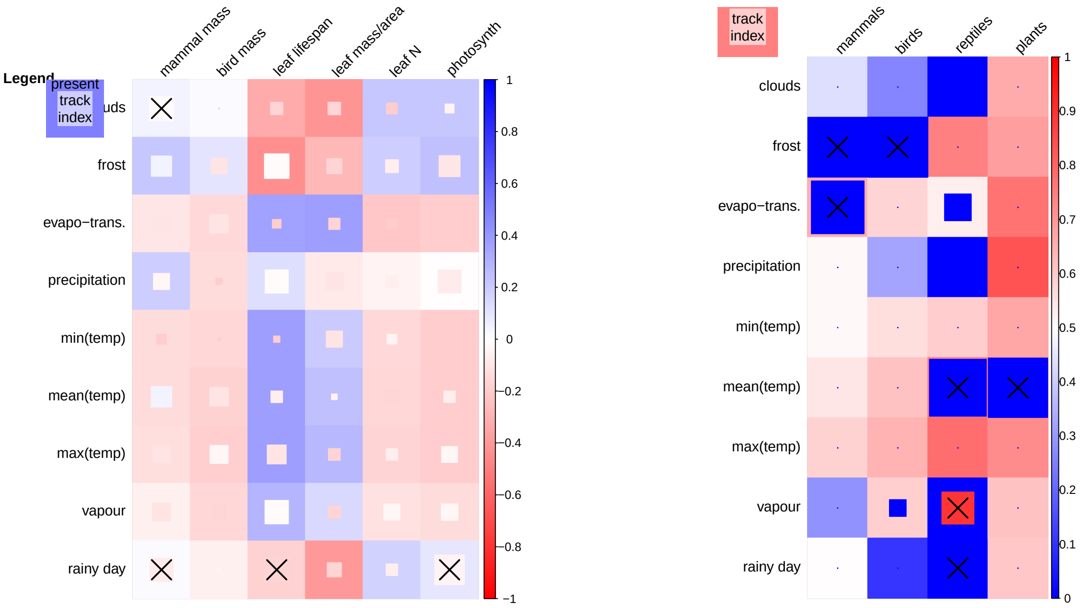


**(a)** Trait correlations

**(b)** Phylogenetic signal

**Figure 48: Trait correlations and phylogenetic signals of the 50th quantiles (medians).** Legend as figure 3 in the main text.

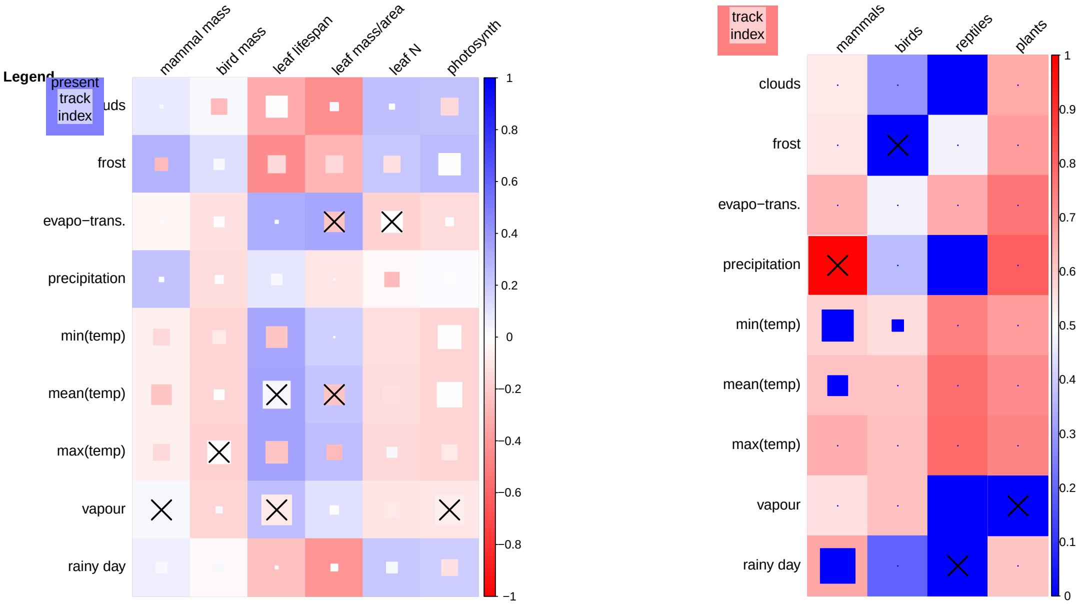


**(a)** Trait correlations

**(b)** Phylogenetic signal

**Figure 49: Trait correlations and phylogenetic signals of the 75th quantiles.** Legend as figure 3 in the main text.

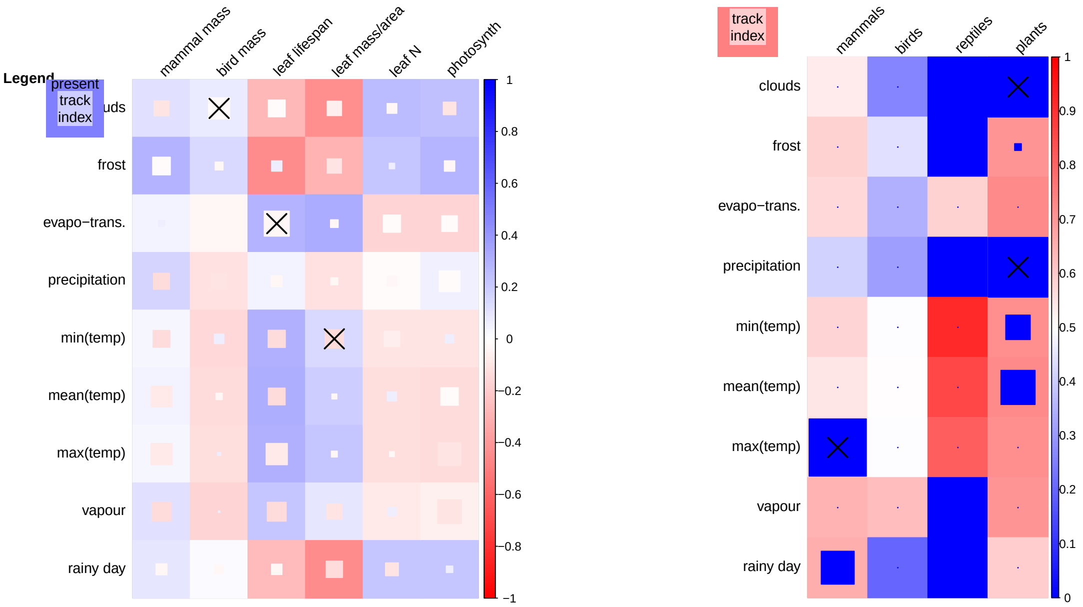


**(a)** Trait correlations

**(b)** Phylogenetic signal

**Figure 50: Trait correlations and phylogenetic signals of the 95th quantiles.** Legend as figure 3 in the main text.

### 3 Principal components analyses (*c.f.* figure 4)

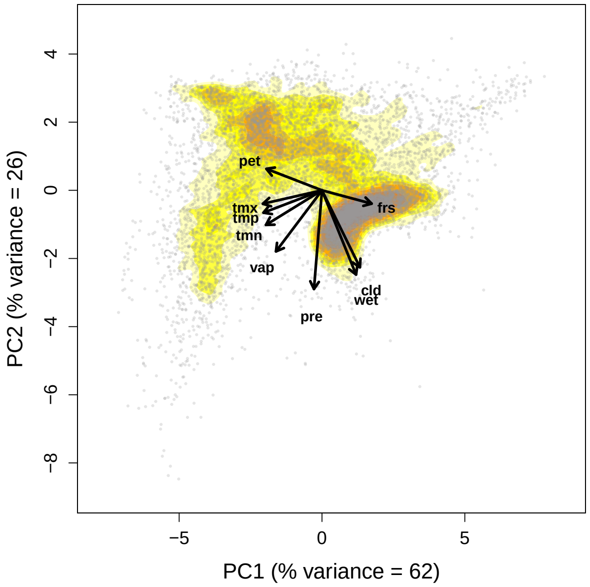


**(a)** Past climate space (1955–1980)

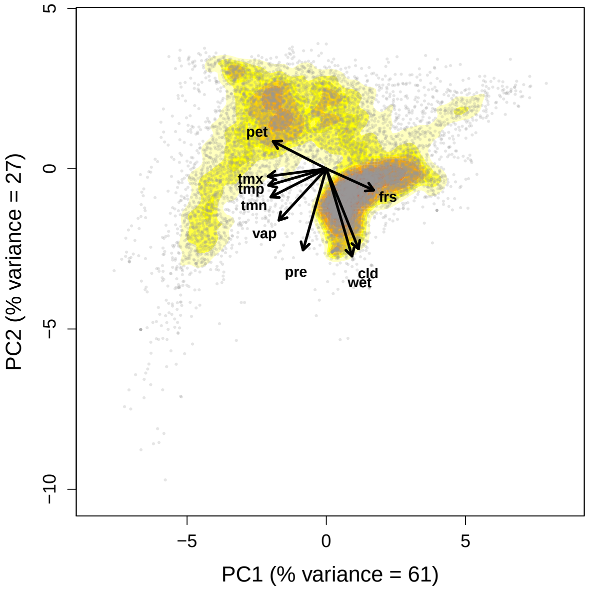


**(b)** Present climate space (1990–2015)

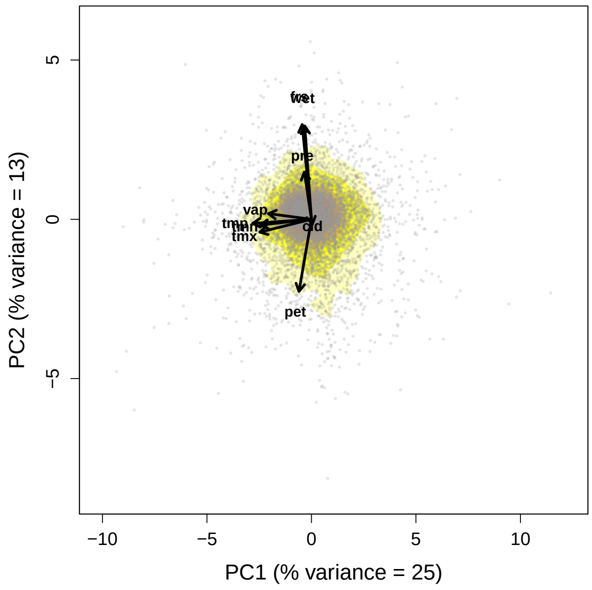


**(c)** Tracking climate space

**Figure 51: Principal components analyses for the 5th quantiles.** Legend as figure 4 in the main text.

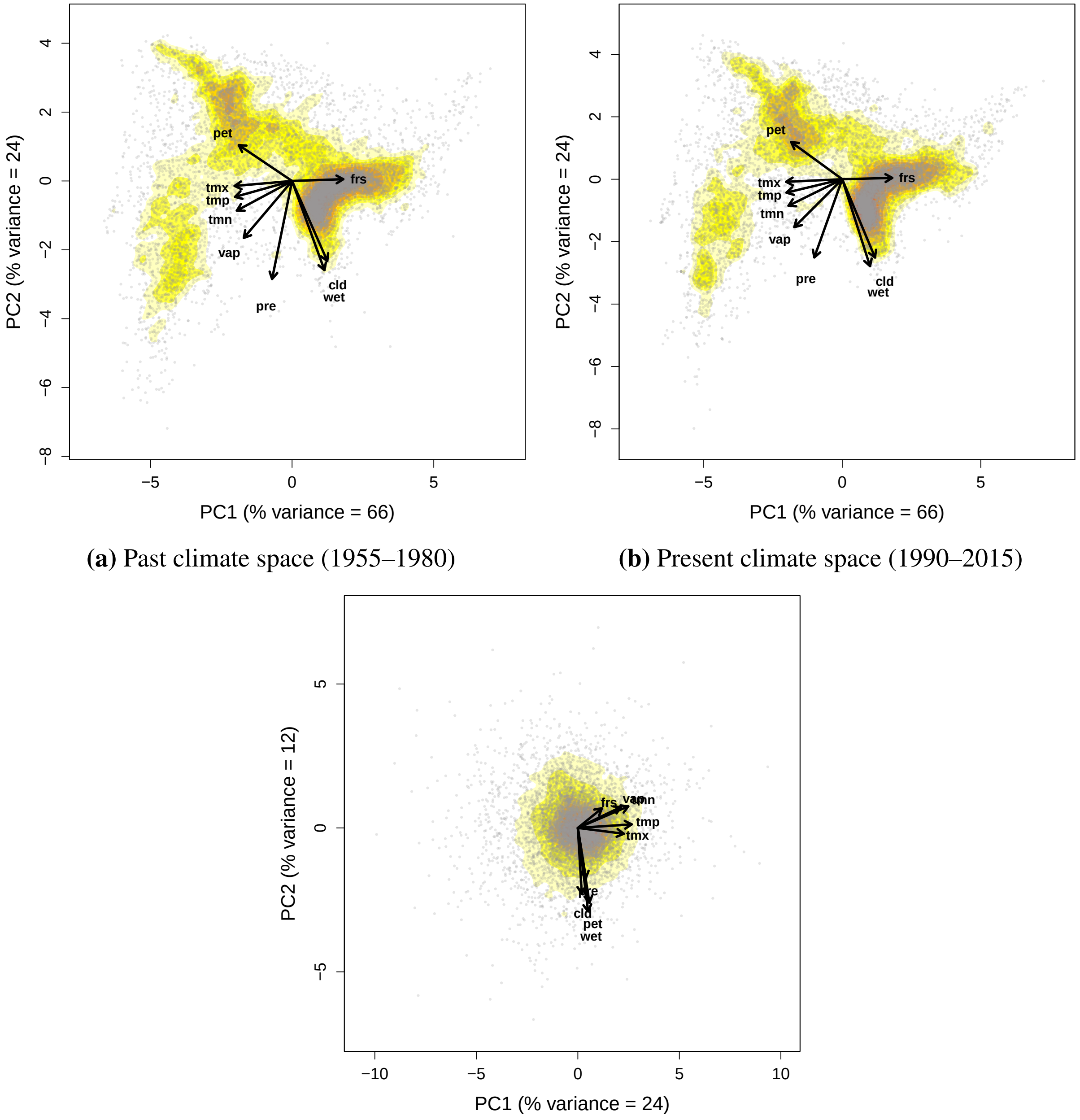


**(a)** Past climate space (1955–1980)

**(b)** Present climate space (1990–2015)

**(c)** Tracking climate space

**Figure 52: Principal components analyses for the 25th quantiles.** Legend as figure 4 in the main text.

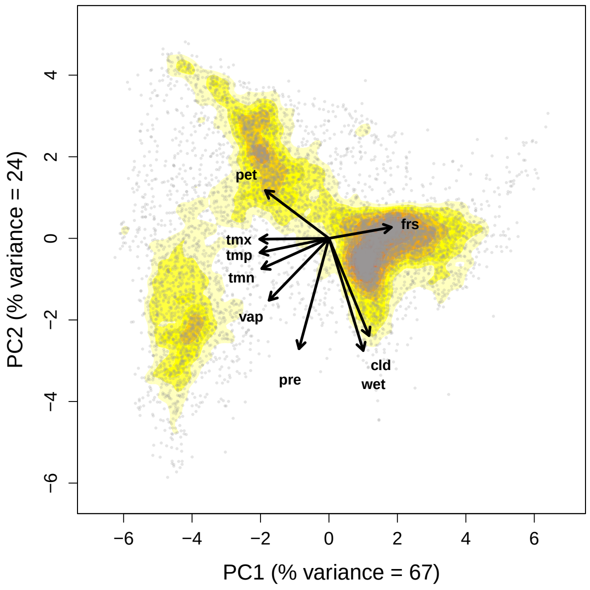


**(a)** Past climate space (1955–1980)

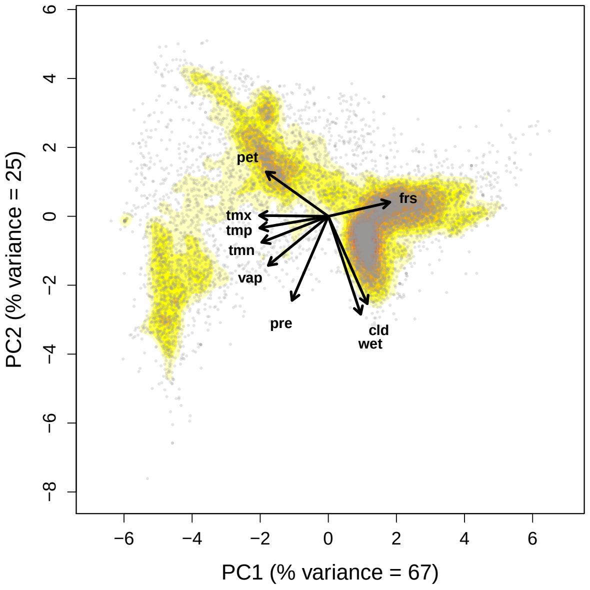


**(b)** Present climate space (1990–2015)

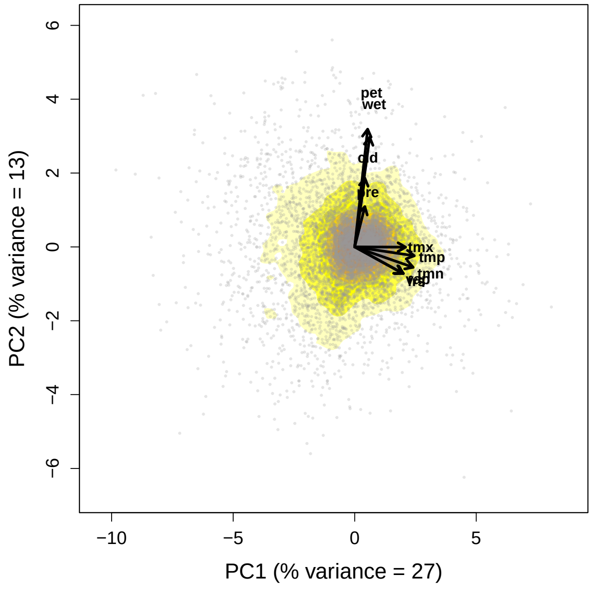


**(c)** Tracking climate space

**Figure 53: Principal components analyses for the 50th quantiles (*i.e.*, medians).** Legend as figure 4 in the main text.

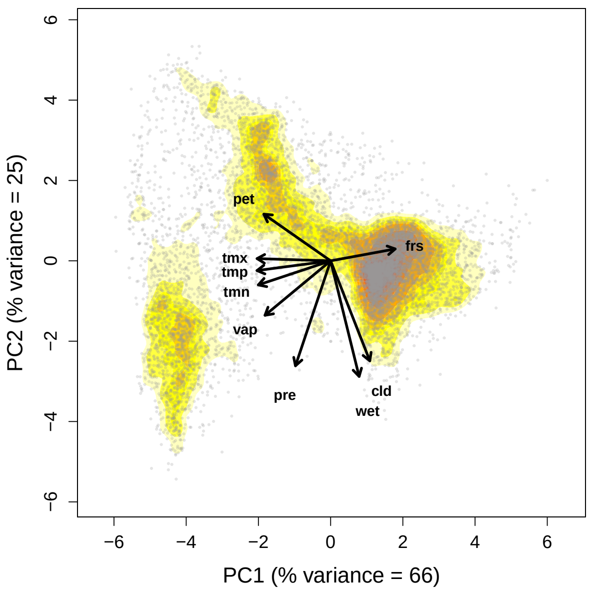


**(a)** Past climate space (1955–1980)

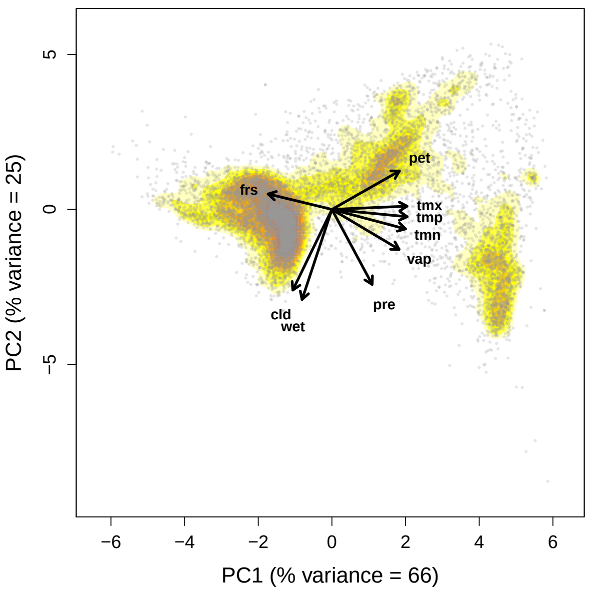


**(b)** Present climate space (1990–2015)

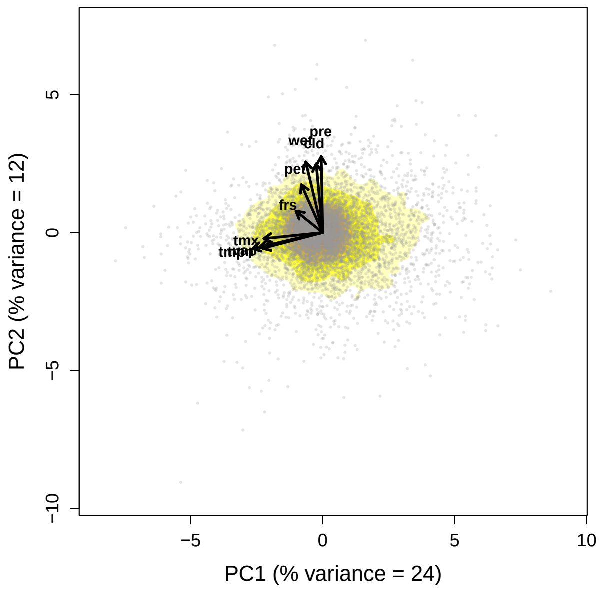


**(c)** Tracking climate space

**Figure 54: Principal components analyses for the 75th quantiles.** Legend as figure 4 in the main text.

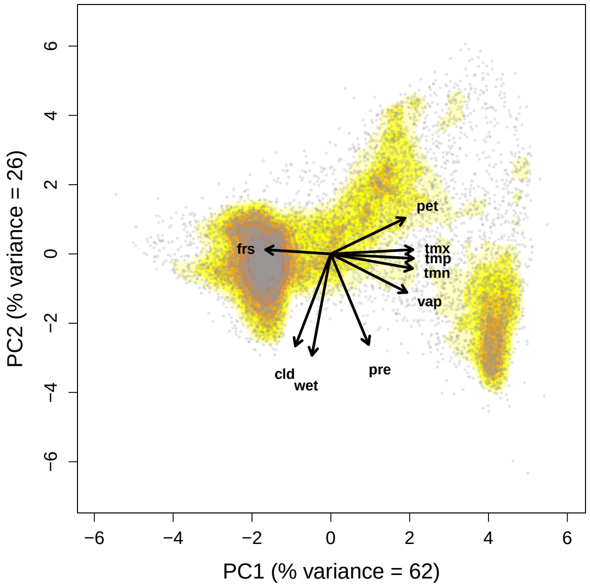


**(a)** Past climate space (1955–1980)

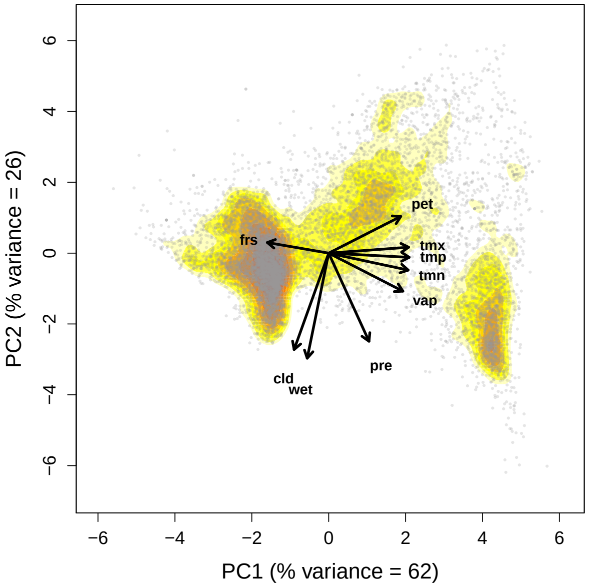


**(b)** Present climate space (1990–2015)

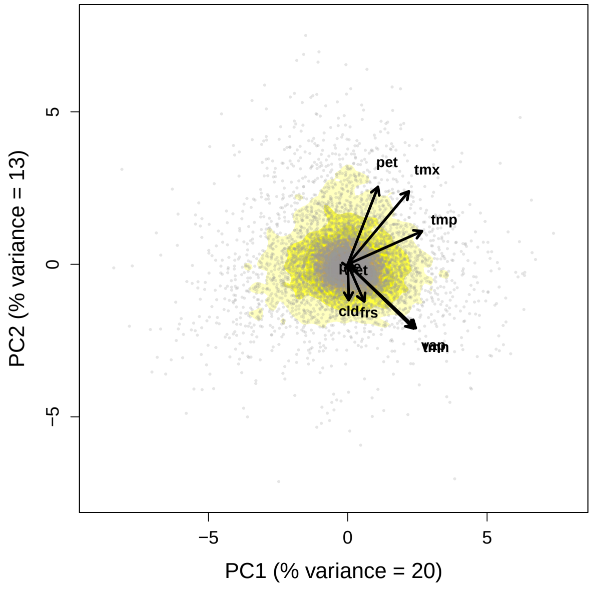


**(c)** Tracking climate space

**Figure 55: Principal components analyses for the 95th quantiles.** Legend as figure 4 in the main text.
