## supplemental code for "The realised velocity of climate change reveals remarkable idiosyncrasy of species’ distributional shifts": ms.tex

\documentclass[12pt]{report}
\usepackage[utf8]{inputenc}
\usepackage{times}
\renewcommand\thesection{\arabic{section}}
\usepackage[parfill]{parskip}
\usepackage{subcaption}
\usepackage{graphicx}
\usepackage[left=0.75in,right=0.75in,top=0.75in,bottom=0.75in]{geometry}
\usepackage{multirow}
\usepackage{hhline}
\usepackage{nth}
\usepackage{bm}
\usepackage{siunitx}
\usepackage[labelfont=bf]{caption}
\usepackage[doublespacing]{setspace}
\usepackage{filecontents}
\usepackage{lineno} \linenumbers
\usepackage{amsmath}
\usepackage{pdflscape}

%%%%%%%%%%%%%%%%%%%%%%%%%%%%%%%%%%%
%Bibliography%%%%%%%%%%%%%%%%%%%%%%
%%%%%%%%%%%%%%%%%%%%%%%%%%%%%%%%%%%
\usepackage[style=numeric-comp,sorting=none,uniquelist=minyear,uniquename=false,maxcitenames=2,maxbibnames=5,doi=false,url=false,isbn=false,defernumbers=true,giveninits=true]{biblatex}
%Italicise et al.
\renewbibmacro*{name:andothers}{
% Based on name:andothers from biblatex.def
\ifboolexpr{ test{\ifnumequal{\value{listcount}}{\value{liststop}}} and test \ifmorenames } {\ifnumgreater{\value{liststop}}{1} {\finalandcomma}
{}%
\andothersdelim\bibstring[\emph]{andothers}} {}}
%Make 'and' an ampersand
\renewcommand*{\finalnamedelim}{%
\ifnumgreater{\value{liststop}}{2}{\finalandcomma}{}%
\addspace\&\space}
%Remove 'in:'
\renewbibmacro{in:}{}
%Remove pp. on pages numbers
\DeclareFieldFormat[article]{pages}{#1}%
%Remove unwanted entries
\AtEveryBibitem{%
  \clearfield{day}%
  \clearfield{month}%
  \clearfield{endday}%
  \clearfield{endmonth}%
  \clearfield{url}%
  \clearfield{URL}%
  \clearfield{eprint}%
}
%Remove spaces between initials
%Remove double quotes around titles
\DeclareFieldFormat*{citetitle}{#1}
\DeclareFieldFormat*{title}{#1}
\addbibresource{/home/will/Papers/Library.bib}
\begin{document}

% To-do before bioRxiv:
% ! p-values in lambda

\section*{Title page}
\textbf{Article title}: The realised velocity of climate change
reveals remarkable idiosyncrasy of species' distributional shifts

\textbf{Authors:} William D.\ Pearse$^{1,*}$ \& T.\ Jonathan
Davies$^{2,*}$

$^1$ Department of Biology \& Ecology Center, Utah State University,
5305 Old Main Hill, Logan UT, 84322

$^2$ Department of Biological Sciences, University of British
Columbia, Vanouver, Canada

$^*$ To whom correspondence should be addressed:
\emph{} and \emph{}

\textbf{Word-count}: 2634 (excluding figure legends)

\clearpage
\textbf{To date, our understanding of how species have shifted in
  response to recent climate warming has been based on a few studies
  with a limited number of species. Here we present a comprehensive,
  global overview of species' distributional responses to changing
  climate across a broad variety of taxa (animals, plants, and
  fungi). We characterise species' responses using a metric that
  describes the realised velocity of climate change: how closely
  species' responses have tracked changing climate through time. In
  contrast to existing `climate velocity' metrics that have focused on
  space, we focus on species and index their responses to a null
  expectation of change in order to examine drivers of inter-specific
  variation. Here we show that species are tracking climate on
  average, but not sufficiently to keep up with the pace of climate
  change. Further, species responses are highly idiosyncratic, and
  thus highlight that projections assuming uniform responses may be
  misleading.  This is in stark contrast to species' present-day and
  historical climate niches, which show strong evidence of the imprint
  of evolutionary history and functional traits. Our analyses are a
  first step in exploring the vast wealth of empirical data on
  species' historic responses to recent climate change.}

The natural environment is changing rapidly. Over the last century,
habitats have become increasingly fragmented and
isolated\supercite{Haddad2015}, the climate has become warmer, and
extreme climate events more frequent\supercite{IPCC2014}. Biological
diversity---the species of plants and animals with which we cohabit
the Earth---has responded predictably. Present-day extinction rates
are estimated to be up to three orders of magnitude greater than
background rates \supercite{Pimm2014} and are projected to increase
further over the next several decades \supercite{MEA2005}. Current
estimates suggest that over one million species may be threatened with
extinction\supercite{IPBESland2019}, and that we have experienced a
50\% decline in animal diversity in the past 40 years
\supercite{McLellan2014}.

The main direct human-induced drivers that impact biodiversity now are
habitat loss \supercite{Newbold2015} and fragmentation
\supercite{Haddad2015}, but climate change is likely to become a
dominant driver in the next few
decades\supercite{Parmesan2003,Thuiller2007,Urban2015}. Projected
impacts of climate change on biodiversity have attracted much
attention, but the uncertainty around the magnitude of future
extinctions (\emph{e.g.}, 7.9\%\supercite{Urban2015} vs.\
37\%\supercite{Thomas2004} of species) highlights the inadequacy of
current forecasts.  In the face of changing climate species must
either shift in space, to track favourable conditions, or time, for
example, flowering and breeding earlier.
% I've dropped the migration reference here because that's shifting in
% both space and time, and so sort of ruins the either/or structure
Evidence suggests they are doing both
\supercite{Parmesan2003,Parmesan2006,Menzel2006,Chen2011}, but not all
species are responding equally. Some species are expanding their
distributions, sometimes invasively \supercite{Hellmann2008}, and some
species are flowering later\supercite{Cook2012}; species may also be
simply becoming less predictable \supercite{Pearse2017phenology},
perhaps as evolved responses to environmental cues break down.  Simply
quantifying species' mean responses ignores the important variability
in responses among species \supercite{Wolkovich2012}. Developing
robust predictions of ecological responses to climate change thus
requires data from diverse species, ecosystems, and climates to
account for this large inter-specific variation.

The velocity of temperature change---the rate a species would have to
shift its range to maintain constant temperature---has been calculated
at an average of 0.42 km/yr \supercite{Loarie2009}. A meta-analysis of
taxonomic groups estimated the average pole-ward shift in species'
distributions has been approximately 1.69 km/yr \supercite{Chen2011},
suggesting that species might be capable of keeping pace with shifting
climate. Yet even this estimate is derived from just 21 studies with
limited taxonomic coverage (361 birds, 9 mammals, and no plants), and
critically any single point estimate does not reflect the large
variation in responses among species or within taxa. Here we leverage
the vast volume of distributional data stored in digital data
repositories\supercite{url_gbif}, representing over 10,000 species of
plants, fungi, mammals, birds, reptiles, amphibians, and insects, to
provide the first global synthesis of species distributional responses
to climate change. By contrasting historical niche dimensions with
those of the present day, we can characterise the realised biotic
velocity of climate change, which represents the relative degree to
which species track climate across space.

We show that, on average, species' historical (pre-1980) ranges have
warmed significantly, and that species have generally shifted their
distributions to ameliorate this warming, but not sufficiently to keep
pace with climate change. Yet this general pattern masks a remarkably
degree of variability in species responses, and many species' ranges
are now even warmer than if their distributions had remained
static. Further, we find that, while species' niche dimensions are
strongly predicted by their evolutionary history and functional
traits, the same is not true of their degree of tracking. Thus while
species' distributions are reasonably predictable, their changes in
distributions are not.

\clearpage
\subsection*{Results and discussion}
\textbf{A comparative index of species' niche tracking.} We propose a
measure that captures the extent to which a species' distribution is
tracking climate by standardising its observed niche change by the
change it would have experienced had it not shifted its
distribution. Formally, we define this as:

\begin{equation}
  track_\tau = \frac{current_\tau - past_\tau}{projected_\tau - past_\tau}
\end{equation}

Where $current_\tau$ is a given quantile ($\tau$) of a species'
climatic distribution across its present-day range, $past_\tau$ the
same quantile of a species' range in the past, and $projected_\tau$
the same quantile of a species' past range under present-day climatic
conditions. This index should be calculated separately for
temperature, pollution, or whatever environmental variable is of
interest. Thus a $track$ of 0 would indicate a species that has
tracked changing climate perfectly, while a $track$ of 1 would be
consistent with a species that had not moved whatsoever to track
changing environmental conditions. Values between 0 and 1 suggest
moderate tracking, while values greater than 1 or less than 0
represent a change in climate relationship greater than if the species
had not moved at all. Uncertainty in either species' past or current
ranges can be accounted for through a bootstrap procedure, whereby the
past and current distributions are re-sampled to standardise sampling
across time periods (see methods).

Our approach has two main innovations: its focus on (1) the whole of a
species' distribution and (2) expected change. (1) When estimating
species' range distributions we typically focus on species'
limits\supercite{Sexton2009} (\emph{e.g.}, upper and lower quantiles)
while in community ecological \supercite{Tilman2004} and
macro-evolutionary \supercite{Zanne2018} studies we tend to focus on
central tendencies (\emph{e.g.}, the median). However, there is no
strong \emph{a priori} reason that species can, or must, track all
quantiles equally. Indeed some evidence suggests different constraints
on species' warm-range limits versus their cool-range limits, with
biotic interactions more influential in the former, and climate
determining the latter\supercite{Freeman2018}.  It is possible,
therefore, that the drivers of species' range edges and centres may be
diverging under climate change. Our approach is to check all quantiles
of a species' distribution. (2) The magnitude of experienced and
projected climate change shows substantial geographical variation
\supercite{Loarie2009}, and so species' responses might thus be
expected to also vary in space. We suggest that species'
distributional responses should be measured on the basis of the
magnitude of change to which they are exposed, and the denominator of
our index re-scales species' change around an expectation under
distributional stasis.  This change reflects the magnitude of
ecological and evolutionary selection, and while there are many ways
of addressing this, we suggest ours is perhaps the simplest and most
transparent, so represents a fair starting point. We note that we
could also rescale our index according to the distance that would need
to be travelled to maintain climate stasis (\emph{sensu}
\citeauthor{Loarie2009}\supercite{Loarie2009}), but prefer to present
here a unitless index of tracking, which is simpler to compare across
taxa and between biogeographic regions.

\textbf{Niche tracking across the tree of life throughout the past
  sixty years.} To assess the extent to which species have tracked
climate, we used the most comprehensively sampled time-series data for
species' distributions (GBIF\supercite{url_gbif}) and climate (UEA's
CRU\supercite{Harris2014}). This represents over 630 million raw
occurrence records collected over the past century. We focus on five
quantiles: the \nth{5}, \nth{25}, \nth{50} (median), \nth{75}, and
\nth{95}. Here, we present species' range data collected from
1955--2015 for 10,700 species (4,879 plants, 1,517 fungi, 273 mammals,
433 birds, 147 reptiles and amphibians, and 3,451 insects), each
observation matched to one of nine climate variables in the year in
which they were sampled (mean, minimum, maximum temperature,
precipitation, `rainy-day' frequency, vapour pressure, potential
evapotranspiration, `frost day' frequency, and cloud cover). In the
methods we give more detail of our reproducible analysis pipeline, and
in the supplementary materials provide instructions for repeating our
analyses with different data curation and cleaning choices. Our metric
was designed to account for known biases in the GBIF data
\supercite{Beck2014}, and in the methods we outline our simulations
which address the influence of sampling uncertainty.

Almost all species are tracking some climate variable through time
(99.5\% have at least one $track_\tau$ between 0 and 1). Most species
are, to some extent, tracking temperature (see Figure \ref{overview});
65\% track at least one quantile and 43\% track at least two.
% Giving an index in units of distance is a good idea; I think it
% needs some thought, though, because our index is unitless because we
% divide the units through; I'm not sure the number we get out of this
% would be directly interpretable like theirs.
The three temperature indices are, however, the least-tracked of any
climate variable, while precipitation is the most-tracked (86\% of
species track at least one of its quantiles). Notably, however, many
taxa are not tracking climate through time, and many have experienced
more change than if their distributions had remained static. Just as
most species are tracking at least one quantile of one climate
variable, 99.9\% of species have `overshoot' (have a $track_\tau$ less
than 0; for example becoming colder in a warming region) in at least
one quantile of at least one climate variable. As we show in Figure
\ref{overview}, these results are consistent with our null simulations
where species' range change is a combination of essentially random
movement and a moderate degree of subsequent climate filtering. Thus
while our results confirm that, by-and-large, species are tracking
climate change, the magnitude of this varies across species, taxa, and
aspects of climate (see Figure \ref{overview}). For example, while
fungi and insects are, on the whole, broadly tracking the median of
temperature (median $track_{50\%}$ of 0.46 and 0.64, respectively),
plants and mammals track temperature (median $track_{50\%}$ of 1.16
and 1.07, respectively) more poorly than precipitation (median
$track_{5\%}$ of 0.79 and 0.89, respectively).

\textbf{Idiosyncratic drivers of species' tracking.} To examine the
potential drivers of species' ranges and range tracking, we first
examined the phylogenetic signal of species' climate quantiles. We
focused on mammals, birds, reptiles, and plants, since we were able to
find broadly inclusive, dated phylogenies for these taxa. We found
strong evidence of phylogenetic signal in species' past (median
Pagel's $\lambda$\supercite{Pagel1999} of 0.60) and current (median
$\lambda$ of 0.57) distributions, but limited signal in our index
(median $\lambda$ of $<$0.001). Past climate showed stronger
phylogenetic signal than our index in 158 of all 180 (4 taxa, 5
quantiles, and 9 climate variables) possible taxon-climate-quantile
combinations (144 for current climate; see Figures \ref{phylo} and
\ref{traits-signal}). This confirms the known result that species'
current-day and past ranges are strongly influenced by
evolution\supercite{Wiens2010}, but suggests that species' realised
velocity of change (our tracking index) may be associated with
different factors.

We additionally examined species' traits as potential drivers of
species' distributions and potential to track changing environment. We
focused on mammals, birds, and plants, since global, openly-released
trait datasets exist for these taxa. Again, we found statistically
significant associations between traits and past distributions of 191
of 270 (at $\alpha_{5\%}$; 6 traits, 9 climate variables, and 5
quantiles) taxon-climate-quantile combinations (193 for current
distributions), but not between the traits and our tracking index (13
combinations or 5\%; see Figure \ref{traits-signal}). While the traits
we used are strongly associated with species' climate
responses\supercite{Wright2004,McCain2014} (hence their strong
predictive power of species' distributions, but not degree of
tracking), we emphasise that we could not find sufficient data to
assess the potential impact of dispersal
ability\supercite{Schloss2012}. These findings, while surprising, are
in keeping with emerging evidence that species' traits are a
relatively poor predictor of change at range edges
\supercite{Angert2011}.

\textbf{Species' movement through climate space is disconnected from
  their position within it.} Given our index is poorly predicted by
traditionally important variables, we now consider whether index
values are predictive of each other. It is well known that the axes of
global climate are not independent and that they are not changing
independently\supercite{Harris2014,IPCC2014}, and so we would expect
species' relative tracking of climate to show similar patterns. As we
show in Figure \ref{pca}, principal components analysis of species'
current and past climate distributions, which we refer to as their
climate space, shows strong correlations across climate variables. Yet
our index does not; species' relative degree of tracking is both of
much higher dimension (the amount of variance explained by each
principal component axis is similar) and shows relatively less
correlation across climates axes . We argue, therefore, that while
species' positions in climate space show strong associations, species'
relative tracking of their position in climate space does not. This
result explains the poor predictive power of species' traits and
evolutionary history for our index: the ecological and evolutionary
rules that determine species' climate relationships are
changing. Thus, in our analysis, we can explain species' current
distributions only by the degree to which they resemble past
distributions.

There are alternative explanations for our results, of which we
highlight three: (1) we are ignoring important niche axes, (2) we are
ignoring biotic interactions, and (3) our index highlights random
variation in distribution. (1) Our climate variables are not
all-encompassing, and we ignore other drivers such as geology and
habitat-type. Yet we do detect strong pattern in current and past
distributions, suggesting that the absence of pattern in our indices
is meaningful. (2) Competition and facilitation among species are
likely strong drivers of species' ability to track
climate\supercite{Davis1999}. Yet, again, they are surely also likely
to have driven species' past distributions, and so it is unclear to us
why only our metric, and not also current and past distributions, is
so poorly predicted by our trait and phylogenetic data. (3) If species
were perfectly tracking climate, $current_\tau$ would be equal to
$past_\tau$ and so our index values could reflect sampling error or
some other purely stochastic input. Yet such noise, if truly random,
would mean our empirical index values would be centred at 0 or
resemble our simulated data, neither or which is the case (see Figure
\ref{overview}).

\clearpage
\subsection*{Conclusion}
Our results are consistent with the processes underlying species'
responses to climate change being distinct from those that generated
their distributions in the past. This is in keeping with emerging
evidence from the paleo-ecological literature that species' historic
ranges were more variable than previously thought, and seemingly
influenced by different environmental factors than today
\supercite{Veloz2012,Maguire2015}. During and after the last ice-age,
for example, `no-analogue' conditions generated previously-unseen
species assemblages \supercite{Williams2007}. Equally, our observation
of idiosyncrasy in species' degree of tracking is consistent with
species' population crashes (and so extinctions) becoming less
predictable during previous mass extinction events
\supercite{Levinton2001,Jablonski2004}. We call upon others to examine
the additional drivers of our metric of relative change, and to extend
its definition to include more nuanced definitions of climate
space. We predict, however, that if the current mass extinction event
continues, it is likely that patterns of idiosyncrasy among species'
declines and distributional changes will become more common.

\clearpage
% Figures
\clearpage
\begin{figure}[h!]
  \begin{center}
    \begin{subfigure}{.7\textwidth}
      \includegraphics[width=\textwidth]{../figures/violin-tmp-0-5.pdf}
      \caption{$track_{50\%}$(mean-temp)}
    \end{subfigure}
    \\
    \begin{subfigure}{.7\textwidth}
      \includegraphics[width=\textwidth]{../figures/violin-pre-0-05.pdf}
      \caption{$track_{5\%}$(precipitation)}
    \end{subfigure}
  \end{center}
  \caption{\textbf{Most species are tracking some aspect of climate,
      but there is pronounced variation among species and clades.} In
    (a) we show the \nth{50} quantile (median) of our tracking index
    of mean annual temperature, and in (b) the \nth{5} quantile of our
    index of precipitation. On the left of each plot, our seven major
    taxonomic groups plotted (in green for plants and fungi, and in
    red for animals). At the far-right of (b), we show the output from
    our simulations of species that are perfectly tracking,
    imperfectly tracking, or not tracking climate at all (in blue; see
    Methods for details). To the right of each distribution of points
    the median (thickest line) and inter-quartile range (thinner
    lines) are shown as horizontal lines for each distribution. While
    the majority of species are tracking global climate to some
    extent, there is is profound variation among species and
    clades. Figure \ref{overview}a makes clear the variation among and
    within taxonomic groups; the distributions of fungi, insects, and
    reptiles are, broadly, tracking changes in mean temperature well,
    while other taxa (\emph{e.g.}, mammals and plants) are
    not. Comparing Figures \ref{overview}a and \ref{overview}b makes
    clear how different taxonomic groups and species may be tracking
    different aspects of climate; plants are broadly tracking the
    lower-limit of precipitation, for instance.}
  \label{overview}
\end{figure}

\clearpage
\begin{landscape}
\begin{figure}[h!]
  \begin{center}
    \includegraphics[width=1.4\textwidth]{../figures/phylo-tmp-50.pdf}
  \end{center}
  \caption{\textbf{Despite strong phylogenetic signal in species'
      static distributions, species' changes in distributions are not
      phylogenetically patterned.} Here we show the plant tree of
    life, with the two oldest branches shortened for compactness
    (emphasised with dashed lines). Internal branches are shaded by
    ancestral state reconstructions, assuming Brownian motion, of
    species' median mean-temperature in the past. Phylogeny tips are
    coloured according to the track index of the same underlying data
    (compare with Figure \ref{overview}). This general pattern of
    strong phylogenetic signal of climate (shown by the coloured
    pattern to the internal branches), but not of degree of tracking,
    is characteristic of almost all of our underlying climate data
    (see also Figure \ref{traits-signal}).}
  \label{phylo}
\end{figure}
\end{landscape}

\clearpage
\begin{figure}[h!]
  \begin{center}
    \begin{subfigure}{.45\textwidth}
      \includegraphics[width=\textwidth]{../figures/traits-50.pdf}
      \caption{Trait correlations (Pearson's $\rho$)}
    \end{subfigure}
    \begin{subfigure}{.45\textwidth}
      \includegraphics[width=\textwidth]{../figures/physig-5.pdf}
      \caption{Phylogenetic signal (Pagel's $\lambda$)}
    \end{subfigure}
  \end{center}
  \caption{\textbf{Species' static climate variables show strong trait
      correlations and phylogenetic signal, but their relative degree
      of tracking does not.} In (a), we show the correlation
    (Pearson's $\rho$) between the median of each climate variable
    (rows) with each of our included traits (columns) using colour
    (see legend at the right). In each cell, the outer, larger box
    depicts the correlation between species' present-day climate and
    the trait, while the smaller, inner box the correlation for
    species' relative tracking (whose size is proportional to
    intensity of correlation). The 4 (of 54) track-index correlations
    that are statistically significant (at $\alpha_{5\%}$) are
    highlighted with crosses. In (b), we show, for each taxon, the
    phylogenetic signal (Pagel's $\lambda$\supercite{Pagel1999}) of
    major climate variables and relative tracking indices (rows) using
    colour (see legend at the right). As with (a), in each cell the
    outer, larger box depicts the correlation between species'
    present-day climate and the trait, while the smaller, inner box
    the correlation for species' relative tracking (whose size is
    proportional to phylogenetic signal strength). The single
    track-index metric for which phylogenetic signal was stronger than
    the underlying data is highlighted with a cross.}
  \label{traits-signal}
\end{figure}
% To-do: find an alternative to the crosses

\clearpage
\begin{figure}[h!]
  \begin{center}
    \begin{subfigure}{.45\textwidth}
      \includegraphics[width=\textwidth]{../figures/pca-05-past.pdf}
      \caption{Past climate space (1955--1980)}
    \end{subfigure}
    \begin{subfigure}{.45\textwidth}
      \includegraphics[width=\textwidth]{../figures/pca-05-present.pdf}
      \caption{Present climate space (1990--2015)}
    \end{subfigure}
    \begin{subfigure}{.45\textwidth}
      \includegraphics[width=\textwidth]{../figures/pca-05-index.pdf}
      \caption{Tracking climate space}
    \end{subfigure}
  \end{center}
  % Note: bad filenames I know, but these are the 50th entries
  \caption{\textbf{Species' movement through climate space differs
      from their current and past positions within it.}  Principal
    component analysis (PCA) biplots of (a) species' median past
    climate values, (b) median current climate, and (c) the median
    track-index of all climate variables. Each PCA axis explains some
    amount of variation in the underlying data (labelled on the axes
    themselves). In (a) and (b), species' climate variables are
    correlated with one-another, and the space reflects global
    patterns in climate. In (c), this pattern has broken down: while
    some associations among climate axes still remain (notably
    temperature), the general pattern is one of uncorrelated shifts
    through climate space.}
  \label{pca}
\end{figure}

\clearpage

\textbf{Acknowledgements} WDP was funded by NSF ABI-1759965, NSF
EF-1802605, and USDA Forest Service agreement 18-CS-11046000-041. TJD
was funded by \emph{Fonds de Recherche Nature et Technologies} grant
number 168004. The images in Figure \ref{overview} are taken, with
gratitude, from \emph{http://phylopic.org/}; all are under the Public
Domain Dedication 1.0 license and uncredited unless otherwise stated
below. The reptile image was drawn by Jack Mayer Wood, the plant by
Jason McNair, the insect by Birgit Lang, the mammal by Scott Hartman
(used under the Creative Commons Attribution 3.0 Unported license),
and the amphibian by Nobu Tamura (used under the Creative Commons
Attribution 3.0 Unported license).

\textbf{Author Contributions} WDP and TJD contributed to all aspects
of the study.

\textbf{Author information} The authors declare no competing financial
interests. Correspondence and requests for materials should be
addressed to \emph{} and \emph{}.

\textbf{Competing interests} The authors declare no competing
financial interests

\clearpage

\section*{\Large Methods}
\renewcommand{\figurename}{Extended Data Figure}
\renewcommand{\tablename}{Extended Data Table}
  
All analyses were conducted in \emph{R} version \emph{3.6.1}
\supercite{R2018}, and all names in \emph{italics} refer to \emph{R}
packages. Code to reproduce all analyses in their entirety is
available in the online supplementary materials and online at
\emph{https://github.com/willpearse/track-index}.  In the
supplementary materials, we provide extended figures summarising all
tracking index quantiles and climate variables in the same detail as
presented in the main text.

\subsection*{Data collation}

Species' distribution data were downloaded from the Global
Biodiversity Information Facility (GBIF)
\supercite{url_gbif,url_gbif_occurrences}. We downloaded only data
highlighted by GBIF as not having any spatial issues, meaning they
contained no obvious location errors (\emph{e.g.}, latitude/longitude
swaps or rounded coordinates). These occurrence data include records
that were flagged as having potential issues and corrected by GBIF;
our results were qualitatively identical if we excluded these records
and so we include them here for completeness. We used `human
observation' occurrences (\emph{e.g.}, records from surveys), and not
vouchered specimens (\emph{e.g.}, pressed specimens in a herbarium)
because preliminary analysis revealed qualitatively identical results
across the two methods, but with many fewer observations vouchered
observations.

We only worked with species that had at least 1000 records on GBIF
within our focal time periods (1955--1980 and 1985--2015). We focused
on these time periods because they contain sufficient observations for
large-scale analysis. Since climate change was underway within the
1980s \supercite{IPCC2014}, this also allows us to have reasonably
large (if na\"{i}vely defined) `pre-change' and `post-change'
periods. Our 1000 observation threshold limits us to better-studied
species, but our results are robust to increases (\emph{e.g.}, 10,000
observations) in this threshold. We only made use of data for which
species identity was known, and we ignored variation among
sub-species. The GBIF taxonomy is, itself, to some extent checked and
harmonised, and so we retained in our analysis any species within GBIF
that was not present in another datasets (see
\citeauthor{Cornwell2019}\supercite{Cornwell2019} for a discussion of
the GBIF taxonomy for plants). In the supplementary materials, we give
instructions for re-running our analysis with differing curation
thresholds to verify our choices.

Climate data were downloaded from
\citeauthor{Harris2014}\supercite{Harris2014}'s nine global
time-series. We split the data into yearly (averaged) 0.5$^\circ$
cells, from which we estimated each of our nine climate variables for
each species observation using
\emph{raster}\supercite{Hijmans2019}. Plant trait data were taken from
\citeauthor{Wright2004}\supercite{Wright2004} (leaf lifespan, leaf
mass-per-unit-area, leaf Nitrogen mass, and leaf photosynthetic
capacity), mammal body mass data from the Amniote Trait
Database\supercite{Myhrvold2015}, and bird body mass data from
EltonTraits \supercite{Wilman2014}. We were unable to find sufficient
coverage of the other taxa in this study in open-access trait
databases to facilitate further analysis. We used existing global
mammal\supercite{Faurby2015}, bird\supercite{Jetz2012}, amphibian
\supercite{Jetz2018}, plant (`ALLOTB' phylogeny
\supercite{Smith2018}), and reptiles\supercite{Zheng2016} phylogenies,
and in cases where posterior distributions of trees were available we
used a single draw from that distribution. Missing species were added
into the phylogenies using \emph{congeneric.merge} in
\emph{pez}\supercite{Pearse2015}.

\subsection*{Calculating and testing our track index}
As we describe in the main text, we quantify the realised velocity of
climate change using a simple index of climate tracking. Our index,
$track_\tau$, scales the observed magnitude of species' shift in
climate ($current_\tau - past_\tau$) according to the degree of change
to which that species was exposed ($projected_\tau - past_\tau$). Our
metric is comparable across species, and can also be calculated in
such a way as to control for changes in sampling. The sampling of
species through time and across space is known to be uneven across the
data in GBIF \supercite{Beck2013,Beck2014}. To ensure our method was
statistically robust to such changes, we employed a bootstrap
procedure during the calculation of $track_\tau$. To do this we
randomly re-sampled, with replacement, the occurrences making up
$current_\tau$ and $projected_\tau$ to be of the same number as
$past_\tau$, and vice-versa for $past_\tau$ and $current_\tau$. We
repeated this process 999 times, and calculated a $track_\tau$ value
for each re-sample, generating 999 $track_\tau$ values whose medians
we report here. This process accounts for uneven sampling by
generating a pseudo-posterior distribution of conservatively-estimated
values. In each bootstrap, the better-sampled time period is
sub-sampled to match the poorer-sampled period, accounting for
differences in sampling. Equally, the poorer-sampled period is
re-sampled in order to increase the variance around the index,
essentially cross-validating our estimate. When applied to empirical
data, our bootstrap approach reveals that the certainty differing
tracking quantiles (and climate variables) is uneven. We therefore
excluded uncertain (\emph{i.e.}, estimates with high variation across
pseudo-posteriors) and outliers from figures within the main text, and
so report sample sizes within Figure \ref{overview} (and its
counterparts in the supplementary materials).

To assess the performance of our (bootstrapped) track index, we
simulated species' with ranges that tracked climate to varying extents
through time. Using the climate data from above in the years 1962 and
2002 (the mid-points of the ranges of our data), we simulated species
with varying maximum possible range sizes (2x2, 5x5, 10x10, or 20x20
grid-cell extents) and varying latitudinal shifts in range through
time (-4, -3, ..., 3, or 4 grid-cells). We also varied species' degree
of environmental tracking ($\alpha$; taking a value of 0, 0.5, or 1)
and probability of occupancy($\sigma$; taking a value of 0.5, .75, or
1). Together, these latter two parameters define a species'
probability of being present in a particular cell within its range:
$(1-\alpha) \times sigma + \alpha \times \mathcal{N}$.  $\mathcal{N}$
is defined as a scaled Normal probability density function with a mean
equal to the median of the species' distribution in 1962 and a
variance of 1. Thus a species with an $\alpha$ of 1 is present only in
cells within its range that resemble its climatic centre (\emph{i.e.},
are similar to the median value of the species' climatic
distribution), and a species with an $\alpha$ of 0 is present in
proportion $\sigma$ of the cells within its potential range. We
simulated all possible combinations of parameters across 100 possible
centre-points of species' ranges, providing a full range of potential
range shifts and 26,598 total simulation runs. As with our empirical
data, in simulation runs where, by chance, simulated species were not
present in the past or current time-period, they were excluded from
the analysis.

\subsection*{Trait, phylogenetic, and PCA analyses}

We calculated the Pearson's correlation between all estimated climatic
niche variables ($current_\tau$, $past_\tau$, $projected_\tau$,
$track_\tau$ and the bootstrapped $track_\tau$). While we show only a
handful of these correlations in Figure \ref{traits-signal}, we
provide all correlations in the supplementary figures (and note that
the qualitative results of all correlations are the same). For the
phylogenetic signal analyses, we likewise calculated Pagel's $\lambda$
(using \emph{phytools}\supercite{Revell2012phytools}) for all indices,
reporting all results in full in the supplementary materials (results
are, again, qualitatively identical to the results in the main
text). Principal component analyses were performed on all current,
past, and track indices across all variables for a given quantile
(\emph{i.e.}, the \nth{5}, \nth{25}, \nth{50}, \nth{75}, and
\nth{95}). While we report only the \nth{50} quantiles in the main
text, in the supplementary materials we give results for all
quantiles, which are qualitatively identical to the results in the
main text.

\printbibliography

\end{document}


%%% Local Variables:
%%% mode: latex
%%% TeX-master: t
%%% End:
