## supplemental code for "The realised velocity of climate change reveals remarkable idiosyncrasy of species’ distributional shifts": supplement.tex

\documentclass[12pt]{report}
\usepackage[utf8]{inputenc}
\usepackage{times}
\renewcommand\thesection{\arabic{section}}
\usepackage[parfill]{parskip}
\usepackage{subcaption}
\usepackage{graphicx}
\usepackage[left=0.75in,right=0.75in,top=0.75in,bottom=0.75in]{geometry}
\usepackage{multirow}
\usepackage{hhline}
\usepackage{nth}
\usepackage{bm}
\usepackage{siunitx}
\usepackage[labelfont=bf]{caption}
\usepackage[doublespacing]{setspace}
\usepackage{filecontents}
\usepackage{lineno} \linenumbers
\usepackage{amsmath}
\usepackage{pdflscape}
\usepackage{url}

\begin{document}

\section*{Supplementary materials}
\textbf{Article title}: The realised velocity of climate change
reveals remarkable idiosyncrasy of species' distributional shifts

\textbf{Authors:} William D.\ Pearse$^{1,*}$ \& T.\ Jonathan
Davies$^{2,*}$

$^1$ Department of Biology \& Ecology Center, Utah State University,
5305 Old Main Hill, Logan UT, 84322

$^2$ Department of Biological Sciences, University of British
Columbia, Vanouver, Canada

$^*$ To whom correspondence should be addressed:
\emph{} and \emph{}

\clearpage

Our supplementary materials are organised according to the figures in
the main text. We replicate the figures (and so analyses) calculated
for the subsets of the data shown there across the entire range of the
data below. Figure legend are terse and perfunctory because the
interpretation of them can be done with reference to the figures in
the main text (which are references in the legends of the
supplementary figures).

\clearpage
\section{Overall plots of tracking index values (\emph{c.f.} figure 1)}
\subsection{Cloud cover}
\begin{figure}[h!]
  \begin{center}
    \includegraphics[width=\textwidth]{../figures/violin-cld-0-05.pdf}
  \end{center}
  \caption{\textbf{Cloud cover, \nth{5} quantile.} Legend as
    figure 1 in main text.}
\end{figure}
\begin{figure}[h!]
  \begin{center}
    \includegraphics[width=\textwidth]{../figures/violin-cld-0-25.pdf}
  \end{center}
  \caption{\textbf{Cloud cover, \nth{25} quantile.} Legend as
    figure 1 in main text.}
\end{figure}
\begin{figure}[h!]
  \begin{center}
    \includegraphics[width=\textwidth]{../figures/violin-cld-0-5.pdf}
  \end{center}
  \caption{\textbf{Cloud cover, \nth{50} quantile (median).}
    Legend as figure 1 in main text.}
\end{figure}
\begin{figure}[h!]
  \begin{center}
    \includegraphics[width=\textwidth]{../figures/violin-cld-0-75.pdf}
  \end{center}
  \caption{\textbf{Cloud cover, \nth{75} quantile.} Legend as
    figure 1 in main text.}
\end{figure}
\begin{figure}[h!]
  \begin{center}
    \includegraphics[width=\textwidth]{../figures/violin-cld-0-95.pdf}
  \end{center}
  \caption{\textbf{Cloud cover, \nth{95} quantile.} Legend as
    figure 1 in main text.}
\end{figure}

\clearpage
\subsection{Frost days}
\begin{figure}[h!]
  \begin{center}
    \includegraphics[width=\textwidth]{../figures/violin-frs-0-05.pdf}
  \end{center}
  \caption{\textbf{Frost days, \nth{5} quantile.} Legend as
    figure 1 in main text.}
\end{figure}
\begin{figure}[h!]
  \begin{center}
    \includegraphics[width=\textwidth]{../figures/violin-frs-0-25.pdf}
  \end{center}
  \caption{\textbf{Frost days, \nth{25} quantile.} Legend as
    figure 1 in main text.}
\end{figure}
\begin{figure}[h!]
  \begin{center}
    \includegraphics[width=\textwidth]{../figures/violin-frs-0-5.pdf}
  \end{center}
  \caption{\textbf{Frost days, \nth{50} quantile (median).}
    Legend as figure 1 in main text.}
\end{figure}
\begin{figure}[h!]
  \begin{center}
    \includegraphics[width=\textwidth]{../figures/violin-frs-0-75.pdf}
  \end{center}
  \caption{\textbf{Frost days, \nth{75} quantile.} Legend as
    figure 1 in main text.}
\end{figure}
\begin{figure}[h!]
  \begin{center}
    \includegraphics[width=\textwidth]{../figures/violin-frs-0-95.pdf}
  \end{center}
  \caption{\textbf{Frost days, \nth{95} quantile.} Legend as
    figure 1 in main text.}
\end{figure}

\clearpage
\subsection{Potential evapotranspiration}
\begin{figure}[h!]
  \begin{center}
    \includegraphics[width=\textwidth]{../figures/violin-pet-0-05.pdf}
  \end{center}
  \caption{\textbf{Potential evapotranspiration, \nth{5} quantile.} Legend as
    figure 1 in main text.}
\end{figure}
\begin{figure}[h!]
  \begin{center}
    \includegraphics[width=\textwidth]{../figures/violin-pet-0-25.pdf}
  \end{center}
  \caption{\textbf{Potential evapotranspiration, \nth{25} quantile.} Legend as
    figure 1 in main text.}
\end{figure}
\begin{figure}[h!]
  \begin{center}
    \includegraphics[width=\textwidth]{../figures/violin-pet-0-5.pdf}
  \end{center}
  \caption{\textbf{Potential evapotranspiration, \nth{50} quantile (median).}
    Legend as figure 1 in main text.}
\end{figure}
\begin{figure}[h!]
  \begin{center}
    \includegraphics[width=\textwidth]{../figures/violin-pet-0-75.pdf}
  \end{center}
  \caption{\textbf{Potential evapotranspiration, \nth{75} quantile.} Legend as
    figure 1 in main text.}
\end{figure}
\begin{figure}[h!]
  \begin{center}
    \includegraphics[width=\textwidth]{../figures/violin-pet-0-95.pdf}
  \end{center}
  \caption{\textbf{Potential evapotranspiration, \nth{95} quantile.} Legend as
    figure 1 in main text.}
\end{figure}

\clearpage
\subsection{Precipitation}
\begin{figure}[h!]
  \begin{center}
    \includegraphics[width=\textwidth]{../figures/violin-pre-0-05.pdf}
  \end{center}
  \caption{\textbf{Precipitation, \nth{5} quantile.} Legend as
    figure 1 in main text.}
\end{figure}
\begin{figure}[h!]
  \begin{center}
    \includegraphics[width=\textwidth]{../figures/violin-pre-0-25.pdf}
  \end{center}
  \caption{\textbf{Precipitation, \nth{25} quantile.} Legend as
    figure 1 in main text.}
\end{figure}
\begin{figure}[h!]
  \begin{center}
    \includegraphics[width=\textwidth]{../figures/violin-pre-0-5.pdf}
  \end{center}
  \caption{\textbf{Precipitation, \nth{50} quantile (median).}
    Legend as figure 1 in main text.}
\end{figure}
\begin{figure}[h!]
  \begin{center}
    \includegraphics[width=\textwidth]{../figures/violin-pre-0-75.pdf}
  \end{center}
  \caption{\textbf{Precipitation, \nth{75} quantile.} Legend as
    figure 1 in main text.}
\end{figure}
\begin{figure}[h!]
  \begin{center}
    \includegraphics[width=\textwidth]{../figures/violin-pre-0-95.pdf}
  \end{center}
  \caption{\textbf{Precipitation, \nth{95} quantile.} Legend as
    figure 1 in main text.}
\end{figure}

\clearpage
\subsection{Minimum temperature}
\begin{figure}[h!]
  \begin{center}
    \includegraphics[width=\textwidth]{../figures/violin-tmn-0-05.pdf}
  \end{center}
  \caption{\textbf{Minimum temperature, \nth{5} quantile.} Legend as
    figure 1 in main text.}
\end{figure}
\begin{figure}[h!]
  \begin{center}
    \includegraphics[width=\textwidth]{../figures/violin-tmn-0-25.pdf}
  \end{center}
  \caption{\textbf{Minimum temperature, \nth{25} quantile.} Legend as
    figure 1 in main text.}
\end{figure}
\begin{figure}[h!]
  \begin{center}
    \includegraphics[width=\textwidth]{../figures/violin-tmn-0-5.pdf}
  \end{center}
  \caption{\textbf{Minimum temperature, \nth{50} quantile (median).}
    Legend as figure 1 in main text.}
\end{figure}
\begin{figure}[h!]
  \begin{center}
    \includegraphics[width=\textwidth]{../figures/violin-tmn-0-75.pdf}
  \end{center}
  \caption{\textbf{Minimum temperature, \nth{75} quantile.} Legend as
    figure 1 in main text.}
\end{figure}
\begin{figure}[h!]
  \begin{center}
    \includegraphics[width=\textwidth]{../figures/violin-tmn-0-95.pdf}
  \end{center}
  \caption{\textbf{Minimum temperature, \nth{95} quantile.} Legend as
    figure 1 in main text.}
\end{figure}

\clearpage
\subsection{Mean temperature}
\begin{figure}[h!]
  \begin{center}
    \includegraphics[width=\textwidth]{../figures/violin-tmp-0-05.pdf}
  \end{center}
  \caption{\textbf{Mean-temperature, \nth{5} quantile.} Legend as
    figure 1 in main text.}
\end{figure}
\begin{figure}[h!]
  \begin{center}
    \includegraphics[width=\textwidth]{../figures/violin-tmp-0-25.pdf}
  \end{center}
  \caption{\textbf{Mean-temperature, \nth{25} quantile.} Legend as
    figure 1 in main text.}
\end{figure}
\begin{figure}[h!]
  \begin{center}
    \includegraphics[width=\textwidth]{../figures/violin-tmp-0-5.pdf}
  \end{center}
  \caption{\textbf{Mean-temperature, \nth{50} quantile (median).}
    Legend as figure 1 in main text.}
\end{figure}
\begin{figure}[h!]
  \begin{center}
    \includegraphics[width=\textwidth]{../figures/violin-tmp-0-75.pdf}
  \end{center}
  \caption{\textbf{Mean-temperature, \nth{75} quantile.} Legend as
    figure 1 in main text.}
\end{figure}
\begin{figure}[h!]
  \begin{center}
    \includegraphics[width=\textwidth]{../figures/violin-tmp-0-95.pdf}
  \end{center}
  \caption{\textbf{Mean-temperature, \nth{95} quantile.} Legend as
    figure 1 in main text.}
\end{figure}

\clearpage
\subsection{Maximum temperature}
\begin{figure}[h!]
  \begin{center}
    \includegraphics[width=\textwidth]{../figures/violin-tmx-0-05.pdf}
  \end{center}
  \caption{\textbf{Maximum temperature, \nth{5} quantile.} Legend as
    figure 1 in main text.}
\end{figure}
\begin{figure}[h!]
  \begin{center}
    \includegraphics[width=\textwidth]{../figures/violin-tmx-0-25.pdf}
  \end{center}
  \caption{\textbf{Maximum temperature, \nth{25} quantile.} Legend as
    figure 1 in main text.}
\end{figure}
\begin{figure}[h!]
  \begin{center}
    \includegraphics[width=\textwidth]{../figures/violin-tmx-0-5.pdf}
  \end{center}
  \caption{\textbf{Maximum temperature, \nth{50} quantile (median).}
    Legend as figure 1 in main text.}
\end{figure}
\begin{figure}[h!]
  \begin{center}
    \includegraphics[width=\textwidth]{../figures/violin-tmx-0-75.pdf}
  \end{center}
  \caption{\textbf{Maximum temperature, \nth{75} quantile.} Legend as
    figure 1 in main text.}
\end{figure}
\begin{figure}[h!]
  \begin{center}
    \includegraphics[width=\textwidth]{../figures/violin-tmx-0-95.pdf}
  \end{center}
  \caption{\textbf{Maximum temperature, \nth{95} quantile.} Legend as
    figure 1 in main text.}
\end{figure}

\clearpage
\subsection{Vapour pressure}
\begin{figure}[h!]
  \begin{center}
    \includegraphics[width=\textwidth]{../figures/violin-vap-0-05.pdf}
  \end{center}
  \caption{\textbf{Vapour pressure, \nth{5} quantile.} Legend as
    figure 1 in main text.}
\end{figure}
\begin{figure}[h!]
  \begin{center}
    \includegraphics[width=\textwidth]{../figures/violin-vap-0-25.pdf}
  \end{center}
  \caption{\textbf{Vapour pressure, \nth{25} quantile.} Legend as
    figure 1 in main text.}
\end{figure}
\begin{figure}[h!]
  \begin{center}
    \includegraphics[width=\textwidth]{../figures/violin-vap-0-5.pdf}
  \end{center}
  \caption{\textbf{Vapour pressure, \nth{50} quantile (median).}
    Legend as figure 1 in main text.}
\end{figure}
\begin{figure}[h!]
  \begin{center}
    \includegraphics[width=\textwidth]{../figures/violin-vap-0-75.pdf}
  \end{center}
  \caption{\textbf{Vapour pressure, \nth{75} quantile.} Legend as
    figure 1 in main text.}
\end{figure}
\begin{figure}[h!]
  \begin{center}
    \includegraphics[width=\textwidth]{../figures/violin-vap-0-95.pdf}
  \end{center}
  \caption{\textbf{Vapour pressure, \nth{95} quantile.} Legend as
    figure 1 in main text.}
\end{figure}

\clearpage
\subsection{Rainday counts}
\begin{figure}[h!]
  \begin{center}
    \includegraphics[width=\textwidth]{../figures/violin-wet-0-05.pdf}
  \end{center}
  \caption{\textbf{Rainday counts, \nth{5} quantile.} Legend as
    figure 1 in main text.}
\end{figure}
\begin{figure}[h!]
  \begin{center}
    \includegraphics[width=\textwidth]{../figures/violin-wet-0-25.pdf}
  \end{center}
  \caption{\textbf{Rainday counts, \nth{25} quantile.} Legend as
    figure 1 in main text.}
\end{figure}
\begin{figure}[h!]
  \begin{center}
    \includegraphics[width=\textwidth]{../figures/violin-wet-0-5.pdf}
  \end{center}
  \caption{\textbf{Rainday counts, \nth{50} quantile (median).}
    Legend as figure 1 in main text.}
\end{figure}
\begin{figure}[h!]
  \begin{center}
    \includegraphics[width=\textwidth]{../figures/violin-wet-0-75.pdf}
  \end{center}
  \caption{\textbf{Rainday counts, \nth{75} quantile.} Legend as
    figure 1 in main text.}
\end{figure}
\begin{figure}[h!]
  \begin{center}
    \includegraphics[width=\textwidth]{../figures/violin-wet-0-95.pdf}
  \end{center}
  \caption{\textbf{Rainday counts, \nth{95} quantile.} Legend as
    figure 1 in main text.}
\end{figure}

\clearpage
\section{Trait correlations and phylogenetic signal (\emph{c.f.} figure 3)}

Please note that, since figure 2 in the main text is an exemplar of
the data underlying the phylogenetic signals shown in these figures,
we do not replicate it here across all data-types as it is
superfluous.

\begin{figure}[h!]
  \begin{center}
    \begin{subfigure}{.45\textwidth}
      \includegraphics[width=\textwidth]{../figures/traits-05.pdf}
      \caption{Trait correlations}
    \end{subfigure}
    \begin{subfigure}{.45\textwidth}
      \includegraphics[width=\textwidth]{../figures/physig-05.pdf}
      \caption{Phylogenetic signal}
    \end{subfigure}
  \end{center}
  \caption{\textbf{Trait correlations and phylogenetic signals of the
      \nth{5} quantiles.} Legend as figure 3 in the main text.}
  \label{traits}
\end{figure}
\begin{figure}[h!]
  \begin{center}
    \begin{subfigure}{.45\textwidth}
      \includegraphics[width=\textwidth]{../figures/traits-25.pdf}
      \caption{Trait correlations}
    \end{subfigure}
    \begin{subfigure}{.45\textwidth}
      \includegraphics[width=\textwidth]{../figures/physig-25.pdf}
      \caption{Phylogenetic signal}
    \end{subfigure}
  \end{center}
  \caption{\textbf{Trait correlations and phylogenetic signals of the
      \nth{25} quantiles.} Legend as figure 3 in the main text.}
  \label{traits}
\end{figure}
\begin{figure}[h!]
  \begin{center}
    \begin{subfigure}{.45\textwidth}
      \includegraphics[width=\textwidth]{../figures/traits-50.pdf}
      \caption{Trait correlations}
    \end{subfigure}
    \begin{subfigure}{.45\textwidth}
      \includegraphics[width=\textwidth]{../figures/physig-5.pdf}
      \caption{Phylogenetic signal}
    \end{subfigure}
  \end{center}
  \caption{\textbf{Trait correlations and phylogenetic signals of the
      \nth{50} quantiles (medians).} Legend as figure 3 in the main
    text.}
  \label{traits}
\end{figure}
\begin{figure}[h!]
  \begin{center}
    \begin{subfigure}{.45\textwidth}
      \includegraphics[width=\textwidth]{../figures/traits-75.pdf}
      \caption{Trait correlations}
    \end{subfigure}
    \begin{subfigure}{.45\textwidth}
      \includegraphics[width=\textwidth]{../figures/physig-75.pdf}
      \caption{Phylogenetic signal}
    \end{subfigure}
  \end{center}
  \caption{\textbf{Trait correlations and phylogenetic signals of the
      \nth{75} quantiles.} Legend as figure 3 in the main text.}
  \label{traits}
\end{figure}
\begin{figure}[h!]
  \begin{center}
    \begin{subfigure}{.45\textwidth}
      \includegraphics[width=\textwidth]{../figures/traits-95.pdf}
      \caption{Trait correlations}
    \end{subfigure}
    \begin{subfigure}{.45\textwidth}
      \includegraphics[width=\textwidth]{../figures/physig-95.pdf}
      \caption{Phylogenetic signal}
    \end{subfigure}
  \end{center}
  \caption{\textbf{Trait correlations and phylogenetic signals of the
      \nth{95} quantiles.} Legend as figure 3 in the main text.}
  \label{traits}
\end{figure}

\clearpage
\section{Principal components analyses (\emph{c.f.} figure 4)}

\begin{figure}[h!]
  \begin{center}
    \begin{subfigure}{.45\textwidth}
      \includegraphics[width=\textwidth]{../figures/pca-005-past.pdf}
      \caption{Past climate space (1955--1980)}
    \end{subfigure}
    \begin{subfigure}{.45\textwidth}
      \includegraphics[width=\textwidth]{../figures/pca-005-present.pdf}
      \caption{Present climate space (1990--2015)}
    \end{subfigure}
    \begin{subfigure}{.45\textwidth}
      \includegraphics[width=\textwidth]{../figures/pca-005-index.pdf}
      \caption{Tracking climate space}
    \end{subfigure}
  \end{center}
  \caption{\doublespacing \textbf{Principal components analyses for
      the \nth{5} quantiles.} Legend as figure 4 in the main text.}
  \label{pca}
\end{figure}
\begin{figure}[h!]
  \begin{center}
    \begin{subfigure}{.45\textwidth}
      \includegraphics[width=\textwidth]{../figures/pca-025-past.pdf}
      \caption{Past climate space (1955--1980)}
    \end{subfigure}
    \begin{subfigure}{.45\textwidth}
      \includegraphics[width=\textwidth]{../figures/pca-025-present.pdf}
      \caption{Present climate space (1990--2015)}
    \end{subfigure}
    \begin{subfigure}{.45\textwidth}
      \includegraphics[width=\textwidth]{../figures/pca-025-index.pdf}
      \caption{Tracking climate space}
    \end{subfigure}
  \end{center}
  \caption{\doublespacing \textbf{Principal components analyses for
      the \nth{25} quantiles.} Legend as figure 4 in the main text.}
  \label{pca}
\end{figure}
\begin{figure}[h!]
  \begin{center}
    \begin{subfigure}{.45\textwidth}
      \includegraphics[width=\textwidth]{../figures/pca-05-past.pdf}
      \caption{Past climate space (1955--1980)}
    \end{subfigure}
    \begin{subfigure}{.45\textwidth}
      \includegraphics[width=\textwidth]{../figures/pca-05-present.pdf}
      \caption{Present climate space (1990--2015)}
    \end{subfigure}
    \begin{subfigure}{.45\textwidth}
      \includegraphics[width=\textwidth]{../figures/pca-05-index.pdf}
      \caption{Tracking climate space}
    \end{subfigure}
  \end{center}
  \caption{\doublespacing \textbf{Principal components analyses for
      the \nth{50} quantiles (\emph{i.e.}, medians).} Legend as figure
    4 in the main text.}
  \label{pca}
\end{figure}
\begin{figure}[h!]
  \begin{center}
    \begin{subfigure}{.45\textwidth}
      \includegraphics[width=\textwidth]{../figures/pca-075-past.pdf}
      \caption{Past climate space (1955--1980)}
    \end{subfigure}
    \begin{subfigure}{.45\textwidth}
      \includegraphics[width=\textwidth]{../figures/pca-075-present.pdf}
      \caption{Present climate space (1990--2015)}
    \end{subfigure}
    \begin{subfigure}{.45\textwidth}
      \includegraphics[width=\textwidth]{../figures/pca-075-index.pdf}
      \caption{Tracking climate space}
    \end{subfigure}
  \end{center}
  \caption{\doublespacing \textbf{Principal components analyses for
      the \nth{75} quantiles.} Legend as figure 4 in the main text.}
  \label{pca}
\end{figure}
\begin{figure}[h!]
  \begin{center}
    \begin{subfigure}{.45\textwidth}
      \includegraphics[width=\textwidth]{../figures/pca-095-past.pdf}
      \caption{Past climate space (1955--1980)}
    \end{subfigure}
    \begin{subfigure}{.45\textwidth}
      \includegraphics[width=\textwidth]{../figures/pca-095-present.pdf}
      \caption{Present climate space (1990--2015)}
    \end{subfigure}
    \begin{subfigure}{.45\textwidth}
      \includegraphics[width=\textwidth]{../figures/pca-095-index.pdf}
      \caption{Tracking climate space}
    \end{subfigure}
  \end{center}
  \caption{\doublespacing \textbf{Principal components analyses for
      the \nth{95} quantiles.} Legend as figure 4 in the main text.}
  \label{pca}
\end{figure}

\end{document}

%%% Local Variables:
%%% mode: latex
%%% TeX-master: t
%%% End:
