## supplemental code for "The realised velocity of climate change reveals remarkable idiosyncrasy of species’ distributional shifts": README.md

The realised velocity of climate change reveals remarkable idiosyncrasy of species distributional shifts
============================================
William D. Pearse and T. Jonathan Davies

## Overview 
This is the code underlying the manuscript "The realised velocity of climate change reveals remarkable idiosyncrasy of species distributional shifts". We emphasise that this code and manuscript has not yet been peer-reviewed.

## How to use this code

Optional steps before running the code:

1. (Optional.) If using different different GBIF data dumps, modify the URLs in `src/download-gbif.rb` to match your new data, and the filenames in `src/calculate_index.R` to match those new files. Note that GBIF sometimes change file header order and contents - check the output from `src/download-gif.rb` to make sure you get what you expect.
2. (Optional.) If you wish to repeat our supplementary analyses of minimum record counts, change the `min.records` arguments in line 53 of `src/calculate_index.R` onwards to be `10000`.
3. (Optional.) If you wish to repeat our supplementary analyses of minimum record counts, specify `clean.gbif=TRUE` in line 53 of `src/calculate_index.R` onwards in the `.load.gbif` function calls. Change line 7 of `src/download-gbif.rb` so that it reads `clean_gbif=true`.

Mandatory steps to re-run our analysis:

1. Open `src/headers.R` and modify the output directories to suit your needs. These changes will propagate through to all the R code.
2. Open `src/download-cru.rb` and modify line 5 to match the directory specified in step 1.
3. Open `src/download-gbif.rb` and modify line 6 to match the directory specified in step 1.
4. Install all necessary R package by running something like `Rscript src/headers.R`. Check for errors and correct as necessary!
5. Open the file `raw-data/metadata.txt` and got to the URLs inside to dwonload the trait data and save it under the filenames listed inside the meta-data file. The licensing restrictions of (some) of these datasets mean I cannot redistribute them in this repostiory.
6. Ensure `ruby`, `get`, `unzip`, `cut`, and (if you want the manuscript itself)`pdflatex` and `biber` are installed on your machine.
7. You probably need to be on a Linux machine; MacOS will probably work if (6) is caried out; Windows likely won't.
8. Run everything by typing `ruby Rakefile`
9. Wait a very long time - potentially several days.
