## supplemental code for "The realised velocity of climate change reveals remarkable idiosyncrasy of species’ distributional shifts": analysis-pca.R

```
# Headers
source("src/headers.R")

# Wrapper functions
.combined.plot <- function(quantile, dims=1:2, clean=5, abs=FALSE){

    index <- na.omit(.simplify.group("bootstrap.index", quantile, clean=clean,  abs=abs)[,1:9])
    present <- na.omit(.simplify.group("present", quantile, abs=abs)[,1:9])
    past <- na.omit(.simplify.group("past", quantile, abs=abs)[,1:9])
    
    #layout(matrix(1:6, nrow=2, byrow=TRUE), widths=c(5,5,5), heights=c(5,1))
    par(mar=c(3.1,3.1,1.1,1.1), mfrow=c(1,3))
    present <- prcomp(present, scale=TRUE)
    past <- prcomp(past, scale=TRUE)
    index <- prcomp(index, scale=TRUE)
    .pca.plot(past, 1:2, bty="o")
    .pca.plot(present, 1:2)
    .pca.plot(index, 1:2)
}
.pca.plot <- function(pca, dims=1:2, scale=5, ...){
    density <- kde(pca$x[,dims])
    var <- pca$sdev^2
    p.var <- (var / sum(var)) * 100
    par(mar=c(4.5, 4.5, 1.1, 1.1))
    plot(density, display="filled.contour", cont=seq(10,90,10), xlab=paste0("PC1 (% variance = ",round(p.var[1]),")"), ylab=paste0("PC2 (% variance = ",round(p.var[2]),")"), cex.lab=1.5, cex.axis=1.25)
    points(pca$x, pch=20, cex=.5, col=alpha("grey60",.25))
    arrows(rep(0,ncol(pca$x)),rep(0,ncol(pca$x)), pca$rotation[,dims[1]]*scale, pca$rotation[,dims[2]]*scale, length=.1, lwd=3)
    text.locs <- pca$rotation*scale*1.3
    text(text.locs[,dims[1]], text.locs[,dims[2]], names(pca$center), font=2)
}

# Load data
plants <- readRDS("clean-data/plants-index.RDS")
fungi <- readRDS("clean-data/fungi-index.RDS")
insects <- readRDS("clean-data/insects-index.RDS")
mammals <- readRDS("clean-data/mammals-index.RDS")
reptiles <- readRDS("clean-data/reptiles-index.RDS")
birds <- readRDS("clean-data/birds-index.RDS")
amphibians <- readRDS("clean-data/amphibians-index.RDS")

# PCAs
for(quant in c("0.05","0.25","0.5","0.75","0.95")){
    past <- prcomp(na.omit(.simplify.group("past", quant)[,1:9]), scale=TRUE)
    pdf(paste0("figures/pca-",gsub("0.","0",quant,fixed=TRUE),"-past.pdf")); .pca.plot(past); dev.off()
    present <- prcomp(na.omit(.simplify.group("present", quant)[,1:9]), scale=TRUE)
    pdf(paste0("figures/pca-",gsub("0.","0",quant,fixed=TRUE),"-present.pdf")); .pca.plot(present); dev.off()
    index <- prcomp(na.omit(.simplify.group("bootstrap.index", quant, clean=10)[,1:9]), scale=TRUE)
    pdf(paste0("figures/pca-",gsub("0.","0",quant,fixed=TRUE),"-index.pdf")); .pca.plot(index); dev.off()
}


pdf("figures/pca-05.pdf", width=15, height=6); .combined.plot("0.05"); dev.off()
pdf("figures/pca-25.pdf", width=15, height=6); .combined.plot("0.25"); dev.off()
pdf("figures/pca-50.pdf", width=15, height=6); .combined.plot("0.5"); dev.off()
pdf("figures/pca-75.pdf", width=15, height=6); .combined.plot("0.75"); dev.off()
pdf("figures/pca-95.pdf", width=15, height=6); .combined.plot("0.95"); dev.off()

# Simulating PCAs
correls <- numeric(100)
real <- prcomp(combined.pres[,1:9], scale=TRUE)
for(i in seq_along(correls)){
    past <- combined.past[,1:9]
    for(i in seq_len(9))
        past[,i] <- past[,i] + rnorm(nrow(past), sd=sd(abs(combined.pres[,i] - combined.past[,i])))
    pres <- past
    for(i in seq_len(9))
        past[,i] <- past[,i] + rnorm(nrow(past), sd=sd(abs(combined.pres[,i] - combined.past[,i])))
    diff <- pres - past
    past.pca <- prcomp(past, scale=TRUE)
    diff.pca <- prcomp(diff, scale=TRUE)
    correls[i] <- mantel.test(abs(diff.pca$rotation), abs(past.pca$rotation))$z.stat
}
```
