## supplemental code for "The realised velocity of climate change reveals remarkable idiosyncrasy of species’ distributional shifts": analysis-phylo.R

```
# Headers
source("src/headers.R")

# Wrapper functions
.make.comparative.data <- function(data, tree, merge=TRUE){
    tree$tip.label <- tolower(gsub(" ", "_", tree$tip.label))
    tree$node.label <- NULL
    data <- as.data.frame(data)
    data$species <- rownames(data)
    data$taxon <- NULL
    data <- data[!duplicated(data$species),]
    if(merge)
        tree <- congeneric.merge(tree, data$species)
    return(comparative.data(tree, data, species))
}
.phy.sig <- function(c.data, method="lambda")
    sapply(c.data$data, function(x) phylosig(c.data$phy, x, method))
.dom.niche <- function(c.data)
    setNames(
        apply(as.matrix(c.data$data), 1, function(x) which.min(abs(x))),
        rownames(c.data$data)
    )
.phy.sig.wrap <- function(data, tree, index, clean, method="lambda", merge=TRUE, p.val=FALSE){
    output <- matrix(NA, ncol=9, nrow=5)
    rownames(output) <- dimnames(data)[[2]]
    colnames(output) <- dimnames(data)[[1]]
    
    for(i in seq_len(nrow(output))){
        c.data <- .make.comparative.data(.simplify(data, index, rownames(output)[i]), tree, merge)
        if(p.val){
            output[i,] <- as.numeric(.phy.sig(c.data)[2,])
        } else {
            output[i,] <- as.numeric(.phy.sig(c.data)[1,])
        }
            
    }
    return(output)
}
.dom.niche.wrap <- function(data, tree, method="lambda", merge=TRUE, clean=5){
    output <- setNames(numeric(nrow(data)), dimnames(data)[[1]])
    for(i in seq_len(nrow(output))){
        c.data <- .make.comparative.data(.simplify(data, "bootstrap.index", rownames(output)[i], clean=clean), tree, merge)
        model <- fitDiscrete(multi2di(c.data$phy), model="SYM", .dom.niche(c.data), transform="lambda")
        output[i] <- model$opt$lambda
    }
    return(output)
}
.get.branch.trait <- function(trait, tree){
    trait <- c(trait[tree$tip.label], fastAnc(tree, trait))
    names(trait)[1:length(tree$tip.label)] <- 1:length(tree$tip.label)
    edge.mat <- matrix(trait[tree$edge], nrow(tree$edge), 2)
    trait <- rowMeans(edge.mat)
    return(trait)
}
.limit <- function(x){
    min <- min(x); max <- max(x)
    x <- x - min
    x <- x/max(x)
    attr(x, "range") <- c(min, max)
    return(x)
}
.plot.phylo.mat <- function(quant){
    q.var <- setNames(
        c("5th","25th","50th","75th","95th"),
        c("0.05","0.25","0.5","0.75","0.95")
    )[quant]
    c.var <- setNames(
        c("clouds","frost","evapo-trans.","precipitation","min(temp)","mean(temp)","max(temp)","vapour","rainy day"),
        c("cld","frs","pet","pre","tmn","tmp","tmx","vap","wet")
    )
    
    index <- matrix(
        c(readRDS("clean-data/mammals-signal.RDS")$bootstrap.index[quant,],
          readRDS("clean-data/birds-signal.RDS")$bootstrap.index[quant,],
          readRDS("clean-data/reptiles-signal.RDS")$bootstrap.index[quant,],
          readRDS("clean-data/plants-signal.RDS")$bootstrap.index[quant,])
      , byrow=FALSE, nrow=length(c.var)
    )
    comp <- matrix(
        c(readRDS("clean-data/mammals-signal.RDS")$present[quant,],
          readRDS("clean-data/birds-signal.RDS")$present[quant,],
          readRDS("clean-data/reptiles-signal.RDS")$present[quant,],
          readRDS("clean-data/plants-signal.RDS")$present[quant,])
      , byrow=FALSE, nrow=length(c.var)
    )
    colnames(comp) <- colnames(index) <- c("mammals","birds","reptiles","plants")
    rownames(comp) <- rownames(index) <- c.var
    
    cols <- colorRampPalette(c("blue", "white", "red"))
    bugged.cols <- colorRampPalette(c("green","green","blue", "white", "red"))
    max.lam <- max(c(index,comp))
    comp.cuts <- as.numeric(cut(as.numeric(comp), breaks=seq(0,max.lam,length.out=201)))
    comp.cuts <- cols(200)[comp.cuts]
    index.cuts <- as.numeric(cut(as.numeric(index), breaks=seq(0,max.lam,length.out=201)))
    index.cuts <- cols(200)[index.cuts]
    index.sig <- matrix(as.numeric(index > comp), nrow=nrow(index))
    
    corrplot(index, method="square", is.corr=FALSE, cl.lim=c(0,max.lam), tl.srt=45, col=bugged.cols(400), tl.col="black")
    corrplot(comp, method="square", is.corr=FALSE, cl.lim=c(0,max.lam), tl.srt=45, bg=comp.cuts, add=TRUE, addgrid.col=NA, col=alpha("white",0), cl.pos="n", tl.pos="n")
    corrplot(index, method="square", is.corr=FALSE, cl.lim=c(0,max.lam), tl.srt=45, bg=alpha("white",0), col=index.cuts, add=TRUE, addgrid.col=NA, cl.pos="n", tl.pos="n", p.mat=index.sig)
    text(-1.3, 10.5, "Legend", font=2)
    rect(-1, 9.5, 0, 10.5, col=cols(200)[150], border=NA)
    rect(-.8, 9.7, -.2, 10.3, col=cols(200)[120], border=NA)
    text(-.5, 10.4, "present")
    text(-.5, 10, "track\nindex")
}

# Load data
plants <- readRDS("clean-data/plants-index.RDS")
fungi <- readRDS("clean-data/fungi-index.RDS")
insects <- readRDS("clean-data/insects-index.RDS")
mammals <- readRDS("clean-data/mammals-index.RDS")
reptiles <- readRDS("clean-data/reptiles-index.RDS")
birds <- readRDS("clean-data/birds-index.RDS")

# Calculate signals
if(FALSE){
    saveRDS(list(
    bootstrap.index=.phy.sig.wrap(mammals, read.tree("phylogenies/faurby_mammals.tre"), "bootstrap.index"),
    index=.phy.sig.wrap(mammals, read.tree("phylogenies/faurby_mammals.tre"), "index"),
    present=.phy.sig.wrap(mammals, read.tree("phylogenies/faurby_mammals.tre"), "present"),
    past=.phy.sig.wrap(mammals, read.tree("phylogenies/faurby_mammals.tre"), "past"),
    projected=.phy.sig.wrap(mammals, read.tree("phylogenies/faurby_mammals.tre"), "projected")
), "clean-data/mammals-signal.RDS")
saveRDS(list(
    bootstrap.index=.phy.sig.wrap(birds, read.tree("phylogenies/bird_tree.tre"), "bootstrap.index"),
    index=.phy.sig.wrap(birds, read.tree("phylogenies/bird_tree.tre"), "index"),
    present=.phy.sig.wrap(birds, read.tree("phylogenies/bird_tree.tre"), "present"),
    past=.phy.sig.wrap(birds, read.tree("phylogenies/bird_tree.tre"), "past"),
    projected=.phy.sig.wrap(birds, read.tree("phylogenies/bird_tree.tre"), "projected")
), "clean-data/birds-signal.RDS")
saveRDS(list(
    bootstrap.index=.phy.sig.wrap(reptiles, read.tree("phylogenies/squamate_tree.tre"), "bootstrap.index"),
    index=.phy.sig.wrap(reptiles, read.tree("phylogenies/squamate_tree.tre"), "index"),
    present=.phy.sig.wrap(reptiles, read.tree("phylogenies/squamate_tree.tre"), "present"),
    past=.phy.sig.wrap(reptiles, read.tree("phylogenies/squamate_tree.tre"), "past"),
    projected=.phy.sig.wrap(reptiles, read.tree("phylogenies/squamate_tree.tre"), "projected")
), "clean-data/reptiles-signal.RDS")
if(FALSE)
    saveRDS(list(
        bootstrap.index=.phy.sig.wrap(amphibians, read.tree("phylogenies/amphibian_tree.tre"), "bootstrap.index"),
        index=.phy.sig.wrap(amphibians, read.tree("phylogenies/amphibian_tree.tre"), "index"),
        present=.phy.sig.wrap(amphibians, read.tree("phylogenies/amphibian_tree.tre"), "present"),
        past=.phy.sig.wrap(amphibians, read.tree("phylogenies/amphibian_tree.tre"), "past"),
        projected=.phy.sig.wrap(amphibians, read.tree("phylogenies/amphibian_tree.tre"), "projected")
    ), "clean-data/amphibians-signal.RDS")
saveRDS(list(
    bootstrap.index=.phy.sig.wrap(plants, read.tree("phylogenies/ALLOTB.tre"), "bootstrap.index"),
    index=.phy.sig.wrap(plants, read.tree("phylogenies/ALLOTB.tre"), "index"),
    present=.phy.sig.wrap(plants, read.tree("phylogenies/ALLOTB.tre"), "present"),
    past=.phy.sig.wrap(plants, read.tree("phylogenies/ALLOTB.tre"), "past"),
    projected=.phy.sig.wrap(plants, read.tree("phylogenies/ALLOTB.tre"), "projected")
), "clean-data/plants-signal.RDS")
}

# Get p-values
saveRDS(list(
    bootstrap.index=.phy.sig.wrap(mammals, read.tree("phylogenies/faurby_mammals.tre"), "bootstrap.index"),
    index=.phy.sig.wrap(mammals, read.tree("phylogenies/faurby_mammals.tre"), "index"),
    present=.phy.sig.wrap(mammals, read.tree("phylogenies/faurby_mammals.tre"), "present"),
    past=.phy.sig.wrap(mammals, read.tree("phylogenies/faurby_mammals.tre"), "past"),
    projected=.phy.sig.wrap(mammals, read.tree("phylogenies/faurby_mammals.tre"), "projected")
), "clean-data/mammals-signal-p.RDS")
saveRDS(list(
    bootstrap.index=.phy.sig.wrap(birds, read.tree("phylogenies/bird_tree.tre"), "bootstrap.index"),
    index=.phy.sig.wrap(birds, read.tree("phylogenies/bird_tree.tre"), "index"),
    present=.phy.sig.wrap(birds, read.tree("phylogenies/bird_tree.tre"), "present"),
    past=.phy.sig.wrap(birds, read.tree("phylogenies/bird_tree.tre"), "past"),
    projected=.phy.sig.wrap(birds, read.tree("phylogenies/bird_tree.tre"), "projected")
), "clean-data/birds-signal-p.RDS")
saveRDS(list(
    bootstrap.index=.phy.sig.wrap(reptiles, read.tree("phylogenies/squamate_tree.tre"), "bootstrap.index"),
    index=.phy.sig.wrap(reptiles, read.tree("phylogenies/squamate_tree.tre"), "index"),
    present=.phy.sig.wrap(reptiles, read.tree("phylogenies/squamate_tree.tre"), "present"),
    past=.phy.sig.wrap(reptiles, read.tree("phylogenies/squamate_tree.tre"), "past"),
    projected=.phy.sig.wrap(reptiles, read.tree("phylogenies/squamate_tree.tre"), "projected")
), "clean-data/reptiles-signal-p.RDS")
if(FALSE)
    saveRDS(list(
        bootstrap.index=.phy.sig.wrap(amphibians, read.tree("phylogenies/amphibian_tree.tre"), "bootstrap.index"),
        index=.phy.sig.wrap(amphibians, read.tree("phylogenies/amphibian_tree.tre"), "index"),
        present=.phy.sig.wrap(amphibians, read.tree("phylogenies/amphibian_tree.tre"), "present"),
        past=.phy.sig.wrap(amphibians, read.tree("phylogenies/amphibian_tree.tre"), "past"),
        projected=.phy.sig.wrap(amphibians, read.tree("phylogenies/amphibian_tree.tre"), "projected")
    ), "clean-data/amphibians-signal-p.RDS")
saveRDS(list(
    bootstrap.index=.phy.sig.wrap(plants, read.tree("phylogenies/ALLOTB.tre"), "bootstrap.index"),
    index=.phy.sig.wrap(plants, read.tree("phylogenies/ALLOTB.tre"), "index"),
    present=.phy.sig.wrap(plants, read.tree("phylogenies/ALLOTB.tre"), "present"),
    past=.phy.sig.wrap(plants, read.tree("phylogenies/ALLOTB.tre"), "past"),
    projected=.phy.sig.wrap(plants, read.tree("phylogenies/ALLOTB.tre"), "projected")
), "clean-data/plants-signal-p.RDS")

# Estimate signal fractions
comp <- matrix(
    c(readRDS("clean-data/mammals-signal.RDS")$present,
      readRDS("clean-data/birds-signal.RDS")$present,
      readRDS("clean-data/reptiles-signal.RDS")$present,
      readRDS("clean-data/plants-signal.RDS")$present)
  , byrow=TRUE, ncol=9
)
sum(comp > .2)
comp <- matrix(
    c(readRDS("clean-data/mammals-signal.RDS")$past,
      readRDS("clean-data/birds-signal.RDS")$past,
      readRDS("clean-data/reptiles-signal.RDS")$past,
      readRDS("clean-data/plants-signal.RDS")$past)
  , byrow=TRUE, ncol=9
)
sum(comp > .2)
index <- matrix(
    c(readRDS("clean-data/mammals-signal.RDS")$bootstrap.index,
      readRDS("clean-data/birds-signal.RDS")$bootstrap.index,
      readRDS("clean-data/reptiles-signal.RDS")$bootstrap.index,
      readRDS("clean-data/plants-signal.RDS")$bootstrap.index)
  , byrow=TRUE, ncol=9
)
sum(index > .2)

# Make phylogenetic signal figures
pdf("figures/physig-05.pdf");.plot.phylo.mat("0.05");dev.off()
pdf("figures/physig-25.pdf");.plot.phylo.mat("0.25");dev.off()
pdf("figures/physig-5.pdf");.plot.phylo.mat("0.5");dev.off()
pdf("figures/physig-75.pdf");.plot.phylo.mat("0.75");dev.off()
pdf("figures/physig-95.pdf");.plot.phylo.mat("0.95");dev.off()


# Plot trait(s) on phylogeny
c.index <- .make.comparative.data(.simplify(plants, "bootstrap.index", "0.5", 10), ladderize(read.tree("phylogenies/ALLOTB.tre")), FALSE)
c.past <- .make.comparative.data(.simplify(plants, "present", "0.5"), c.index$phy, FALSE)

branches <- .get.branch.trait(setNames(c.past$data$tmp,rownames(c.past$data)), c.past$phy)
cut.branches <- as.numeric(cut(branches, seq(floor(min(branches)), ceiling(max(branches)), length.out=10000)))
cut.tips <- as.numeric(cut(c.index$data$tmp, seq(floor(min(c.index$data$tmp)),ceiling(max(c.index$data$tmp)), length.out=10000)))
cols <- viridis(10001)

pdf("figures/phylo-tmp-50.pdf", width=15, height=4)
edge.types <- rep(1, nrow(c.past$phy$edge))
edge.types[c(1,3353)] <- 2
tree <- c.past$phy
tree$edge.length[c(1,3353)] <- tree$edge.length[c(1,3353)] - 100
bg <- plot(tree, direction="up", no.margin=TRUE, show.tip.label=FALSE, edge.color=cols[cut.branches], x.lim=c(-70, 1850), y.lim=c(0,240), edge.lty=edge.types)
tiplabels(pch="|", col=cols[cut.tips], adj=c(0,10))
gradient.rect(bg$x.lim[1]+5, 30, bg$x.lim[2]*.6, 45, col=cols, border=NA)
xs <- seq(bg$x.lim[1]+5,bg$x.lim[2]*.6,length.out=10)
text(xs, 30, labels=seq(floor(min(branches)),ceiling(max(branches)), length.out=10))
text(xs, 45, labels=round(seq(floor(min(c.index$data$tmp)),ceiling(max(c.index$data$tmp)), length.out=10)))
text(xs[10]*1.1, c(30,45), labels=c(expression(temp[50]), expression(track[50](temp))))
text(bg$x.lim[1], bg$y.lim[2]-5, expression(track[50](temp)))
text(bg$x.lim[1], bg$y.lim[2]-20, expression(temp[50]))
text(bg$x.lim[2], bg$y.lim[2]-5, expression(track[50](temp)))
text(bg$x.lim[2], bg$y.lim[2]-20, expression(temp[50]))
abline(h=227, col="grey40", lty=2)
dev.off()
```
