## supplemental code for "The realised velocity of climate change reveals remarkable idiosyncrasy of species’ distributional shifts": analysis-traits.R

```
# Headers
source("src/headers.R")

# Wrapper functions
.univariate <- function(data, explanatory){
    tests <- lapply(data, function(x) cor.test(x, explanatory))
    cors <- sapply(tests, function(x) x$estimate)
    p <- sapply(tests, function(x) x$p.value)
    p <- ifelse(p < .2, round(p,3), NA)
    output <- matrix(c(cors,p), nrow=2, byrow=TRUE)
    colnames(output) <- names(data); rownames(output) <- c("r","p")
    return(output)
}
.combine.data <- function(index, quant, clean=10, abs=FALSE){
    data <- .simplify.group(index, quant, clean, abs)
    data$species <- rownames(data)
    
    data$bm.mamm <- neotoma$log.mass[match(data$species, neotoma$binomial)]
    data$bm.mamm[data$taxon != "mammals"] <- NA
    data$bm.bird <- elton.bm[data$species]
    data$bm.bird[data$taxon != "birds"] <- NA

    data$log.ll <- glopnet$log.LL[match(data$species, glopnet$Species)]
    data$log.lma <- glopnet$log.LMA[match(data$species, glopnet$Species)]
    data$log.Nmass <- glopnet$log.Nmass[match(data$species, glopnet$Species)]
    data$log.Amass <- glopnet$log.Amass[match(data$species, glopnet$Species)]
    return(data)    
}
.corr.mat <- function(index, quant, clean, abs=FALSE, p.val=FALSE){
    data <- .combine.data(index, quant, clean, abs)

    c.var <- setNames(
        c("clouds","frost","evapo-trans.","precipitation","min(temp)","mean(temp)","max(temp)","vapour","rainy day"),
        c("cld","frs","pet","pre","tmn","tmp","tmx","vap","wet")
    )
    q.var <- setNames(
        c("5th","25th","50th","75th","95th"),
        c("0.05","0.25","0.5","0.75","0.95")
    )[quant]
    t.var <- setNames(
        c("mammal mass","bird mass","leaf lifespan","leaf mass/area","leaf N","photosynth"),
        c("bm.mamm","bm.bird","log.ll","log.lma","log.Nmass","log.Amass")
    )
    
    mat <- matrix(nrow=length(t.var), ncol=length(c.var), dimnames=list(t.var, c.var))
    for(i in seq_along(c.var)){
        for(j in seq_along(t.var)){
            if(p.val){
                mat[j,i] <- cor.test(data[,names(c.var)[i]], data[,names(t.var)[j]])$p.value               
            } else {
                mat[j,i] <- cor.test(data[,names(c.var)[i]], data[,names(t.var)[j]])$estimate
            }
        }
    }
    return(mat)
}
.plot.corr.mat <- function(index, comparison, quant, clean, abs=FALSE){
    index.mat <- t(.corr.mat(index, quant, clean, abs))
    index.mat.p <- t(.corr.mat(index, quant, clean, abs, p.val=TRUE))
    signif <- index.mat.p < .05
    index.mat.p <- matrix(0, nrow=nrow(index.mat), ncol=ncol(index.mat))
    index.mat.p[signif] <- 1
    comparison.mat <- t(.corr.mat(comparison, quant, clean=NA))
    cols <- colorRampPalette(c("red", "white", "blue"))
                   
    comparison.cuts <- as.numeric(cut(as.numeric(comparison.mat), breaks=seq(-1,1,length.out=201)))
    comparison.cuts <- cols(200)[comparison.cuts]
    index.cuts <- as.numeric(cut(as.numeric(index.mat), breaks=seq(-1,1,length.out=201)))
    index.cuts <- cols(200)[index.cuts]
    dummy.mat <- matrix(0, nrow=nrow(index.mat), ncol=ncol(index.mat), dimnames=dimnames(index.mat))
    
    corrplot(comparison.mat, method="square", is.corr=FALSE, cl.lim=c(-1,1), tl.srt=45, col=cols(200), tl.col="black")
    corrplot(comparison.mat, method="square", is.corr=FALSE, cl.lim=c(-1,1), tl.srt=45, bg=comparison.cuts, add=TRUE, addgrid.col=NA, col=alpha("white",0), cl.pos="n", tl.pos="n")
    corrplot(index.mat, method="square", is.corr=FALSE, cl.lim=c(-1,1), tl.srt=45, bg=alpha("white",0), col=index.cuts, add=TRUE, addgrid.col=NA, cl.pos="n", , tl.pos="n", p.mat=index.mat.p, sig.level=.05)
    text(-1.3, 9.5, "Legend", font=2)
    rect(-1, 8.5, 0, 9.5, col=cols(200)[150], border=NA)
    rect(-.8, 8.7, -.2, 9.3, col=cols(200)[120], border=NA)
    text(-.5, 9.4, "present")
    text(-.5, 9, "track\nindex")
}

# Load tracking data
plants <- readRDS("clean-data/plants-index.RDS")
fungi <- readRDS("clean-data/fungi-index.RDS")
insects <- readRDS("clean-data/insects-index.RDS")
mammals <- readRDS("clean-data/mammals-index.RDS")
reptiles <- readRDS("clean-data/reptiles-index.RDS")
birds <- readRDS("clean-data/birds-index.RDS")
amphibians <- readRDS("clean-data/amphibians-index.RDS")
combined <- .simplify.group("bootstrap.index", "0.5", clean=10)

# Load Neotoma data
neotoma <- read.csv("raw-data/Amniote_Database_Aug_2015.csv", as.is=TRUE)
for(i in seq_along(names(neotoma)))
    neotoma[neotoma[,i]==-999,i] <- NA
neotoma$binomial <- with(neotoma, tolower(paste(genus, species, sep="_")))
neotoma$log.mass <- log10(neotoma$adult_body_mass_g)

# Load Glopnet
glopnet <- read.xls("raw-data/glopnet.xls")
glopnet$Species <- tolower(gsub(" ", "_", glopnet$Species))

# Load Elton
elton.birds <- read.delim("raw-data/BirdFuncDat.txt", as.is=TRUE)
elton.birds$Scientific <- tolower(gsub(" ", "_", elton.birds$Scientific))
elton.mam <- read.delim("raw-data/MamFuncDat.txt", as.is=TRUE)
elton.mam$Scientific <- tolower(gsub(" ", "_", elton.mam$Scientific))
elton.bm <- setNames(c(elton.mam$BodyMass.Value, elton.birds$BodyMass.Value), c(elton.mam$Scientific,elton.birds$Scientific))
elton.bm <- log10(elton.bm)

# Combine
pdf("figures/traits-05.pdf"); .plot.corr.mat("bootstrap.index", "present", "0.05", clean=5); dev.off()
pdf("figures/traits-25.pdf"); .plot.corr.mat("bootstrap.index", "present", "0.25", 5); dev.off()
pdf("figures/traits-50.pdf"); .plot.corr.mat("bootstrap.index", "present", "0.5", 5); dev.off()
pdf("figures/traits-75.pdf"); .plot.corr.mat("bootstrap.index", "present", "0.75", 5); dev.off()
pdf("figures/traits-95.pdf"); .plot.corr.mat("bootstrap.index", "present", "0.95", 5); dev.off()

# Correlations

index.p <- c(
    .corr.mat("bootstrap.index", "0.05", clean=100, p.val=TRUE),
    .corr.mat("bootstrap.index", "0.25", clean=100, p.val=TRUE),
    .corr.mat("bootstrap.index", "0.5", clean=100, p.val=TRUE),
    .corr.mat("bootstrap.index", "0.75", clean=100, p.val=TRUE),
    .corr.mat("bootstrap.index", "0.95", clean=100, p.val=TRUE)
)
sum(index.p < .05); length(index.p)

pres.p <- c(
    .corr.mat("present", "0.05", clean=NA, p.val=TRUE),
    .corr.mat("present", "0.25", clean=NA, p.val=TRUE),
    .corr.mat("present", "0.5", clean=NA, p.val=TRUE),
    .corr.mat("present", "0.75", clean=NA, p.val=TRUE),
    .corr.mat("present", "0.95", clean=NA, p.val=TRUE)
)
sum(pres.p < .05); length(pres.p)
past.p <- c(
    .corr.mat("past", "0.05", clean=NA, p.val=TRUE),
    .corr.mat("past", "0.25", clean=NA, p.val=TRUE),
    .corr.mat("past", "0.5", clean=NA, p.val=TRUE),
    .corr.mat("past", "0.75", clean=NA, p.val=TRUE),
    .corr.mat("past", "0.95", clean=NA, p.val=TRUE)
)
sum(past.p < .05); length(past.p)
```
