## supplemental code for "The realised velocity of climate change reveals remarkable idiosyncrasy of species’ distributional shifts": analysis-violin.R

```
# Headers
source("src/headers.R")

# Load empirical data
plants <- readRDS("clean-data/plants-index.RDS")
fungi <- readRDS("clean-data/fungi-index.RDS")
insects <- readRDS("clean-data/insects-index.RDS")
mammals <- readRDS("clean-data/mammals-index.RDS")
reptiles <- readRDS("clean-data/reptiles-index.RDS")
birds <- readRDS("clean-data/birds-index.RDS")
amphibians <- readRDS("clean-data/amphibians-index.RDS")

# Load simulated data
load("src/sim-range.RData")
mid <- t(sapply(results, function(x) x[1,]))
data <- cbind(data, mid)
data <- data[!is.na(data$bootstrap.index),]
sim.data <- data[is.finite(data$index),]

# Do the plots
.outline.plot <- function(climate, quant, clean=NA, se.thresh=5, abs=FALSE, index="bootstrap.index", sims=FALSE){
    # Prep simulated data
    #sim.data <- sim.data[as.character(sim.data$quantile)==quant,]
    warning("Currently only have simulated data for 0.5 quantile and temperature")
    sim.data <- sim.data[order(sim.data$alpha),]
    sim.data$index <- sim.data[,index]
    sim.data$alpha.cat <- ifelse(sim.data$alpha==0.5, "intermediate", ifelse(sim.data$alpha==0, "static", "tracking"))
    sim.data$j.a.cat <- factor(sim.data$alpha.cat, levels=c("static","intermediate","tracking"))
    sim.data$j.a.cat <- as.numeric(sim.data$j.a.cat)

    # Load and match data
    index <- .simplify.group(index, quantile=quant, clean=clean, abs=abs)
    data <- data.frame(index=index[,climate], taxon=index$taxon)
    
    # Prepare colours and order for plotting
    cols <- setNames(c("forestgreen","forestgreen","red", "red", "red", "red","red"), c("plants", "fungi", "insects", "mammals", "reptiles", "amphibians", "birds"))
    data$j.taxon <- as.numeric(factor(data$taxon))

    # Check for order of names as otherwise plots will be wrong
    if(!identical(names(cols), as.character(unique(data$taxon))))
        stop("Names of taxa in wrong order; plots will be wrong!")
    
    
    # Lookups for nicer names
    c.var <- setNames(c("cloud-cover","frost-days","potential-evapotranspiration","precipitation","min-temperature","mean-temperature","max-temperature","vapour-pressure"),
                      c("cld","frs","pet","pre","tmn","tmp","tmx","vap")
                      )[climate]
    q.var <- setNames(c("5th","25th","50th","75th","95th"), c("0.05","0.25","0.5","0.75","0.95"))[quant]

    # Calculate plot-points
    medians <- with(data, tapply(index, taxon, quantile, probs=.5))
    lower <- with(data, tapply(index, taxon, quantile, probs=.25))
    upper <- with(data, tapply(index, taxon, quantile, probs=.75))
    n.real <- with(data, tapply(index, taxon, length))
    s.medians <- with(sim.data, tapply(index, j.a.cat, quantile, probs=.5))
    s.lower <- with(sim.data, tapply(index, j.a.cat, quantile, probs=.25))
    s.upper <- with(sim.data, tapply(index, j.a.cat, quantile, probs=.75))
    n.sim <- with(sim.data, tapply(index, j.a.cat, length))

    
    par(mar=c(6.1,5.1,1.1,1.1))
    if(sims){
        xlims <- c(1,10.5)} else xlims <- c(1,7.5)
    eval(substitute(
        with(data, plot(index ~ jitter(j.taxon, factor=1.25), pch=20, col=alpha(cols,.25)[taxon], cex=.75, cex.lab=1.5, axes=FALSE, xlab="", xlim=xlims, ylab=expression(track[XXX](YYY)))), list(XXX=q.var, YYY=c.var)
    ))
    if(sims)
        with(sim.data, points(index ~ I(jitter(j.a.cat)+7), pch=20, col=alpha("blue",.25), cex=.25))
    
    
    arrows(1:7+.45, medians, 1:7+.55, col=cols, length=0, lwd=10)
    arrows(1:7+.45, lower, 1:7+.55, col=cols, length=0, lwd=5)
    arrows(1:7+.45, upper, 1:7+.55, col=cols, length=0, lwd=5)
    if(sims){
        arrows(8:10+.45, s.medians, 8:10+.55, col="blue", length=0, lwd=10)
        arrows(8:10+.45, s.lower, 8:10+.55, col="blue", length=0, lwd=5)
        arrows(8:10+.45, s.upper, 8:10+.55, col="blue", length=0, lwd=5)
    }
        
    abline(h=0, col="grey40", lwd=3, lty=2)
    abline(h=1, col="grey40", lwd=3, lty=2)
    
    # Do work
    axis(1, labels=names(cols), at=1:7, cex.axis=1.5)
    if(sims)
        axis(1, labels=unique(sim.data$alpha.cat), at=8:10, cex.axis=1.5)
    axis(2)
    if(sims){
        mtext(side=1, text=paste0("(n=",c(n.real,n.sim),")"), at=1:10, line=2)
    } else {
        mtext(side=1, text=paste0("(n=",n.real,")"), at=1:7, line=2)
    }

    # Add rasters
    if(sims){
        grid.raster(readPNG("phylopics/plant.png"),     .09, .12, width=.04, just="bottom")
        grid.raster(readPNG("phylopics/fungus.png"),    .18, .12, width=.04, just="bottom")
        grid.raster(readPNG("phylopics/insect.png"),    .27, .12, width=.02, just="bottom")
        grid.raster(readPNG("phylopics/mammal.png"),    .36, .12, width=.04, just="bottom")
        grid.raster(readPNG("phylopics/reptile.png"),   .45, .12, width=.04, just="bottom")
        grid.raster(readPNG("phylopics/amphibian.png"), .54, .12, width=.04, just="bottom")
        grid.raster(readPNG("phylopics/bird.png"),      .64, .12, width=.04, just="bottom")
    } else {
        grid.raster(readPNG("phylopics/plant.png"),     .095, .12, width=.04, just="bottom")
        grid.raster(readPNG("phylopics/fungus.png"),    .225, .12, width=.04, just="bottom")
        grid.raster(readPNG("phylopics/insect.png"),    .36, .12, width=.02, just="bottom")
        grid.raster(readPNG("phylopics/mammal.png"),    .49, .12, width=.04, just="bottom")
        grid.raster(readPNG("phylopics/reptile.png"),   .62, .12, width=.04, just="bottom")
        grid.raster(readPNG("phylopics/amphibian.png"), .76, .12, width=.04, just="bottom")
        grid.raster(readPNG("phylopics/bird.png"),      .89, .12, width=.04, just="bottom")
    }
    
}

for(climate in c("cld","frs","pet","pre","tmn","tmp","tmx","vap","wet")){
    for(quant in c("0.05","0.25","0.5","0.75","0.95")){
        pdf(paste0("figures/violin-",climate,"-",gsub(".","-",quant,fixed=TRUE),".pdf"), width=17, height=10)
        .outline.plot(climate, quant, clean=5)
        dev.off()
    }
}        
pdf(paste0("figures/violin-tmp-0-5.pdf"), width=17, height=10);.outline.plot("tmp", "0.5", clean=5, sims=TRUE);dev.off()


# Calculate the numbers
.percentage.stats <- function(index, var, clean, abs=FALSE, overshoot=FALSE, process=TRUE){
    index <- .simplify.group(index, var, clean, abs)
    
    if(overshoot){
        index <- index > 1
    } else index <- index < 1 & index > 0
    if(process){
        percentages <- table(rowSums(index)) / nrow(index)
        percentages <- percentages[order(names(percentages))]
        something <- 1-percentages[1]
        at.least.two <- 1 - sum(percentages[1:2])
        at.least.three <- 1 - sum(percentages[1:3])
        everything <- percentages[6]
        return(setNames(
            c(something,at.least.two,at.least.three, everything),
            c("something","at.least.two","at.least.three","everything")))
    } else return(index)
}
.percentage.stats("bootstrap.index", "cld", clean=100)
.percentage.stats("bootstrap.index", "frs", clean=100)
.percentage.stats("bootstrap.index", "pet", clean=100)
.percentage.stats("bootstrap.index", "pre", clean=100)
.percentage.stats("bootstrap.index", "tmn", clean=100)
.percentage.stats("bootstrap.index", "tmp", clean=100)
.percentage.stats("bootstrap.index", "tmx", clean=100)
.percentage.stats("bootstrap.index", "vap", clean=100)
.percentage.stats("bootstrap.index", "wet", clean=100)

combined <- .percentage.stats("bootstrap.index", "wet", process=FALSE, clean=100) |
    .percentage.stats("bootstrap.index", "frs", process=FALSE, clean=100) |
    .percentage.stats("bootstrap.index", "pet", process=FALSE, clean=100) |
    .percentage.stats("bootstrap.index", "pre", process=FALSE, clean=100) |
    .percentage.stats("bootstrap.index", "tmn", process=FALSE, clean=100) |
    .percentage.stats("bootstrap.index", "tmp", process=FALSE, clean=100) |
    .percentage.stats("bootstrap.index", "tmx", process=FALSE, clean=100) |
    .percentage.stats("bootstrap.index", "vap", process=FALSE, clean=100) |
    .percentage.stats("bootstrap.index", "wet", process=FALSE, clean=100)
combined <- rowSums(combined)
combined <- table(combined) / length(combined)
1 - combined["0"]

.percentage.stats("bootstrap.index", "cld", overshoot=TRUE, clean=100)
.percentage.stats("bootstrap.index", "frs", overshoot=TRUE, clean=100)
.percentage.stats("bootstrap.index", "pet", overshoot=TRUE, clean=100)
.percentage.stats("bootstrap.index", "pre", overshoot=TRUE, clean=100)
.percentage.stats("bootstrap.index", "tmn", overshoot=TRUE, clean=100)
.percentage.stats("bootstrap.index", "tmp", overshoot=TRUE, clean=100)
.percentage.stats("bootstrap.index", "tmx", overshoot=TRUE, clean=100)
.percentage.stats("bootstrap.index", "vap", overshoot=TRUE, clean=100)
.percentage.stats("bootstrap.index", "wet", overshoot=TRUE, clean=100)

o.combined <- .percentage.stats("bootstrap.index", "wet", process=FALSE, clean=100) |
    .percentage.stats("bootstrap.index", "frs", overshoot=TRUE, process=FALSE, clean=100) |
    .percentage.stats("bootstrap.index", "pet", overshoot=TRUE, process=FALSE, clean=100) |
    .percentage.stats("bootstrap.index", "pre", overshoot=TRUE, process=FALSE, clean=100) |
    .percentage.stats("bootstrap.index", "tmn", overshoot=TRUE, process=FALSE, clean=100) |
    .percentage.stats("bootstrap.index", "tmp", overshoot=TRUE, process=FALSE, clean=100) |
    .percentage.stats("bootstrap.index", "tmx", overshoot=TRUE, process=FALSE, clean=100) |
    .percentage.stats("bootstrap.index", "vap", overshoot=TRUE, process=FALSE, clean=100) |
    .percentage.stats("bootstrap.index", "wet", overshoot=TRUE, process=FALSE, clean=100)
o.combined <- rowSums(o.combined)
o.combined <- table(o.combined) / length(o.combined)
1 - o.combined["0"]


.violin.plot <- function(climate, quant, clean=NA, se.thresh=5, abs=FALSE, index="bootstrap.index"){
    # Load and match data
    index <- .simplify.group(index, quantile=quant, clean=clean, abs=abs)
    se <- .simplify.group("mad.index", quantile=quant, abs=abs)
    se <- se[se[,climate] < se.thresh,]
    se <- se[rownames(se) %in% rownames(index),]
    index <- index[rownames(index) %in% rownames(se),]
    data <- data.frame(index=index[,climate], se=se[,climate], taxon=index$taxon)

    # Prepare colours and order for plotting
    cols <- setNames(c("forestgreen","forestgreen","red", "red", "red", "red","red"), c("plants", "fungi", "insects", "mammals", "reptiles", "amphibians", "birds"))
    data$col <- alpha(cols[data$taxon], alpha=1-((data$se^2)/max(data$se^2)))
    data <- data[order(data$taxon, data$index),]
    data$x <- as.numeric(factor(data$taxon))
    counts <- table(data$taxon)
    curr <- 0
    for(i in seq_along(counts)){
        taxon <- names(counts)[i]
        data$x[data$taxon==taxon] <- data$x[data$taxon==taxon] + seq(-.3, .3, length.out=counts[i])
    }

    # Lookups for nicer names
    c.var <- setNames(c("cloud-cover","frost-days","potential-evapotranspiration","precipitation","min-temperature","mean-temperature","max-temperature","vapour-pressure"),
                      c("cld","frs","pet","pre","tmn","tmp","tmx","vap")
                      )[climate]
    q.var <- setNames(c("5th","25th","50th","75th","95th"), c("0.05","0.25","0.5","0.75","0.95"))[quant]

    # Calculate plot-points
    medians <- with(data, tapply(index, taxon, quantile, probs=.5))
    lower <- with(data, tapply(index, taxon, quantile, probs=.25))
    upper <- with(data, tapply(index, taxon, quantile, probs=.75))

    # Do work
    eval(substitute(
        with(data, plot(index ~ x, xlab="", ylab=expression(track[XXX](YYY)), axes=FALSE, pch=20, col=col)),
        list(XXX=q.var, YYY=c.var)
    ))
    arrows(1:6-.2, medians, 1:6+.2, col=cols, length=0, lwd=5)
    arrows(1:6-.2, lower, 1:6+.2, col=cols, length=0, lwd=5)
    arrows(1:6-.2, upper, 1:6+.2, col=cols, length=0, lwd=5)
    abline(h=0, col="grey40", lwd=3, lty=2)
    abline(h=1, col="grey40", lwd=3, lty=2)
    axis(1, labels=names(cols), at=1:6)    
    axis(2)
}
```
