## supplemental code for "The realised velocity of climate change reveals remarkable idiosyncrasy of species’ distributional shifts": calculate-index.R

```
# Headers
source("src/headers.R")

# Wrapper functions
.calc.track.parallel <- function(data, quantiles, rasters, taxon){
    .int.calc <- function(sp){
        # Setup and subset data
        curr.data <- data[scientificname == sp,]
        post.data <- curr.data[year %in% 1990:2015,]
        pre.data <- curr.data[year %in% 1955:1980,]
        post.years <- unique(post.data$year)
        output <- array(
            dim=c(length(rasters), length(quantiles), 7),
            dimnames=list(names(rasters), as.character(quantiles), c("index","bootstrap.index", "mad.index", "quantile","present","past","projected"))
        )
        if(nrow(pre.data) == 0 | nrow(post.data) == 0)
            return(FALSE)        
        
        # Loop over rasters
        for(i in seq_along(rasters)){
            # Prepare to store data
            post.data$env <- -999
            pre.data$pre.env <- -999
            pre.data$post.env <- -999
            pre.years <- unique(pre.data$year)

            # Match climate data (in chunks to save memory and for speed)
            for(k in seq_along(post.years))
                post.data[year==post.years[k],]$env <- extract(rasters[[i]][[post.data$year[k]-1900]], post.data[year==post.years[k],.(decimallongitude,decimallatitude)])
            for(k in seq_along(pre.years)){
                pre.data[year==pre.years[k],]$pre.env <- extract(rasters[[i]][[pre.data$year[k]-1900]], pre.data[year==pre.years[k],.(decimallongitude,decimallatitude)])
                pre.data[year==pre.years[k],]$post.env <- extract(rasters[[i]][[pre.data$year[k]-1900+25]], pre.data[year==pre.years[k],.(decimallongitude,decimallatitude)])
            }

            # Do work
            output[i,,] <- t(sapply(quantiles, function(x) .calc.metric(pre.data$pre.env, post.data$env, pre.data$post.env, x)))
        }

        saveRDS(output, paste0(output.dir,taxon,"/",sp,".RDS"))
        return(TRUE)
    }
    
    mcMap(.int.calc, unique(data$scientificname))
    return(TRUE)
}
.merge.results <- function(taxon){
    files <- list.files(paste0(output.dir,taxon))
    results <- Map(function(x) readRDS(paste0(output.dir,taxon,"/",x)), files)
    names(results) <- gsub(".RDS", "", names(results), fixed=TRUE)
    return(abind(results, along=4))
}

# Load and clean GBIF data
plants <- .load.gbif(
    obs=paste0(gbif.dir,"plantae/0001849-180131172636756_abbreviated.txt")#,
    #specimens=paste0(gbif.dir,"plantae/0001851-180131172636756_abbreviated.txt")
, min.records=1000, clean.binomial=TRUE)
mammals <- .load.gbif(obs=paste0(gbif.dir,"mammalia/0000171-180412121330197_abbreviated.txt"), min.records=1000, clean.binomial=TRUE)
amphibians <- .load.gbif(obs=paste0(gbif.dir,"amphibia/0034734-180508205500799_abbreviated.txt"), min.records=1000, clean.binomial=TRUE)
fungi <- .load.gbif(obs=paste0(gbif.dir,"asco-basidio/0034724-180508205500799_abbreviated.txt"), min.records=1000, clean.binomial=TRUE)
birds <- .load.gbif(obs=paste0(gbif.dir,"aves/0000172-180412121330197_abbreviated.txt"), min.records=1000, clean.binomial=TRUE)
insects <- .load.gbif(obs=paste0(gbif.dir,"insecta/0034726-180508205500799_abbreviated.txt"), min.records=1000, clean.binomial=TRUE)
reptiles <- .load.gbif(obs=paste0(gbif.dir,"reptillia/0034728-180508205500799_abbreviated.txt"), min.records=1000, clean.binomial=TRUE)

# Load climate rasters
cru <- list(
    cld=readRDS(paste0(output.dir,"cru-cld.RDS")),
    frs=readRDS(paste0(output.dir,"cru-frs.RDS")),
    pet=readRDS(paste0(output.dir,"cru-pet.RDS")),
    pre=readRDS(paste0(output.dir,"cru-pre.RDS")),
    tmn=readRDS(paste0(output.dir,"cru-tmn.RDS")),
    tmp=readRDS(paste0(output.dir,"cru-tmp.RDS")),
    tmx=readRDS(paste0(output.dir,"cru-tmx.RDS")),
    vap=readRDS(paste0(output.dir,"cru-vap.RDS")),
    wet=readRDS(paste0(output.dir,"cru-wet.RDS"))
)    
    
# Do work and save results
print("mammals")
.calc.track.parallel(mammals, quantiles, cru, "mammals")
print("amphibians")
.calc.track.parallel(amphibians, quantiles, cru, "amphibians")
print("fungi")
.calc.track.parallel(fungi, quantiles, cru, "fungi")
print("insects")
.calc.track.parallel(insects, quantiles, cru, "insects")
print("reptiles")
.calc.track.parallel(reptiles, quantiles, cru, "reptiles")
print("plants")
.calc.track.parallel(plants, quantiles, cru, "plants")
print("birds")
.calc.track.parallel(birds, quantiles, cru, "birds")

# Merge the results back together again
saveRDS(.merge.results("mammals"), paste0(output.dir,"mammals-index.RDS"))
saveRDS(.merge.results("amphibians"), paste0(output.dir,"amphibians-index.RDS"))
saveRDS(.merge.results("fungi"), paste0(output.dir,"fungi-index.RDS"))
saveRDS(.merge.results("insects"), paste0(output.dir,"insects-index.RDS"))
saveRDS(.merge.results("reptiles"), paste0(output.dir,"reptiles-index.RDS"))
saveRDS(.merge.results("plants"), paste0(output.dir,"plants-index.RDS"))
saveRDS(.merge.results("birds"), paste0(output.dir,"birds-index.RDS"))
```
