## supplemental code for "The realised velocity of climate change reveals remarkable idiosyncrasy of species’ distributional shifts": check-index.R

```
# Load data
source("headers.R")
temp <- readRDS("../clean-data/temp.RDS")

# Wrapper functions
.make.dist <- function(x, y, range)
    return(as.matrix(expand.grid(x+seq(-range,range), y+seq(-range,range))))
.check.viable <- function(dist, raster)
    !(any(dist < 0) | any(dist[,1] > nrow(raster)) | any(dist[,2] > ncol(raster)))
.assign.outcome <- function(present, past, projected){
    if(present==past)
        return("track")
    if(present==projected)
        return("static-distribution")

    # Warming
    if(projected > past){
        if(present < projected){
            if(present > past)
                return("lag")
            return("overshoot")
        }
        if(present > projected){
            return("undershoot")
        }
    }
    
    # Cooling
    if(projected < past){
        if(present > projected){
            if(present < past)
                return("overshoot")
            return("lag")
        }
        if(present < projected){
            return("undershoot")
        }
    }
    return("ERROR")
}

# Prepare pre- and post- data
early <- as.matrix(temp[[1]])
mid <- as.matrix(temp[[50]])
late <- as.matrix(temp[[100]])

# Select viable starting distributions
centers <- which(!is.na(as.matrix(early)), arr.ind=TRUE)
centers <- centers[sample(nrow(centers), 100),]
viable <- rep(FALSE, nrow(centers))
for(i in seq_along(viable))
    viable[i] <- .check.viable(.make.dist(centers[i,1], centers[i,2], 10), early)
centers <- centers[viable,]

# Add parameters for simulations
species <- expand.grid(shift=seq(-2,2), range=c(1,2,5), merge=paste(centers[,1], centers[,1], sep="-"))
centers <- data.frame(x=centers[,1], y=centers[,2], merge=paste(centers[,1], centers[,1], sep="-"))
data <- merge(species, centers, by="merge")

# Run simulation
mid.index <- late.index <- summaries <- vector("list", nrow(data))
for(i in seq_len(nrow(data))){
    prog.bar(i, nrow(data))
    # Make distributions
    early.dist <- .make.dist(data$x[i], data$y[i], data$range[i])
    mid.dist <- .make.dist(data$x[i], data$y[i]+data$shift[i], data$range[i])
    late.dist <- .make.dist(data$x[i], data$y[i]+data$shift[i]*2, data$range[i])
    # Calculate index
    mid.index[[i]] <- .calc.metric(as.numeric(mid[mid.dist]), as.numeric(early[early.dist]), as.numeric(mid[early.dist]), quantile=.5, n.boot=99)
    late.index[[i]] <- .calc.metric(as.numeric(late[late.dist]), as.numeric(early[early.dist]), as.numeric(late[early.dist]), quantile=.5, n.boot=99)
}

# Some post-processing
for(i in seq_along(late.index)){
    if(is.null(late.index[[i]]))
        late.index[[i]] <- setNames(rep(NA, 7), c("index","bootstrap.index", "mad.index", "quantile","present","past","projected"))
    if(is.null(mid.index[[i]]))
        mid.index[[i]] <- setNames(rep(NA, 7), c("index","bootstrap.index", "mad.index", "quantile","present","past","projected"))
}
late.index <- do.call(rbind, late.index)
late.index <- cbind(data, late.index)
mid.index <- do.call(rbind, mid.index)
mid.index <- cbind(data, mid.index)
late.index <- na.omit(late.index)
mid.index <- na.omit(mid.index)

# Assign what happened
mid.index$outcome <- with(mid.index, mapply(.assign.outcome, present, past, projected))
late.index$outcome <- with(late.index, mapply(.assign.outcome, present, past, projected))
outcome.col <- setNames(c("forestgreen","blue","grey60","red"), c("lag","overshoot","static-distribution","undershoot"))

# "Remove" outliers
mid.index$backup <- mid.index$index
late.index$backup <- late.index$index
mid.index$index[abs(mid.index$index) > 50] <- NA
late.index$index[abs(late.index$index) > 50] <- NA

#mid.index$jitter <- jitter(mid.index$shift)
mid.index$c.change <- with(mid.index, ifelse(present-past > 0, "red", "blue"))
mid.index$c.change[mid.index$shift==0] <- "black"

mid.index <- mid.index[order(mid.index$shift, mid.index$index),]
unique.shifts <- sort(unique(mid.index$shift))
mid.index$jitter <- -1
for(i in seq_along(unique.shifts)){
    which <- which(mid.index$shift==unique.shifts[i])
    mid.index$jitter[which] <- mid.index$shift[which] + seq(-.4,.4, length.out=length(which))
}

pdf("null-movement.pdf", width=10, height=5)
with(mid.index, plot(bootstrap.index ~ jitter, ylim=c(-10,10), pch=20, col=outcome.col[outcome], axes=FALSE, xlab="Grid cell shift", ylab="Bootstrapped index"))
axis(1)
axis(2)
with(mid.index, arrows(x0=jitter, y0=bootstrap.index-mad.index, y1=bootstrap.index+mad.index, col=outcome.col[outcome], lwd=.5, code=3, angle=0))
abline(h=1, col="grey60", lwd=3)
abline(h=0, col="forestgreen", lwd=3)
legend("topright", pch=20, col=outcome.col, legend=names(outcome.col), bty="n")
dev.off()

with(mid.index, summary(lm(index ~ outcome -1)))
```
