## supplemental code for "The realised velocity of climate change reveals remarkable idiosyncrasy of species’ distributional shifts": download-cru.rb

```
#!/usr/bin/ruby

# Change the location (and its match in `headers.R`) to download CRU
# data elsewhere
Dir.chdir "/home/will/bear-share/cru/"

def download_wrapper(cru_id, var, stem="cruts.1905011326.v4.03", delay=60)
  sleep delay
  `wget https://crudata.uea.ac.uk/cru/data/hrg/cru_ts_4.03/#{stem}/#{var}/#{cru_id}`
  `gunzip #{cru_id}`
end

#download_wrapper("cru_ts4.03.1901.2018.cld.dat.nc.gz", "cld")
#download_wrapper("cru_ts4.03.1901.2018.dtr.dat.nc.gz", "dtr")
#download_wrapper("cru_ts4.03.1901.2018.frs.dat.nc.gz", "frs")
download_wrapper("cru_ts4.03.1901.2018.pet.dat.nc.gz", "pet")
download_wrapper("cru_ts4.03.1901.2018.pre.dat.nc.gz", "pre")
download_wrapper("cru_ts4.03.1901.2018.tmn.dat.nc.gz", "tmn")
download_wrapper("cru_ts4.03.1901.2018.tmp.dat.nc.gz", "tmp")
download_wrapper("cru_ts4.03.1901.2018.tmx.dat.nc.gz", "tmx")
download_wrapper("cru_ts4.03.1901.2018.vap.dat.nc.gz", "vap")
download_wrapper("cru_ts4.03.1901.2018.wet.dat.nc.gz", "wet")
```
