## supplemental code for "The realised velocity of climate change reveals remarkable idiosyncrasy of species’ distributional shifts": download-gbif.rb

```
#!/usr/bin/ruby

# Change the location (and its match in `headers.R`) to download GBIF
# data elsewhere

Dir.chdir "~/bear-share/gbif/"
def download_wrapper(gbif_id, taxon, clean_csv=true, clean_zip=true, clean_gbif=false)
  Dir.chdir taxon
  `wget http://api.gbif.org/v1/occurrence/download/request/#{gbif_id}.zip`
  `unzip #{gbif_id}.zip #{gbif_id}.csv`
  `cut -f13,17,18,28 #{gbif_id}.csv > #{gbif_id}_abbreviated.txt`
  `cut -f13,17,18,28,44 #{gbif_id}.csv > #{gbif_id}_short_issues.txt`
  if clean_csv then `rm #{gbif_id}.csv` end
  if clean_zip then `rm #{gbif_id}.zip` end
  Dir.chdir '..'
end

# If you want to use different GBIF inputs for each taxon, replace the
# IDs below with something different. You will then need to change
# `calculate-index.R` since the raw data will have changed
download_wrapper "0034734-180508205500799" "amphibia"
download_wrapper "0034724-180508205500799" "asco-basidio"
download_wrapper "0000172-180412121330197" "aves"
download_wrapper "0034726-180508205500799" "insecta"
download_wrapper "0000171-180412121330197" "mammalia"
download_wrapper "0001849-180131172636756" "plantae"
download_wrapper "0001851-180131172636756" "plantae"
download_wrapper "0034728-180508205500799" "reptillia"
```
