## supplemental code for "The realised velocity of climate change reveals remarkable idiosyncrasy of species’ distributional shifts": headers.R

```
####################################
# Packages #########################
####################################
if(!require("data.table")){
    install.packages("data.table"); library(data.table)}
if(!require("raster")){
    install.packages("raster"); library(raster)}
if(!require("RNetCDF")){
    install.packages("RNetCDF"); library(RNetCDF)}
if(!require("parallel")){
    install.packages("parallel"); library(parallel)}
if(!require("phytools")){
    install.packages("phytools"); library(phytools)}
if(!require("caper")){
    install.packages("caper"); library(caper)}
if(!require("abind")){
    install.packages("abind"); library(abind)}
if(!require("ks")){
    install.packages("ks"); library(ks)}
if(!require("geiger")){
    install.packages("geiger"); library(geiger)}
if(!require("suppdata")){
    install.packages("suppdata"); library(suppdata)}
if(!require("pez")){
    install.packages("pez"); library(pez)}
if(!require("scales")){
    install.packages("scales"); library(scales)}
if(!require("reldist")){
    install.packages("reldist"); library(reldist)}
if(!require("gdata")){
    install.packages("gdata"); library(gdata)}
if(!require("corrplot")){
    install.packages("corrplot"); library(corrplot)}
if(!require("viridis")){
    install.packages("viridis"); library(viridis)}
if(!require("plotrix")){
    install.packages("plotrix"); library(plotrix)}
if(!require("png")){
    install.packages("png"); library(png)}
if(!require("grid")){
    install.packages("grid"); library(grid)}

####################################
# Global settings ##################
####################################
gbif.dir <- "/home/will/bear-share/gbif/"
if(gbif.dir!="/home/will/bear-share/gbif/")
    warning("GBIF data is not in default directory: ensure `download-gbif.rb` is similarly updated")

cru.dir <- "/home/will/bear-share/cru/"
if(cru.dir!="/home/will/bear-share/cru/")
    warning("CRU data is not in default directory: ensure `download-cru.rb` is similarly updated")

output.dir <- "clean-data/"

quantiles <- c(.05, .25, .5, .75, .95)

options(mc.cores=24)
if(options("mc.cores")==24)
    warning("Assuming you have 24 cores on your computer: if not, change `headers.R`")

####################################
# Functions ########################
####################################
prog.bar <- function(x, y){
    if(y < 100){
        cat(".")} else {
            z <- Filter(function(z) z>=0, seq(1,y,length.out=100)-x)
            if(length(z) > 0)
                tryCatch(if(z[1] < 1) if((length(z) %% 10)==0) cat("|") else cat("."), error=function(z) cat("."))
        }
}    
.load.gbif <- function(obs, specimens, min.records=2000, clean.binomial=FALSE, clean.gbif=FALSE){
    data <- data.table(scientificname="", decimallatitude=0.0, decimallongitude=0.0, year=0, type="")
    if(clean.gbif)
        data$issue <- ""
    if(!missing(obs)){
        tmp <- fread(obs)
        tmp$type <- "observations"
        data <- rbind(data, tmp)
    }
    if(!missing(specimens)){
        tmp <- fread(specimens)
        tmp$type <- "specimens"
        data <- rbind(data, tmp)
    }
    if(missing(specimens) & missing(obs))
        stop("Must supply *some* data to load!...")
    data <- data[-1,]

    if(clean.gbif)
        data <- data[issue == "",]
    
    if(clean.binomial){
        spp <- unique(data$scientificname)
        spp <- setNames(tolower(sapply(strsplit(spp, " "), function(x) paste(x[1:2], collapse="_"))), spp)
        data$scientificname <- spp[data$scientificname]
    }

    # Find most common species and subset
    spp.table <- sort(table(data$scientificname), TRUE)
    spp.table <- names(spp.table)[spp.table >= min.records]
    data <- data[scientificname %in% spp.table,]

    # Subset by year
    data <- data[year > 1955 & year <= 2015,]
    return(data)
}
.calc.track <- function(i){
    name <- paste0("taxon_", i, "_results.RDS")
    data <- wrap.list[[i]]
    quantiles <- seq(.1,.9,.05)
    results <- array(dim=c(length(unique(data$scientificname)),ncol=14,length(quantiles)), dimnames=list(unique(data$scientificname), c("index","rank.pres","rank.past"), quantiles))
    for(i in seq_len(nrow(results))){
        # Pre-allocation and calculation
        curr.data <- data[scientificname == rownames(results)[i],]
        post.data <- curr.data[year %in% 1990:2015,]
        pre.data <- curr.data[year %in% 1955:1980,]
        post.years <- unique(post.data$year)
        post.data$env <- -999
        for(k in seq_along(post.years))
            post.data[year==post.years[k],]$env <- extract(temp[[post.data$year[k]-1900]], post.data[year==post.years[k],.(decimallongitude,decimallatitude)])

        # Chuck out if potential sampling problems
        if(nrow(pre.data) == 0 | nrow(post.data) == 0)
            next
        
        # Past and projected data extraction
        pre.data$pre.env <- -999
        pre.data$post.env <- -999
        pre.years <- unique(pre.data$year)
        for(k in seq_along(pre.years)){
            pre.data[year==pre.years[k],]$pre.env <- extract(temp[[pre.data$year[k]-1900]], pre.data[year==pre.years[k],.(decimallongitude,decimallatitude)])
            pre.data[year==pre.years[k],]$post.env <- extract(temp[[pre.data$year[k]-1900+25]], pre.data[year==pre.years[k],.(decimallongitude,decimallatitude)])
        }

        # Calculate for all quantiles
        results[i,,] <- sapply(quantiles, function(x) .calc.metric(pre.data$pre.env, post.data$env, pre.data$post.env, x))
    }
    saveRDS(results, name)
    return(results)
}
.calc.metric <- function(present, past, projected, quantile=.5, n.boot=999){
    # Calculate observed quantiles
    m.present <- quantile(present, quantile, na.rm=TRUE)
    m.past <- quantile(past, quantile, na.rm=TRUE)
    m.projected <- quantile(projected, quantile, na.rm=TRUE)
    index <- (m.present-m.past) / (m.projected-m.past)

    # Bootstrap past and present sampling
    b.present <- replicate(n.boot, sample(present, length(past), replace=TRUE), simplify=FALSE)
    b.present <- sapply(b.present, quantile, probs=quantile, na.rm=TRUE)
    b.past <- replicate(n.boot, sample(past, length(present), replace=TRUE), simplify=FALSE)
    b.past <- sapply(b.past, quantile, probs=quantile, na.rm=TRUE)
    b.projected <- replicate(n.boot, sample(projected, length(present), replace=TRUE), simplify=FALSE)
    b.projected <- sapply(b.projected, quantile, probs=quantile, na.rm=TRUE)
    b.mad <- mad((b.present-b.past) / (b.projected-b.past))
    b.index <- median((b.present-b.past) / (b.projected-b.past))
    
    # Format and return
    return(setNames(
        c(index, b.index, b.mad, quantile,m.present,m.past,m.projected),
        c("index","bootstrap.index", "mad.index", "quantile","present","past","projected")
    ))
}
.assign.outcome <- function(present, past, projected){
    if(present==past)
        return("track")
    if(present==projected)
        return("static-distribution")

    # Warming
    if(projected > past){
        if(present < projected){
            if(present > past)
                return("lag")
            return("overshoot")
        }
        if(present > projected){
            return("undershoot")
        }
    }
    
    # Cooling
    if(projected < past){
        if(present > projected){
            if(present < past)
                return("lag")
            return("overshoot")
        }
        if(present < projected){
            return("undershoot")
        }
    }
    return("ERROR")
}
.simplify <- function(data, index, quantile, clean=NA, abs=FALSE){
    data <- t(data[,quantile,index,])
    if(!is.na(clean)){
        keep <- rep(TRUE, nrow(data))
        for(i in seq_len(ncol(data)))
            keep <- keep & abs(data[,i]) <= clean & (!is.na(data[,i]))
        data <- data[keep,]
    }
    if(abs)
        data <- abs(data)
    return(data)
}
.simplify.group <- function(index, quantile, clean=NA, abs=FALSE)
    return(rbind(
        data.frame(.simplify(plants, index, quantile, clean, abs), taxon="plants"),
        data.frame(.simplify(fungi, index, quantile, clean, abs), taxon="fungi"),
        data.frame(.simplify(insects, index, quantile, clean, abs), taxon="insects"),
        data.frame(.simplify(mammals, index, quantile, clean, abs), taxon="mammals"),
        data.frame(.simplify(reptiles, index, quantile, clean, abs), taxon="reptiles"),
        data.frame(.simplify(amphibians, index, quantile, clean, abs), taxon="amphibians"),
        data.frame(.simplify(birds, index, quantile, clean, abs), taxon="birds")
    ))
.simplify.quantile <- function(index, var, clean=NA, abs=FALSE)
    return(rbind(
        data.frame(.simplify(plants, index, quantile, clean, abs), taxon="plants"),
        data.frame(.simplify(fungi, index, quantile, clean, abs), taxon="fungi"),
        data.frame(.simplify(insects, index, quantile, clean, abs), taxon="insects"),
        data.frame(.simplify(mammals, index, quantile, clean, abs), taxon="mammals"),
        data.frame(.simplify(reptiles, index, quantile, clean, abs), taxon="reptiles"),
        data.frame(.simplify(amphibians, index, quantile, clean, abs), taxon="amphibians"),
        data.frame(.simplify(birds, index, quantile, clean, abs), taxon="birds")
    ))
```
