## supplemental code for "The realised velocity of climate change reveals remarkable idiosyncrasy of species’ distributional shifts": process-cru.R

```
# Headers
source("src/headers.R")

# Wrapper functions
.create.raster.stack <- function(x, long=seq(-179.75,179.75,by=0.5), lat=seq(-89.75,89.75,by=0.5)){
    output <- raster(t(x[,,1][,ncol(x):1]), xmn=min(long), xmx=max(long), ymn=min(lat), ymx=max(lat))
    base <- raster(xmn=min(long), xmx=max(long), ymn=min(lat), ymx=max(lat))
    for(i in seq(2, dim(x)[3]))
        output <- stack(output, raster(t(x[,,i][,ncol(x):1]), xmn=min(long), xmx=max(long), ymn=min(lat), ymx=max(lat)))
    return(output)
}
.clean.cru <- function(file, var){
    nc.raw <- read.nc(open.nc(file))
    output <- array(dim=c(dim(nc.raw[[var]])[1:2], length(1901:2018)))
    for(i in seq_len(dim(output)[3])){
        prog.bar(i, dim(output)[3])
        curr <- seq(12*(i-1)+1, by=1, length.out=12)
        output[,,i] <- apply(nc.raw[[var]][,,curr], 1:2, mean, na.rm=TRUE)
    }
    return(.create.raster.stack(output))
}

# Do work
saveRDS(
    .clean.cru(paste0(cru.dir,"cru_ts4.03.1901.2018.cld.dat.nc"), "cld"),
    paste0(output.dir,"cru-cld.RDS")
)
saveRDS(
    .clean.cru(paste0(cru.dir,"cru_ts4.03.1901.2018.dtr.dat.nc"), "dtr"),
    paste0(output.dir,"cru-dtr.RDS")
)
saveRDS(
    .clean.cru(paste0(cru.dir,"cru_ts4.03.1901.2018.frs.dat.nc"), "frs"),
    paste0(output.dir,"cru-frs.RDS")
)
saveRDS(
    .clean.cru(paste0(cru.dir,"cru_ts4.03.1901.2018.pet.dat.nc"), "pet"),
    paste0(output.dir,"cru-pet.RDS")
)
saveRDS(
    .clean.cru(paste0(cru.dir,"cru_ts4.03.1901.2018.pre.dat.nc"), "pre"),
    paste0(output.dir,"cru-pre.RDS")
)
saveRDS(
    .clean.cru(paste0(cru.dir,"cru_ts4.03.1901.2018.tmn.dat.nc"), "tmn"),
    paste0(output.dir,"cru-tmn.RDS")
)
saveRDS(
    .clean.cru(paste0(cru.dir,"cru_ts4.03.1901.2018.tmp.dat.nc"), "tmp"),
    paste0(output.dir,"cru-tmp.RDS")
)
saveRDS(
    .clean.cru(paste0(cru.dir,"cru_ts4.03.1901.2018.tmx.dat.nc"), "tmx"),
    paste0(output.dir,"cru-tmx.RDS")
)
saveRDS(
    .clean.cru(paste0(cru.dir,"cru_ts4.03.1901.2018.vap.dat.nc"), "vap"),
    paste0(output.dir,"cru-vap.RDS")
)
saveRDS(
    .clean.cru(paste0(cru.dir,"cru_ts4.03.1901.2018.wet.dat.nc"), "wet"),
    paste0(output.dir,"cru-wet.RDS")
)
```
