## supplemental code for "The realised velocity of climate change reveals remarkable idiosyncrasy of species’ distributional shifts": sim-range.R

```
# Load data
source("src/headers.R")
temp <- readRDS("clean-data/cru-tmp.RDS")

# Wrapper functions
.make.block.dist <- function(x, y, range, raster){
    block <- as.matrix(expand.grid(x+seq(-range,range), y+seq(-range,range)))
    if(!missing(raster)){
        block <- block[block[,1] > 0 & block[,1] <= nrow(raster),]
        block <- block[block[,2] > 0 & block[,2] <= ncol(raster),]
    }
    return(block)
}
.check.viable <- function(dist, raster)
    !(any(dist < 0) | any(dist[,1] > nrow(raster)) | any(dist[,2] > ncol(raster)))
.get.niche <- function(dist, raster)
    median(raster[dist], na.rm=TRUE)
.prob.occur <- function(dist, raster, alpha, sd, optimum, sigma){
    env <- raster[dist]
    niche <- dnorm(env, optimum, sd) / dnorm(optimum, optimum, sd)
    niche <- ((1-alpha)*sigma) + (niche*alpha)
    present <- dist[runif(length(env)) <= niche,]
    return(present)
}
.dist.wrap <- function(x, y, range, raster, alpha, sd, optimum, sigma){
    block <- .make.block.dist(x, y, range, raster)
    return(.prob.occur(block, raster, alpha, sd, optimum, sigma))
}
.do.calc <- function(i){
    locker <- FALSE
    tryCatch({
        # Make distributions
        optimum <- .get.niche(.make.block.dist(data$x[i], data$y[i], data$range[i], temp), early)
        early.dist <- .dist.wrap(data$x[i], data$y[i], data$range[i], early, data$alpha[i], 1, optimum, data$sigma[i])
        mid.dist <- .dist.wrap(data$x[i], data$y[i], data$range[i], mid, data$alpha[i], 1, optimum, data$sigma[i])
        late.dist <- .dist.wrap(data$x[i], data$y[i], data$range[i], late, data$alpha[i], 1, optimum, data$sigma[i])
        # Calculate index
        mid.index <- .calc.metric(as.numeric(mid[mid.dist]), as.numeric(early[early.dist]), as.numeric(mid[early.dist]), quantile=.5, n.boot=99)
        late.index <- .calc.metric(as.numeric(late[late.dist]), as.numeric(early[early.dist]), as.numeric(late[early.dist]), quantile=.5, n.boot=99)
        locker <- TRUE
    }, error=function(x) "got a species with no distribution")
    
    # Check for successful completion and return
    if((!locker) || is.null(mid.index) || is.null(late.index))
        return(matrix(NA, 2, 7, dimnames=list(c("mid","late"), c("index","bootstrap.index", "mad.index", "quantile","present","past","projected"))))
    return(matrix(c(mid.index,late.index), 2, 7, dimnames=list(c("mid","late"), c("index","bootstrap.index", "mad.index", "quantile","present","past","projected")), byrow=TRUE))
}    

# Prepare pre- and post- data
early <- as.matrix(temp[[62]])
mid <- as.matrix(temp[[102]])
late <- as.matrix(temp[[100]])

# Select viable starting distributions
centers <- which(!is.na(as.matrix(early)), arr.ind=TRUE)
centers <- centers[sample(nrow(centers), 100),]
viable <- rep(FALSE, nrow(centers))
for(i in seq_along(viable))
    viable[i] <- .check.viable(.make.block.dist(centers[i,1], centers[i,2], 10), early)
centers <- centers[viable,]

# Add parameters for simulations
species <- expand.grid(shift=c(-4,-2,0,2,4), range=c(2,5,10,20), alpha=c(0,.5,1), merge=paste(centers[,1], centers[,1], sep="-"), sigma=c(.5,.75,1))
centers <- data.frame(x=centers[,1], y=centers[,2], merge=paste(centers[,1], centers[,1], sep="-"))
data <- merge(species, centers, by="merge")

# Do work
results <- mcMap(.do.calc, seq_len(nrow(data)))
save.image("sim-range.RData")


if(FALSE){
# Merge data
mid <- t(sapply(results, function(x) x[1,]))
data <- cbind(data, mid)
data <- data[!is.na(data$bootstrap.index),]
data <- data[is.finite(data$index),]

# Assign what happened
data$outcome <- with(data, mapply(.assign.outcome, present, past, projected))
outcome.col <- setNames(c("forestgreen","red","grey60","orange","forestgreen"), c("lag","overshoot","static-distribution","undershoot","track"))

# Summary plot
pdf("sim-range.pdf", width=10, height=10)
par(mar=c(c(5,5,2,2)))
with(data, plot(bootstrap.index ~ jitter(alpha), pch=20, col=outcome.col[outcome], cex=.5, xlab=expression(alpha), ylab="", cex.lab=2, axes=FALSE))
axis(1, cex.axis=1.25)
axis(2, at=-1:2, cex.axis=1.25)
abline(h=0, lwd=3, lty=2)
abline(h=1, lwd=3, lty=2)
mtext("(static)", side=2, at=1, line=2, cex=1.5)
mtext("(track)", side=2, at=0, line=2, cex=1.5)
mtext("(neutral)", side=1, at=0, line=2, cex=1.5)
mtext("(track)", side=1, at=1, line=2, cex=1.5)
medians <- with(data, tapply(bootstrap.index, alpha, median, na.rm=TRUE))
mads <- with(data, tapply(bootstrap.index, alpha, mad, na.rm=TRUE))
arrows(seq(0,1,.1)-.05, medians, seq(0,1,.1)+.05, code=3, length=0, lwd=5)
arrows(seq(0,1,.1), medians-mads, seq(0,1,.1), medians+mads, code=3, length=0, lwd=5)
legend("bottomleft", pch=20, col=c("grey60","forestgreen","red","orange"), legend=c("stationary","track/lag","overshoot","undershoot"), title="Range shift", bty="n", cex=1.5)
dev.off()

with(data, plot(bootstrap.index ~ jitter(alpha), pch=20, col="grey40", cex=.5))

model <- lm(bootstrap.index ~ alpha + shift + range, data=data)
summary(model)

with(data, tapply(bootstrap.index, alpha, median, na.rm=TRUE))

with(data, plot(bootstrap.index ~ jitter(alpha), pch=20, col=outcome.col[outcome]))


# "Remove" outliers
data$backup <- data$index
data$index[abs(data$index) > 50] <- NA

#mid.index$jitter <- jitter(mid.index$shift)
data$c.change <- with(data, ifelse(present-past > 0, "red", "blue"))
data$c.change[data$shift==0] <- "black"

data <- data[order(data$shift, data$bootstrap.index),]
unique.shifts <- sort(unique(data$shift))
data$jitter <- -1
for(i in seq_along(unique.shifts)){
    which <- which(data$shift==unique.shifts[i])
    data$jitter[which] <- data$shift[which] + seq(-.4,.4, length.out=length(which))
}

.plot <- function(subset.data, ...){
    subset.data <- subset.data[order(subset.data$shift, subset.data$index),]
    unique.shifts <- sort(unique(subset.data$shift))
    subset.data$jitter <- -1
    for(i in seq_along(unique.shifts)){
        which <- which(subset.data$shift==unique.shifts[i])
        subset.data$jitter[which] <- subset.data$shift[which] + seq(-.4,.4, length.out=length(which))
    }
    
    with(subset.data, plot(index ~ jitter, ylim=c(-10,10), pch=20, col=outcome.col[outcome], axes=FALSE, xlab="Grid cell shift", ylab="Bootstrapped index", ...))
    axis(1)
    axis(2)
    with(subset.data, arrows(x0=jitter, y0=bootstrap.index-mad.index, y1=bootstrap.index+mad.index, col=outcome.col[outcome], lwd=.5, code=3, angle=0))
    abline(h=1, col="grey60", lwd=3)
    abline(h=0, col="forestgreen", lwd=3)
    legend("topright", pch=20, col=outcome.col, legend=names(outcome.col), bty="n")    
}

pdf("null-movement.pdf", width=10, height=15)
par(mfcol=c(3,1))
.plot(data[data$alpha==0,], main="alpha=0")
.plot(data[data$alpha==.5,], main="alpha=0.5")
.plot(data[data$alpha==1,], main="alpha=1.0")
dev.off()

with(data, summary(lm(index ~ outcome -1)))
}
```
