## Supplementary figures and images for "The realised velocity of climate change reveals remarkable idiosyncrasy of species’ distributional shifts"

### amphibian.png

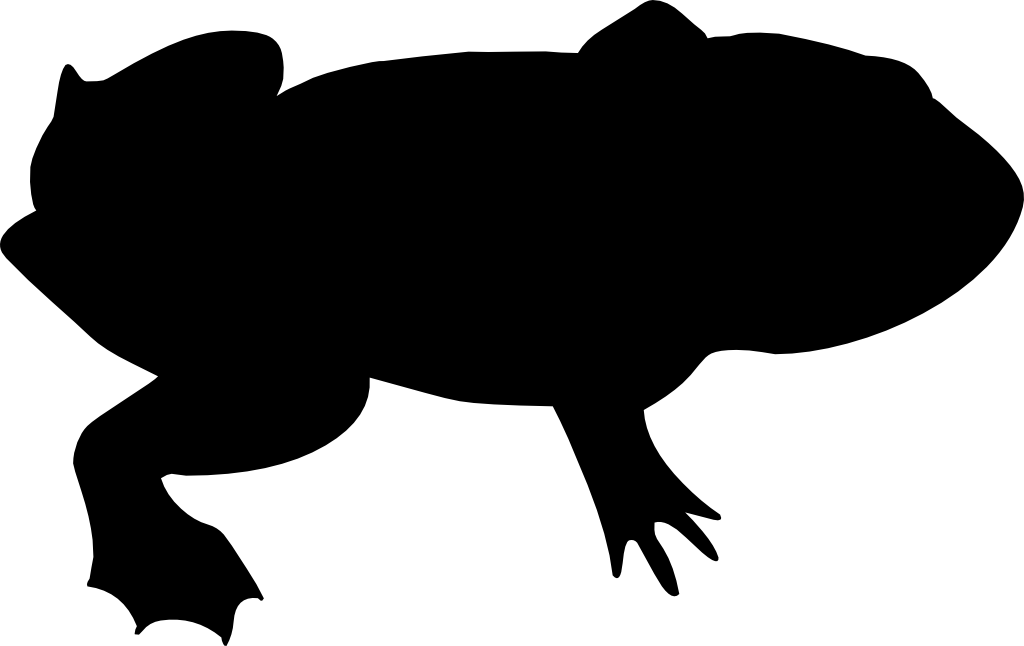

### bird.png

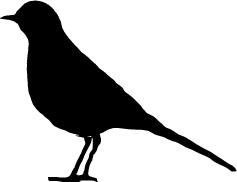

### fungus.png

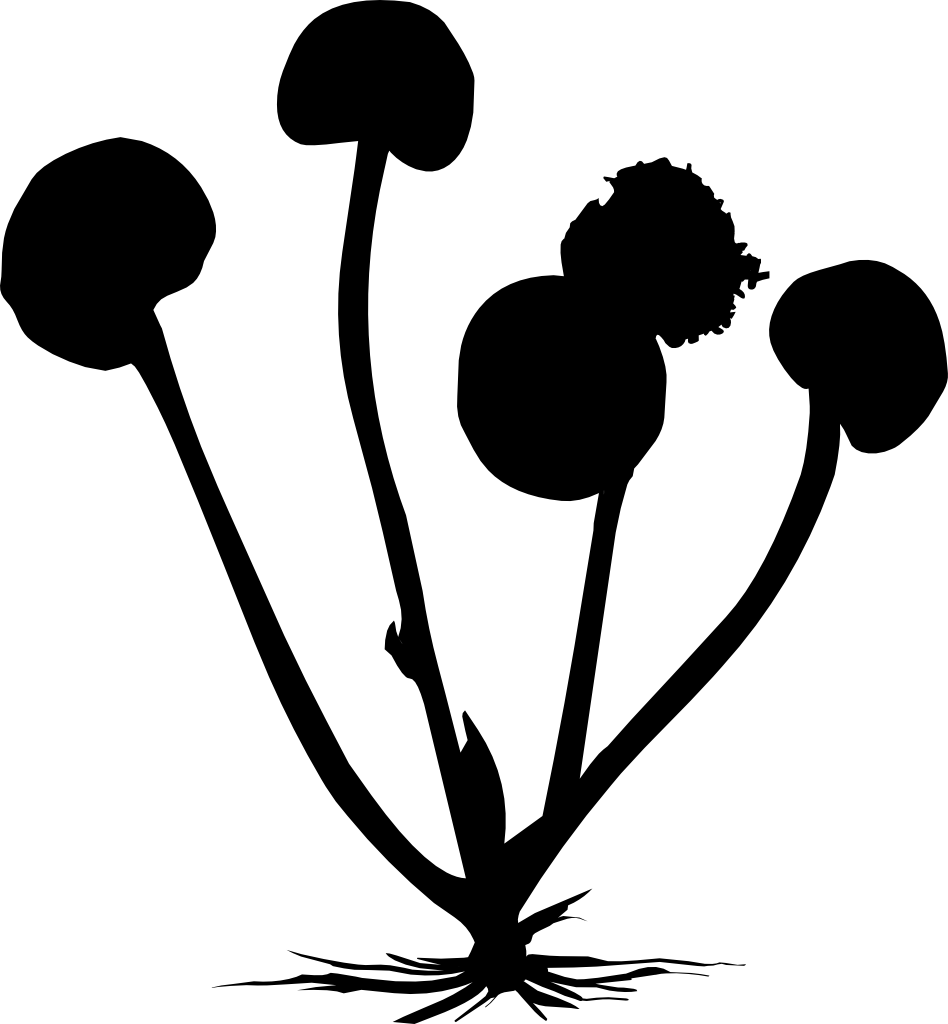

### insect.png

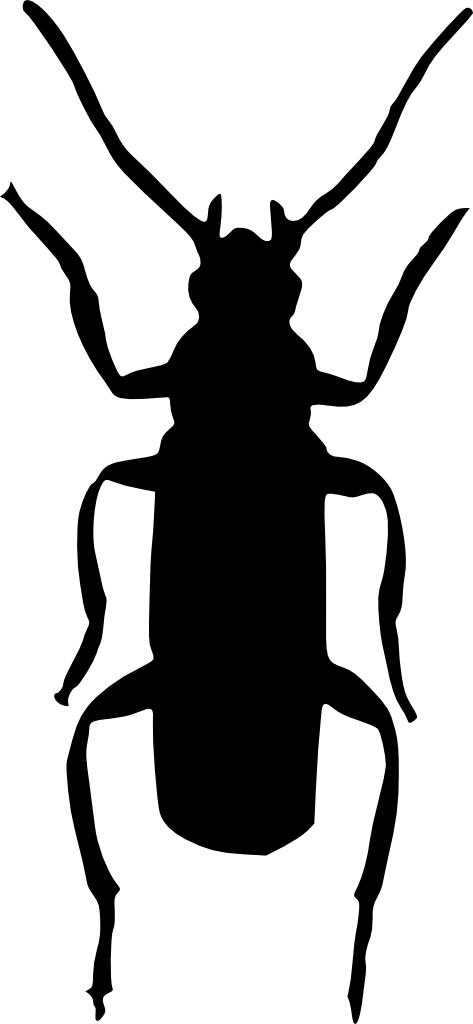

### mammal.png

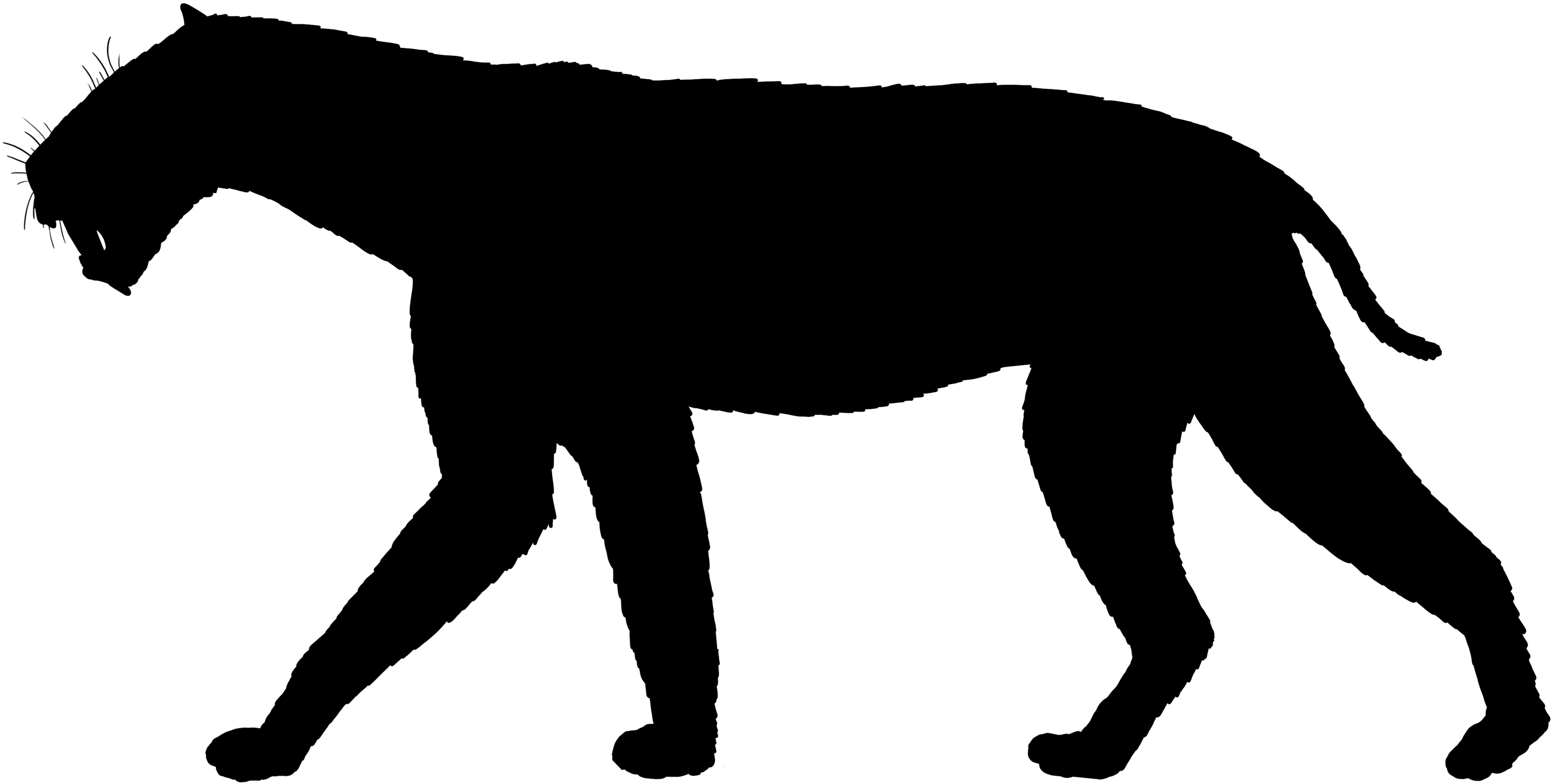

### plant.png

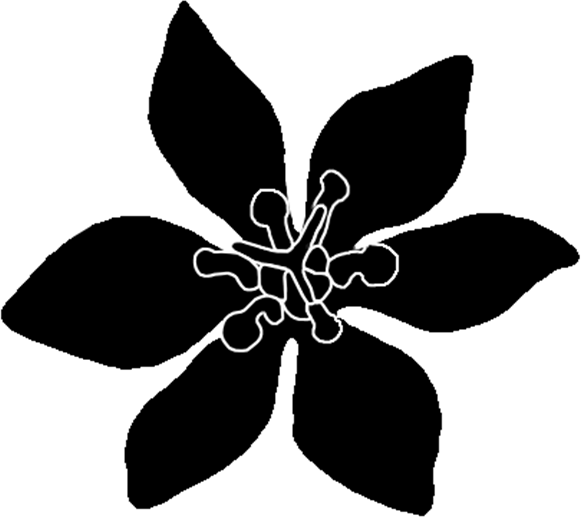

### reptile.png

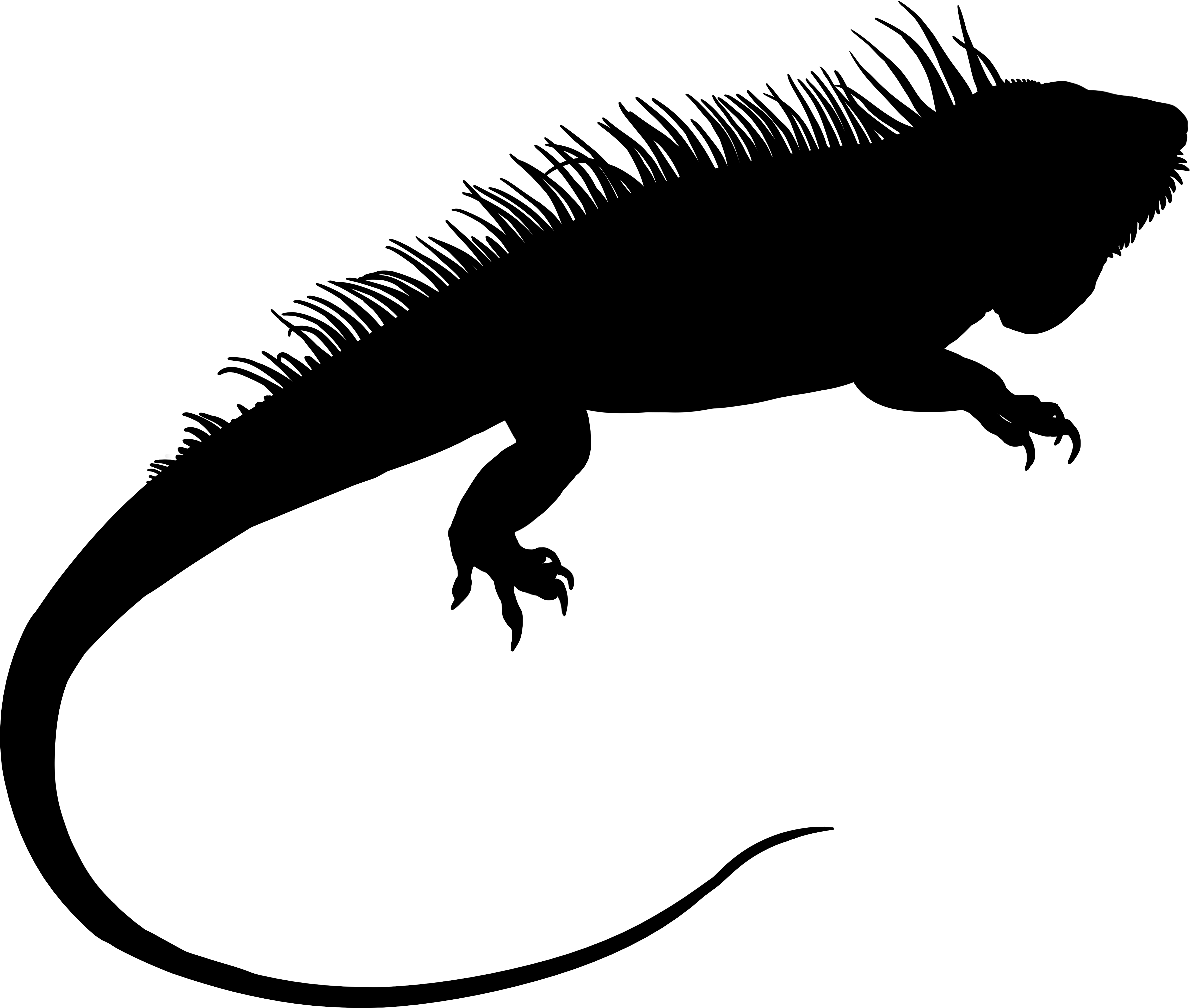
